# Development of microsatellite markers with the SSRseq method on *Glossina palpalis gambiensis*, and *G. p. palpalis*, and analysis of corresponding field samples from three sleeping sickness foci: Boffa, Dubreka (Guinea), and Bonon (Côte d’Ivoire)

**DOI:** 10.64898/2025.12.19.695359

**Authors:** Thierry de Meeûs, Sophie Ravel, Moïse Kagbadouno, Djakaridja Berté, Adeline Ségard, Olivier Lepais

## Abstract

Tsetse flies are strictly found is sub-Saharan Africa where they are responsible for the transmission and maintenance of African trypanosomiases in humans (HAT) and animals (AAT). Vector control has been recognized as an essential tool to fight against these diseases. Nevertheless, it requires the best possible knowledge of the biology of the targeted population. Population genetics tools can prove very useful to obtain such information but require the use of polymorphic and reliable genetic markers. In this paper we present the development of microsatellite markers using a new high-throughput sequencing based technology (SSRseq). We applied it on two species of tsetse flies from three HAT foci and obtained more accurate results as compared to microsatellite loci developed with classic methods. We could indeed use 9 to 14 SSRseq loci without the several problems generally met with classic microsatellites, as all were located in autosomes, without short allele dominance or stuttering and very few null alleles. SSRseq loci appeared much more polymorphic in tsetse flies as compared to other species (fungi, trees, bees, or fishes), which suggested much higher effective population sizes, much higher mutation rates of the genome or both, suggesting high capacities for evolutionary adaptation. With the 9-14 loci kept, we confirmed the propensity of these flies to move almost freely in the whole zones investigated, and also highlighted the possible evolutionary response of one of the unkept loci regarding vector control devices used in these HAT foci, which will require further studies. We suggest for further population structure studies to use only loci with less than 1% missing data, which proved being a good and fast selection strategy to get the most reliable results.

## Introduction

Tsetse flies (genus Glossina) are particular Diptera of the superfamily Hippoboscoidea, which are all blood sucking arthropods where, instead of laying eggs, a female lays full-sized pupae one at a time (Krafsur, 2009). Tsetse flies are strictly found is sub-Saharan Africa where they are responsible for the transmission and maintenance of pathogenic trypanosomes to humans and animals. In humans, the Human African Trypanosomiasis (HAT) (or sleeping sickness) has been eliminated as a public health problem in several countries and is targeted by the World Health Organization (WHO) for interruption of transmission (sustainable zero cases) by 2030 (Lejon et al., 2025). On the other side, Animal African Trypanosomiasis (AAT) (or nagana) is still representing a major burden to the economy of impacted countries, estimated in billions of dollars every year (Cecchi et al., 2024). Vector control has now been recognized as an essential tool to fight against these diseases, and has demonstrated its ability to protect hosts from infections by tsetse bites (Courtin et al., 2015), even when the impact on the population of vectors is moderate (Kagbadouno et al., 2024). The best of knowledge on the biology of a targeted population of vector can be key in setting the best control strategy and/or understanding the unsuccess of a campaign (De Meeûs et al., 2019). Fulfilling this goal can be achieved relatively easily by the use of population genetics tools applied to the study of the spatio-temporal variation of molecular markers. Indeed, such research can provide valuable information on reproductive strategies, population sizes and dispersal of targeted populations, which may be decisive in designing the best control strategies or in understanding the causes of failures of past campaigns (Solano et al., 2010). This however requires polymorphic genetic markers, with reliable genotyping profiles, in sufficient numbers to increase whole genome representativity as regard to the different signatures that are sought, neutrality regarding natural selection and appropriate sampling strategies (De Meeûs et al., 2007). Microsatellite markers still represent relatively cheap and efficient markers for non-model organisms such as ticks, sand flies or tsetse flies as is testified by recent publications (Prudhomme et al., 2020; Taraveau et al., 2024; Konan, et al., 2025; Dorsey et al., 2025)..

Tsetse flies, as non-model organisms, are challenging in this respect. Their whole genome is contained in very rare chromosomes, with a very large X chromosome that contains around 1/3^rd^ of microsatellite loci on average (Ravel et al., 2020), which restricts their use to only females, or reduces the number of useful markers to only 2/3rd of the few developed markers developed during each round of random libraries. Moreover, many of these markers suffer from stuttering, short allele dominance (SAD) and frequent null alleles. The algorithm developed by Chapuis and Estoup (Chapuis & Estoup, 2007) has proven very efficient at correcting for the effect of null alleles where they are not too frequent (Séré et al., 2017). Nevertheless, if fixing for stuttering and SAD is sometimes possible (De Meeûs et al., 2021), it is more challenging and even often impossible, leading to the withdrawal of affected loci.

Despite these issues and recent democratisation of SNPs technologies (which are not immune of genotyping imperfections), microsatellite loci still represent markers of choice for non-model organisms such as tsetse flies, as recently advocated in several papers (e.g.Layton et al. (2020); Lepais et al. (2020); De Meeûs & Noûs (2022)). SSRseq technology brings more arguments in favor of microsatellite markers for the study of the biology of natural populations given its ability to reduce homoplasy compared to capillary electrophoresis-based microsatellite genotyping (hereafter, “classic”) (Layton et al., 2020; Lepais et al., 2020, 2022).

SSRseq methodology (Lepais et al., 2020), uses high-throughput sequencing technologies to quickly develop and genotype sequence-based microsatellite markers. The method allows selecting location of loci, reject loci with stuttering or most unreliable ones (as for SAD) upstream. SSRseq thus theoretically allows avoiding most of the caveats described above with “old fashioned” microsatellite markers met in tsetse flies. Moreover, this technology offers ready to use datasets, the analysis of which does do not require specific genomics and/or bioinformatic skills, as it is the case for SNPs.

The aim of the present paper is to tranfer the SSRseq technology to the study of tsetse flies, vector of neglected tropical diseases, and which continue to affect subsaharan Africa, and present many issues regarding “old fashioned” species specific microsatellite markers. We describe the development of the same 26 SSRseq loci on two closely related taxa of tsetse flies in three HAT foci: *Glossina palpalis gambiensis* from Boffa and Dubreka in the Guinean Mangrove; and *G. p. palpalis* from Bonon in Côte d’Ivoire, which we consider separate species (De Meeûs et al., 2015), despite what the current nomenclature suggests. After quality testing and selection of the most reliable loci, we compare our results to those already published with old fashioned (hereafter labelled old) microsatellite markers (Berté et al., 2019; Kagbadouno et al., 2024). We finally conclude by providing some clues for the most efficient use of these new markers.

## Material and methods

### Ethical statement

A prior informed consent (PIC) was obtained from the local focal point and a mutually agreed terms (MAT) form was written and approved between Guinean and Côte d’Ivoire and French laboratories involved in the study for the use of the genetic diversity found in tsetse flies from these two countries.

### SSRseq development

The SSRseq loci were developed on the two closely related *G. p. gambiensis* (Gpg) and *G. p. palpalis* (Gpp).

#### DNA extraction, purification and library preparation

In Montpellier laboratory, 16 teneral (unfed) flies were used, four females and four males of Gpg and four females and four males of Gpp from the colonies reared at Cirad Montpellier were killed by freezing at -20°C just before DNA extraction. DNA was extracted from a mix of legs and head from the four females together and from the four males together of Gpg and of Gpp using DNeasy Blood and Tissue Kit (Qiagen, 157 Valencia, CA, USA) following the manufacturer’s instructions. In Bordeaux laboratory, DNA from these four samples (pool of four males and pool of four females from Gpg and from Gpp) were purified using 1.8 X Agencourt AMPure XP beads (Beckman Coulter, UK), quantified with a Qubit Fluorometer with the Qubit™ dsDNA BR assay kit and pooled in equimolar concentration. Whole genome DNA library were prepared using Qiagen QIASeq FX DNA library preparation kit and sequenced at a loading of 5 pM on an Illumina Miseq using half of a V2 flow cell in a 250 bp paired-end configuration.

#### Microsatellite loci detection, PCR conditions and sequencing

Paired-reads were merged using BBmerge v38.87 (Bushnell et al., 2017) and merged reads with a minimum of 150 bp were kept for microsatellite discovery using QDD software (Meglécz et al., 2010, 2014). Primer design parameters were set to target 100 bp to 180 bp amplicons and with primer parameters optimized for multiplex PCR (Lepais et al., 2020). To avoid selecting the numerous microsatellites located in the X chromosome, primer sequences were aligned to the reference genome of a female Gpg (accession number: GCA_000818775.1). We kept as suitable candidate loci only primer pairs that aligned to the reference and on the same contig at expected genomic distance from each other (no more than 250 bp). A total of 60 primer pairs were selected based on criteria maximizing amplification success (Meglécz et al., 2014, pure repeat instead of compound motifs, showing at least 20 bp between the primers and the repeat motif, and with flanking region showing high complexity (QDD cathegory A): no minisatellite, no other microsatellite in the flanking region) and polymorphisms (i.e. high number of repeats). Standard desalt oligonucleotides were ordered from Integrated DNA Technologies on a 96-well plate format at 25 nmol synthesis scale. We added the universal Illumina adapter sequences TCGTCGGCAGCGTCAGATGTGTATAAGAGACAG to the 5’ end of each forward primer and GTCTCGTGGGCTCGGAGATGTGTATAAGAGACAG at the 5’ end of each reverse primer. The DNA pool utilized for whole genome sequencing was used for PCR amplification of each primer pair. The simplex PCR was performed in a final volume of 10µL using Hot Firepol Blend master mix (Solis Biodyne), 10 ng of DNA and 0.2 µM of each primer. The PCR conditions consisted of an initial denaturation at 95°C for 15 min followed by 35 cycles of denaturation at 95°C for 20 s, annealing at 59°C for 60 s, extension at 72°C for 30 s, and a final extension step at 72°C for 10 min. Amplification was checked on a 3% agarose gel with a 100v migration for 25 min. Primer pairs that showed consistent amplification with amplicon at the expected size integrated the primer pool for the multiplex PCR. Multiplexed PCR amplification of the selected markers was performed in a final volume of 10µL using Hot Firepol Multiplex master mix (Solis Biodyne), 0.05 µM of each primer, and 20 ng of DNA. The PCR conditions consisted of an initial denaturation at 95°C for 12 min followed by 35 cycles of denaturation at 95°C for 30 s, annealing at 59°C for 180 s, extension at 72°C for 30 s, and a final extension step at 72°C for 10 min. The sequencing library was constructed using a second PCR that attached adapters and sample-specific pairs of indexes (8bp unique sequences, Nextera Indexing kit, https://www.illumina.com/products/by-type/accessory-products/nextera-dna-indexes.html) to each side of the amplicons by targeting the universal sequence attached to the locus specific primers. The indexing PCR is setup in a volume of 20 μL using Hot Firepol Multiplex master mix (Solis Biodyne), 5 µL of amplicon and 0.5 µM of each of the forward and reverse adapters. The PCR conditions consisted in an initial denaturation at 95°C for 12 min followed by 15 cycles of denaturation at 95°C for 30 s, annealing at 59°C for 90 s, extension at 72°C for 30 s, and a final extension step at 72°C for 10 min.

Markers validation was performed on a representative set of 95 blind repeated samples (48 Gpg from Boffa, Guinea and 47 Gpp from Bonon, Côte d’Ivoire) (see “Samples” section for more information on the samples from which these flies come from) and a negative control (containing water instead of DNA), and independently repeated from the multiplex PCR once on all samples in order to 1) optimize the bioinformatic pipeline to each locus, and 2) estimate locus-level allelic error rate (number of allele mismatches between replicates divided by the total number of alleles compared), and finally 3) select loci that produced repeatable genotypes for the final genotypic dataset (see next section “Bioinformatic pipeline for genotyping”).

Amplicons were pooled and purified with 1.8X Agencourt AMPure XP beads. Quality was checked on a Tapestation 4200 (Agilent) with a D1000 HS kit. Quantification was done using QIAseq Library Quant Assay kit (Qiagen, Hilden, Germany)) in accordance with the supplier’s protocol for Illumina using a Roche Light Cycler 480 quantitative PCR (95°C for 10 min followed by 35 cycles of denaturation at 95°C for 15 s, annealing at 60°C for 30 s, extension at 72°C for 120 s).. The pool was sequenced on an iSeq100 sequencer (Illumina, San Diego, CA, USA) with a 2×150 bp kit.

Short reads supporting the study has been submitted to European Nucleotid Archive (European Nucleotid Archive run accession number: ERR17897884) for shotgun whole genome sequencing used for microsatellite discovery and primer developments.

#### Bioinformatic pipeline for genotyping

The bioinformatics pipeline (Lepais et al., 2020) (https://doi.org/10.15454/HBXKVA) integrating the FDSTools analysis toolkit (Hoogenboom et al., 2017) was used to call genotypes (microhaplotypes) from raw sequences. All polymorphisms were integrated into micro-haplotypes including variation of the number of repeats, SNP and insertion-deletion (indel) in the microsatellite sequence and SNP and indel in its flanking sequences. The bioinformatics pipeline also compared genotypes from blind-repeated genotyping of the 95 samples described above plus 24 *G. p. palpalis* (see supplementary File S2, Sheet “Replicate_Identities”) to identify the loci that could not be reliably genotyped or for which the genotyping accuracy was only acceptable when analyzing the polymorphism within the repeat motif itself (i.e. not accounting for the polymorphism in the flanking sequence (Lepais et al., 2020)). For each locus, every allele differing from the other by any polymorphism type was coded under an arbitrary three-digit scheme with a unique number assigned to each microhaplotype per locus. A second dataset coding alleles according to allele size was also built, to compare the information content of sequence-aware coding scheme (micro-haplotype) and more conventional microsatellite dataset (i.e. as classically obtained by capillary-based electrophoresis).

Since amplification, sequencing and genotyping was repeated for each of these 110 individual, each fly was then characterised by two genotypes (labelled A and B in Table S1), for exactly 1/3 of gentyped individuals (Supplementary File S2, Sheet “Replicate_Identities”).

### Samples

The SSRseq loci developed (see Results section) were amplified and sequenced as described previously for the genotyping of all flies of the two different available sampled populations: *G. p. gambiensis* from the HAT foci of Boffa and Dubreka in Guinea; and *G. p. palpalis* from the focus of Bonon in Côte d’Ivoire. These sampled populations were already analysed with “old” microsatellite loci for Boffa (Kagbadouno et al., 2024) and Bonon (Berté et al., 2019), except for the 11 flies from Dubreka that happened to be available during the course of this study and were also analysed, which allowed extending the geographic range. As in other published results (Berté et al., 2019; Kagbadouno et al., 2024), we used a two months generation time. We have decided that C0 should always correspond to the generation before the beginning of the vector control campaigns (VCC) to which these populations have been submitted. All flies came from sleeping sickness foci: Boffa and Dubreka, in the Guinean mangrove (Gpg), and from Bonon (Gpp) (Côte d’Ivoire), covered with mesophilic forest, and plantations (cocoa and coffee). Available data consisted in 117 Gpg from Guinea (Boffa 2019 (cohort C50, 62 flies), Boffa 2020 (C56, 44 flies), and Dubreka 2020 (C56, 11 flies)), and 207 Gpp from the HAT focus of Bonon, Côte d’Ivoire (Berté et al., 2019) in June 2015 (C0, before control, 47 flies), in June 2016 (C6, 27 flies), in September 2016 (C8, 52 flies), in December 2016 (C9, 56 flies), and March 2017 (C10, 25 flies). These two data sets may be compared with published data on other old microsatellite loci (Berté et al., 2019; Kagbadouno et al., 2024), which we labelled “old loci”. The data are available in supplementary File S1 (with all loci characteristics and raw data) and S2 (final genotypes with all geographic and/or ecological information). Locations of samples can be visualized in Figure 1 and in published maps (Courtin et al., 2015; Berté et al., 2019; 2024; Kagbadouno et al., 2024). In Guinea, VCC followed different strategies according to the focus (Courtin et al., 2015; Camara et al., 2021). In Boffa, VCC began in 2012 in Eastern sites (around cohort C4) and was generalized in 2016 (C28). In Dubreka, it was deployed in 2016. More precisions can be found elswhere (Courtin et al., 2015; Camara et al., 2021; Kagbadouno et al., 2024). In Bonon, VCC began in 2016.

**Figure 1.**
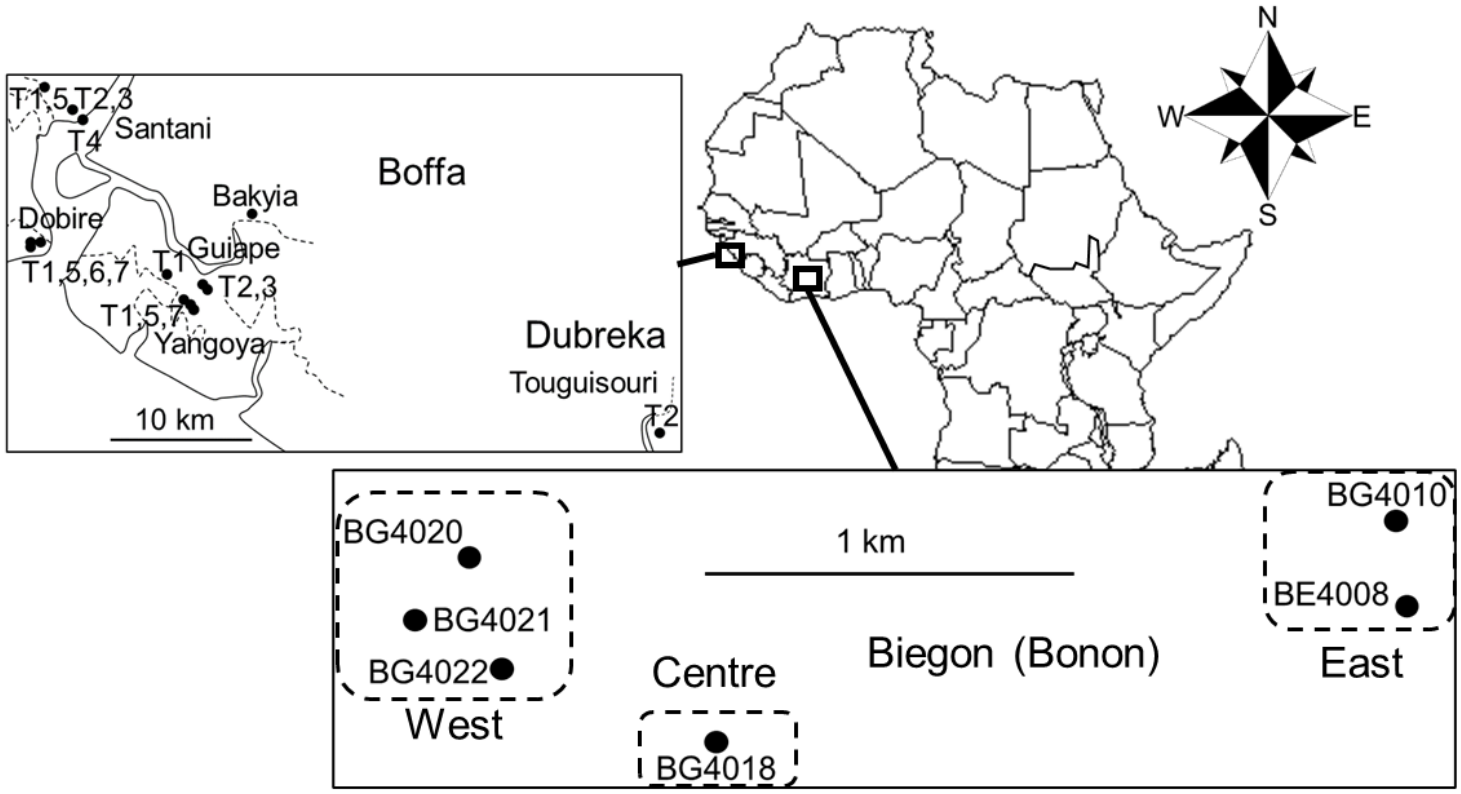
Sample locations of tsetse flies in different HAT foci: Boffa and Dubreka in Guinea, Bonon in Côte d’Ivoire.

### Population genetics analyses

#### Data set formatting

What we labelled raw data consisted in two genotypes per individual for the 95 flies used for the validation of the 26 SSRseq loci, plus 14 randomly chosen Gpp individuals (see Table S1). For each locus and each individual, the two genotypes differed in 10% of the time (from 80 to 95% across loci). When the two were heterozygotes, we picked the first one. When one genotype was homozygote and the other heterozygote, we kept the heterozygous profile. When one was missing, we kept the visible genotype. In the rarest cases where, at one locus, the two repeats were heterozygous with one allele in common, we kept the allele in common and kept the allele with the highest frequency in the sample for the second allele, or the first one (when equifrequent). This reinterpretation of data lead to what we called processed data (Supplementary Table S2) as opposed to what we called raw data (Supplementary Table S1)

Genotypes were available as alleles differing both for the number of motif repeats and sequences inside the amplified fragment (SNPs and indels). We first analyzed the data set considering all polymorphisms. We also analyzed a dataset where only variations in microsatellite repeats were considered (allele size).

Data were then coded in Create software format (Coombs et al., 2008) and converted into appropriate formats when needed, for each of the two species, and for the different kind of analyses performed.

All analyses were undertaken on two datasets of the same individual tsetse flies. The first dataset considered all allele kinds, i.e. alleles with different repeat numbers of the microsatellite motif, SNPs, insertions and deletions (indels). The second data set did not consider SNPs, but only change on allele length instead (microsatellite repeat number variation in addition to eventual indels). The first was labelled dataset with all polymorphisms, and the second dataset with allele size polymorphism.

#### Wright’s fixation indices

In a three-level hierarchy (individuals, subsamples and total), Wright’s *F*-statistics (Wright, 1965) are indices that measure the relative contribution to inbreeding of a given hierarchical level as compared to another level, upper in the hierarchy: *F*_IS_ is the contribution of individuals relative to subsamples, *F*_ST_ is the contribution of subsamples relative to the total sample, and *F*_IT_ is the contribution of individuals relative to the total. More specifically, *F*_IS_ measures the effects of deviations from local panmixia, *F*_ST_ measures the effect of subdivision and *F*_IT_ results from the two previous ones. These statistics were estimated with Weir and Cockerham’s unbiased *f* (*F*_IS_), *θ* (*F*_ST_) and *F* (*F*_IT_) (Weir & Cockerham, 1984). The significant deviation from 0 of these statistics was undertook with 10,000 randomizations of alleles between individuals within subsamples to test for local panmixia and of individuals between subsamples to test for significant subdivision. For panmixia, the statistics used was *f*, while we used the logarithm of the likelihood ratio *G* for subdivision (Goudet et al., 1996). For panmixia, tests were two-sided as heterozygote deficits or excesses could be expected under different alternative hypotheses (H1). Indeed, heterozygote excesses are expected in dioecious organisms, in random mating populations (De Meeûs & Noûs, 2023), as it is the case here, and heterozygote deficits can be met for many reasons as systematic sib-mating, Wahlund effects or amplification problems (De Meeûs et al., 2007). The two-sided *p*-values were obtained by doubling the *p*-value if it was below 0.5, or doubling 1-*p*-value otherwise. For subdivision, tests were one-sided (observed *G* exceeds the expected one under the null hypothesis, H0). All these were computed with Fstat 2.9.4 (Goudet, 2003), updated from the published version (Goudet, 1995). More details can be found in another paper (De Meeûs et al., 2007). In some instances and some loci, we tested if subdivision was smaller than expected by chance (homogenizing selection). In case of homogenizing selection, the estimator of *F*_ST_ is expected to display negative values (*θ*<0). To test if *F*_ST_ was significantly negative, we used the one sample and one-sided (H1: *θ*<0) Wicoxon signed rank test on the distribution of *F*_ST_ estimated per allele provided by Fstat analysis. For this test we used the package R-commander (Rcmdr) (Fox, 2005, 2007) for R (R-Core-Team, 2025).

The smallest geographic scale was the trap. We undertook the first analyses at that scale. Other hierarchical (i.e. nested) levels were the village or site, the focus (in Guinea), the cohort and the total sample.

#### Detection of significant levels of subdivision by the Wahlund effect procedure

We tested at which level population genetic subdivision occurred with the *F*_IS_ based method (Goudet et al., 1994). We used multiple comparisons with a Wilcoxon signed rank test for data paired by locus, with alternative hypotheses: *F*_IS-traps_<*F*_IS-villages_ or *F*_IS-villages_<*F*_IS-focus_<*F*_IS-cohort_. These tests were undertaken with Rcmdr). We also measured and tested the significance of subdivision at each of these levels. Comparisons of *F*_ST_’s followed the same pattern, except that the alternative hypothesis was in the other direction (Wahlund effect tends to decrease *F*_ST_ between remaining subsamples (De Meeûs, 2018)). We finally compared the different sample designs with the number of locus pairs that were found in significant linkage disequilibrium (LD). Linkage disequilibrium was tested between each pair of loci with the multi-sample *G*-based test with Fstat, and 10,000 randomizations. It is the most powerful procedure when used across different subsamples (De Meeûs et al., 2009). The number of significant LD tests is expected to increase in case of Wahlund effects of moderate amplitude (Manangwa et al., 2019). The comparisons of the proportion of significant LD tests were undertaken with the Fisher exact tests with R (command fisher.test) with a one-sided alternative hypothesis (H1: the proportion of significant tests increases when pooling subunits).

All paired tests produced series of *p*-values with some degree of dependency. As expounded in Appendix 1, we used Benjamini and Hochberg correction (BH) (Benjamini & Hochberg, 2000) for all multiple testing situations in this paper.

#### Quality testing of the different loci

We first checked the statistical independence of loci with the G-based test for LD across subsamples. There are as many non-independent tests as there are locus pairs (here 325 pairs). These *p*-values were adjusted with the BH procedure.

We also computed standard error of *F*_IS_ and *F*_ST_ from jackknives over loci, SE_FIS and SE_FST, to be used for null allele detection (De Meeûs, 2018): in case of null alleles, the ratio SE_FIS/SE_FST=*r*_SE_>2. We also computed 95% confidence intervals (95%CI) with 5,000 bootstraps over loci. All these statistics were computed with Fstat. We also utilized Genetix (Belkhir et al., 2004) to compute 95%CI of *F*_IS_ with 5,000 bootstraps over individuals, for each locus, in each subsample. We computed the averages across subsamples of these 95%CI for each locus. Genetic diversity and subsamples sizes could vary substantially across subsamples. We thus weighted these averages with subsample sizes (number of observed genotypes) and the corresponding unbiased local genetic diversities (Nei’s *H*_S_) (Nei & Chesser, 1983), computed by Fstat.

In case of significant heterozygote deficit, we first looked for the existence of null alleles. We estimated *F*_IS_/*F*_ST_ ratio of standard errors (SE) of jackknives over loci (*r*_SE_). Additionally, we computed the correlation between *F*_IS_ and *F*_ST_ and between *N*_blanks_ (number of missing genotypes) and *F*_IS_ with the Spearman’s rank correlation test. The graphic of the regression *F*_IS_∼*N*_blanks_ helped us determining loci for which missing genotypes actually correspond to null homozygotes, and those for which other problems occurred. We estimated the frequencies of null alleles for each locus with the EM algorithm (Dempster et al., 1977) with the software FreeNA (Chapuis & Estoup, 2007). For this, and unless specified otherwise, we recoded missing genotypes as homozygous for allele 999, except for loci for which missing data occurred for other reasons, as recommended. With these, we computed the average (weighted for subsample sizes) of null allele frequencies *p*_nulls_ to be used for the graphic of the regression *F*_IS_∼*p*_nulls_ across loci and the averaged 95%CI of bootstraps over individuals described above. If some loci, coded 999999 for missing genotypes, displayed points below the slope, we recoded missing data as “0” for these loci and re-ran FreeNA and undertook a new regression. This process was repeated as many times as necessary, until an optimal result was obtained (highest *R*^2^ and smallest intercept). We then compared the observed number of missing genotypes at each locus to the expected one (i.e. *N*_miss-exp_=∑*p*_nulls-*s*_^2^×*N*_*s*_, where *p*_nulls-*s*_ and *N*_*s*_ are the frequency of null alleles and the size of subsample *s*, respectively), for each locus, with a one-sided exact binomial test with R (command binom.test) (H1: there are not enough missing genotypes observed in the data as compared to the number of null homozygotes expected). All loci that significantly did not exhibit enough missing data as compared to the expected value (*N*_miss-exp_) were excluded from the data. The regression *F*_IS_∼*p*_nulls_ also allowed computing the proportion of the variance of *F*_IS_ explained by null alleles (*R*^2^) and the value of *F*_IS_ predicted when there is no null alleles (intercept: *F*_IS-0_) and its 95%CI.

In case of doubt, to check the neutrality of loci, we computed the raw standard error of *F*_ST_ over alleles (SEalleles(*F*_ST_)) and tested its correlation with *F*_ST_ across loci. If significantly positive (Spearman with Rcmdr), we progressively removed loci with the highest SEalleles(*F*_ST_), and retested the correlation with the remaining loci, until it became not significant. This provided the threshold SEalleles(*F*_ST_) above which loci were suspected to be under selection.

We also computed the number of times each locus displayed a significant LD with another locus (NLDSig) and the average across loci (AvNLDSig) and its 95%CI computed with the standard error of the mean.

We also excluded all loci with a frequency of null alleles *p*_nulls_>0.3 (Séré et al., 2017). After that, we likewise excluded all loci with at least two of the following criteria: SEalleles(*F*_ST_)>MaxSE, *F*_ST_ not in the 95%CI of bootstraps over loci and a significant difference from 0, and NLDSig greater than the upper limit of the 95%CI. We also plotted the evolution of allele frequencies across cohorts to check if the evolution of allele frequency responded to VCC or to something else (supplementary file S3).

Finally, for allele size datasets, we tested for stuttering and SAD for loci not explained by null alleles. Stuttering was tested with the spreadsheet method (De Meeûs & Noûs, 2022). We then tested for the influence of SAD with the regression *F*_IS-i_∼*A*_*i*_ weighted by *p*_*i*_(1-*p*_*i*_), where the subscript *i* means allele of size *A*_*i*_ and *p*_*i*_ the corresponding allele frequency (De Meeûs et al., 2004). These were undertaken with Rcmdr. For the regression, Rcmdr provides two-sided *p*-values. One-sided *p*-values (H1: the slope is negative) were computed by hand as *p*-value/2 (negative slope) or 1-*p*-value/2.

#### Subdivision measures and testing

For ordinal data that corresponded to *n* matched subsample groups (i.e. cohorts), we compared different statistics (e.g. *H*_S_) between these groups (cohorts) with the Friedman two-way analysis of variance with Rcmdr, with matching units being the different loci. If significant, we compared paired cohorts with the Wilcoxon signed rank test for paired data with rcmdr and adjusted *p*-values with the BH procedure.

We used the recoded data as described above for FreeNA to compute *F*_ST-ENA_ corrected for null alleles with the ENA algorithm (Chapuis & Estoup, 2007), with 5,000 bootstraps over loci to compute 95%CI.

Microsatellite markers are so polymorphic that the maximum *F*_ST_<1, meaning its value both reflects population size, migration and mutation. We thus estimated a standardized version of *F*_ST_ independent of mutation rates. Two standardizations are available: the *F*_ST_’ and the *G*_ST_” based methods. The first is computed as *F*_ST_’=*F*_ST_/*F*_ST-max_, where *F*_ST-max_ is the maximum *F*_ST_ when all subsamples, with the same degree of polymorphism of raw data (same *H*_*s*_), harbor no allele in common (Hedrick, 2005b; Meirmans, 2006). Alternatively, *G*_ST_’’=*n*(*H*_T_-*H*_S_)/[(*nH*_T_-*H*_S_)(1-*H*_S_) (Meirmans & Hedrick, 2011) is computed with the number of subsamples (*n*), Nei’s unbiased estimators of local and total genetic diversities (*H*_S_ and *H*_T_, respectively). Nevertheless, no known software allows computing 95%CI of *G*_ST_”, and in case of null alleles, no known procedure allows correcting *G*_ST_” for those. Moreover, as *n* increases, *G*_ST_”→*G*_ST_/*G*_ST-max_≈*F*_ST_’. This is why we systematically computed *F*_ST_’. Maximum possible subdivision index *F*_ST-max_ was computed with a dataset recoded by RecodeData (Meirmans, 2006), from the initial Fstat dataset. We thus could compute *F*_ST-ENA_’=*F*_ST-ENA_/*F*_ST-max_ to obtain a subdivision measure corrected for null alleles and polymorphism, and its 95%CI. Due to the impossibility to handle missing data appropriately with RecodeData, we decided to not correct *F*_ST-max_ for null alleles. This will, at worst, provide a slight over estimation of the maximum possible *F*_ST_ and hence a slight underestimate of the standardized *F*_ST-ENA_’.

#### Effective population sizes

Effective population size (*N*_*e*_) was estimated with the LD method with correction for missing data (Waples, 2006; Waples & Do, 2010; Peel et al., 2013), the coancestry method (Nomura, 2008) (CoA), with NeEstimator (Do et al., 2014), the intra and inter loci correlations (Vitalis & Couvet, 2001a) with Estim 1.2 (Vitalis, 2002) updated from (Vitalis & Couvet, 2001b) (1L2L), the heterozygote excess method (De Meeûs & Noûs, 2023) (HEx), and the sibship frequency method (Wang, 2009), computed with Colony (Jones & Wang, 2010) (Sib). For LD, we assumed random mating, and set the minimum allele frequency for alleles at 0.05 as recommended in NeEstimator documentation. Values were averaged across subsamples, and the grand average was computed across methods, with the number of usable values (other than infinity) as weights as in other papers (De Meeûs & Noûs, 2023). When appropriate, we also used the three temporal methods available in NeEstimator (Nei & Tajima, 1981; Pollak, 1983; Jorde & Ryman, 2007), and the two methods computed in mne (Wang & Whitlock, 2003): maximum likelihood (ML) and moment methods. For all methods we ignored “Infinite” or negative values (equivalent to “Infinite”) and computed weighted arithmetic means of *N*_*e*_ and of minimum and maximum values obtained across single sample and across temporal methods, as described in De Meeûs and Noûs (De Meeûs & Noûs, 2023). Confidence intervals may be computed but are useless here as most of the time these did not contain the average, due to the many “Infinite” results. According to Waples, ignoring “Infinite” results necessarily leads to underestimate *N*_*e*_ (Waples, 2024). Alternatively, he proposed to set such values at 999999 and compute the harmonic means. We thus followed this advice, setting “Infinite” values as the maximum value we found across all methods and rounded to the closest upper figure with zeros (e.g. 56 gives 100). We then used 95%CI as outputted in the different procedures (jackknives for CoA, bootstraps for Hex, or parametric for other procedures).

#### Dispersal

In case of significant subdivision, we computed *F*_ST-ENA_’between each pair of subsample.. We also calculated the corresponding geographic distances *D*_geo_ with the GPS coordinates of each subsample and the package geosphere (Hijmans et al., 2019) for R (see Appendix 3). We then estimated the number of immigrants in each subpopulation, assuming an Island model of migration, as (see Appendix 2):

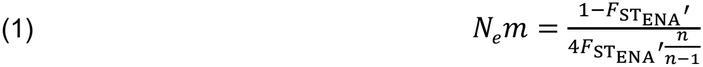

where *N*_*e*_ is the effective subpopulation size, *m* is the immigration rate and *n* is the number of subpopulations (Islands) in the total population (assumed infinite if unknown). We computed average distances between paired subsamples av(*D*_geo_).

These averages could be used to compute immigration rates between each pair of subsamples as *m*=*N*_*e*_*m*/*N*_*e*_. When *F*_ST-ENA_’≤0, we set *m* as the maximum possible one *m*_max_=1-1/*n* (assumed = 1 if *n* unknown). Dispersal distances per generation could then be computed as *δ*=*m*×av(*D*_geo_). In case of absence of subdivision signature, we assumed free dispersal across the whole range of the sampled area: *δ*=*D*_geo-max_, where *D*_geo-max_ is the distance between the two most remote traps of the study where tsetse flies were genotyped.

When applicable, we studied isolation by distance through Rousset’s model in two dimensions (Rousset, 1997): *F*_R_=*c*+*b*×ln(*D*_geo_), where *F*_R_ is *F*_ST_ENA_/(1-*F*_ST_ENA_) between each subsample pairs, as recommended in case of null alleles (Séré et al., 2017), *c* is the intercept, *b* the slope of the regression and ln(*D*_geo_) is the natural logarithm of the geographic distance between subsamples. Confidence intervals were computed with FreeNA after 5,000 bootstraps over loci, on a dataset recoded as described above. If the relationship is significant (all slopes>0), then the reverse of the slope describes the neighborhood (*Nb*) (effective number of connected individuals) of the population under investigation, and 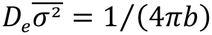. In this formula, *D*_*e*_ is the effective population density, and 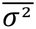 is the average of the squared axial distance between adults and their parents (Rousset, 1997). Axial distance is half the distance between adults and their parents (*δ*), and if the effective population size of the population (*N*_*e*_) and the surface occupied by concerned individuals (*S*) are known, then *D*_*e*_=*N*_*e*_/*S* and 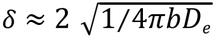 (De Meeûs et al., 2019).

Some distances between specific points were approximated with the tool “Draw a path” in Google Earth Pro. We also used the button “Draw a polygon” to obtain surface measures of concerned areas. Under Google Earth Pro, each trap can be located by clikcing on the button (yellow pin) “Add Placemark” and filling the information frame with the GPS coordinates (Supplementary File S2). Drawing a path or a polygon with the corresponding buttons is then straightforward.

## Results and Discussion

### SSRseq development

On the 60 developed primer pairs, only 2 failed to produce specific amplification, thus 58 loci were kept in the multiplexed PCR (Supplementary file S1). Among the 58 loci, 32 produced unreliable genotypes due to low amplification and sequencing, in addition to unspecific amplification and strong stuttering, that hampered accurate genotyping (Supplementary File S1). Notably, di-nucleotide repeats microsatellites showed low development success compared to tri- and tetra-nucleotide repeats microsatellites (respectively 16%, 80% and 82%, Table 1). This pattern is likely caused by high level of nucleotide polymorphism of the species leading to numerous substitutions in noncoding genomic region where dinucleotides microsatellite loci are more numerous. Mismatches in primer hybridization zones can cause suboptimal amplification efficiency and null alleles for dinucleotide microsatellites. Alternatively, tri- and tetra-nucleotide microsatellite loci are more frequent in genic and intergenic region and are thus more likely found in less polymorphic part of the genome. This probably why tri- and tetra-nucleotide loci showed more consistent amplification and sequencing, producing more reliable microsatellite compared to dinucleotide microsatellites. The 26 remaining microsatellite producing repeatable genotypes identified a total of 669 alleles differing in sequence (mean: 25.73 alleles per loci) and only 320 allele differencing by size (mean: 12.31 alleles per loci) demonstrating high size homoplasy for the species (109% additional allele revealed by sequence information). Dinucleotide microsatellites showed higher size homoplasy compared to tri- and tetra-nucleotide repeats, corroborating the hypothesis that they are located in more polymorphic part of the genome. Therefore, they are more subject to missing data due to less efficient amplification, which might translate into higher occurrence of null alleles for this class of microsatellite. In addition, they are more difficult to accurately genotype as showed by the higher allelic error rate which might be due to lower coverage but also to more complex variability and stuttering caused by higher instability of dinucleotide repeats as shown by the higher number of alleles observed for this class of microsatellite loci (Table 1).

**Table 1.** Development success compared among SSR motif types.

| **SSR in the multiplex PCR** |  |  |  |  |  |
|---|---|---|---|---|---|
| **Motif** | **N** | **Xc** |  |  |  |
| 2 | 32 | 97 |  |  |  |
| 3 | 15 | 211 |  |  |  |
| 4 | 11 | 154 |  |  |  |
| **Unreliable SSR** |  |  |  |  |  |
| **Motif** | **N** | **%** | **Xc** |  |  |
| 2 | 27 | 84% | 86 |  |  |
| 3 | 3 | 20% | 69 |  |  |
| 4 | 2 | 18% | 23 |  |  |
| **SSR producing reproductible genotypes** |  |  |  |  |  |
| **Motif** | **N** | **%** | **Xc** |  |  |
| 2 | 5 | 16% | 157 |  |  |
| 3 | 12 | 80% | 246 |  |  |
| 4 | 9 | 82% | 183 |  |  |
| **Raw genotypic dataset characteristics** |  |  |  |  |  |
| **Motif** | **AllSeq** | **AllSiz** | **Homop** | **Miss** | **Error** |
| 2 | 32 | 13.8 | 132% | 7% | 2.70% |
| 3 | 24.8 | 12.9 | 92% | 2% | 0.30% |
| 4 | 23.4 | 10.7 | 119% | 2% | 1.00% |

Motif: 2: di-nucleotide; 3: tri-nucleotide; 4: tetra-nucleotide; N: number of SSR, Xc: mean sequence coverage per sample and SSR, %: percentage of SSR; AllSeq: mean number of alleles based on sequence per loci; AllSiz : mean number of alleles based on size per loci ; Homop: percentage of additional allele revealed by sequence information over number of allele differing by size; Miss: percentage of missing genotypes; Error: percentage of allele mismatch identified between replicated genotypes over total number of compared replicated genotypes.

### SSRseq loci in Gpg from the Boffa and Dubreka foci in Guinea

#### Dataset formatting

Results of this procedure can be checked in the Supplementary File S2. In this file, we can see that among the replicated individual samples, 96.68% replicated genotypings provided an identical genotype (averaged across loci and individuals) in minimax=[87.27, 100] across loci or minimax= [79.92, 100] across individuals.

#### All polymorphisms

##### Detection of significant levels of subdivision by the Wahlund effect procedure

The results of sampling design comparisons are presented in the Figure 2. It can be seen that there is in fact no geographic structure, and no temporal structure, as already shown with old loci (Kagbadouno et al., 2024). For LD tests, no subsequent sampling designs displayed significantly different amounts of significant tests, but the significant difference observed, say, between traps and cohorts can be attributed to increase in power resulting from bigger subsample sizes (Figure 2).

**Figure 2.**
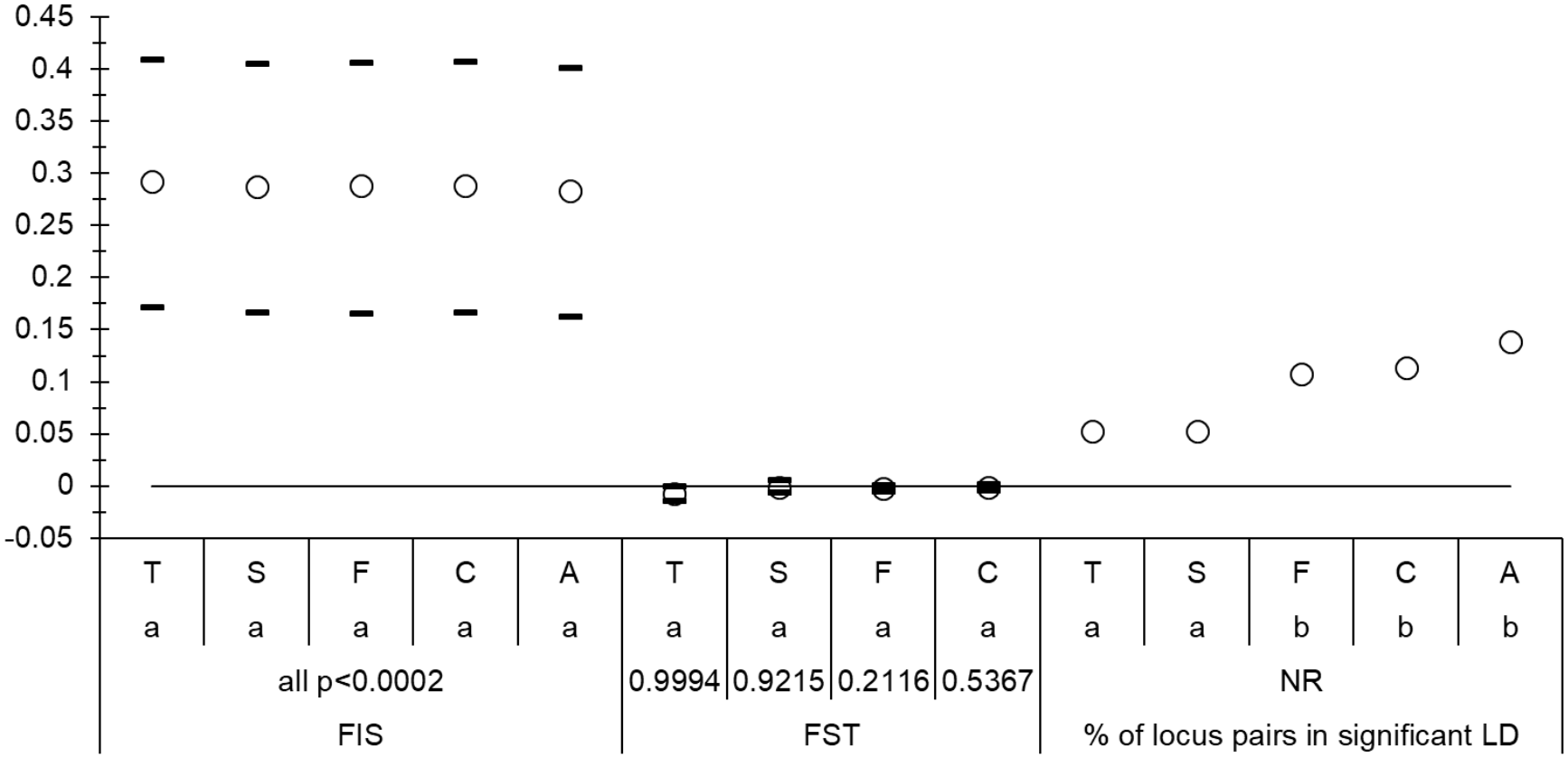
Values of *F*_IS_ (left), *F*_ST_ (center) and proportion of locus pairs in significant linkage disequilibrium (LD) (right for the SSRseq loci with all polymorphisms for *Glossina palpalis gambiensis* from Boffa and Dubreka (Guinea). Average values (empty circles) are surrounded by the 95% confidence intervals (black dashes) from bootstraps over loci (only for *F*_IS_ and *F*_ST_). Comparisons of different sampling designs regarding *F*_IS_ and *F*_ST_ (paired comparisons) and the proportions of locus pairs in significant linkage (paired Fisher’s exact tests) are presented. T, S, F, C, and A correspond to the sampling designs: Traps, Sites, Focus, Cohorts, and All in one sample respectively. Small captions (a, b) correspond to groups of non-significant paired tests (sharing one letter). Paired tests were adjusted for repetitions with the BH procedure. Significant deviation from 0 are also provided for *F*_IS_ and *F*_ST_ (not relevant for LF: NR).

Taking into account the aforementioned results, and unless mentioned otherwise, we considered the cohort as the relevant subsample unit for subsequent analyses, and thus ignored geographic information.

##### Quality testing of the different loci

The number of locus pairs in significant LD (38, 12%) was relatively high, with an average AvNLDSig=3 per locus and a 95%CI=[2, 4].

We observed a highly significant and variable *F*_IS_=0.287 in 95%CI=[0.165, 0.406] *(p*-value<0.0002). The ratio *r*_SE_=31.5, was far above 2, which suggests an important proportion of null alleles. This was not confirmed by the correlation between *F*_IS_ and *F*_ST_, which was negative. This would mean that null alleles have a poor effect on *F*_ST_, which was very small and not significant anyway (*F*_ST_=-0.001 in 95%CI=[-0.004, 0.002], *p*-value=0.5367). Neither *F*_IS_ or *H*_S_ differed significantly between the two cohorts (*p*-values>0.18).

We noticed a strong and highly significant correlation between *F*_ST_ and SEAlleles(*F*_ST_) (*ρ*=0.5364, *p*-value=0.0024). The correlation stayed significant as long as we kept loci with SEAlleles(*F*_ST_)>0.0025 (*ρ*=0.2711, *p*-value=0.1549).

The correlation between *N*_blanks_ and *F*_IS_ was positive though not significant, and the corresponding regression (Figure 3) allowed identifying four loci (L027-Tri, L028-Di, L032-Di and L053-Tetra) that displayed too many missing data. For these four loci, missing data were thus not coded as homozygous for null alleles (i.e. 999999) for the estimation of null allele frequencies with FreeNA.

**Figure 3.**
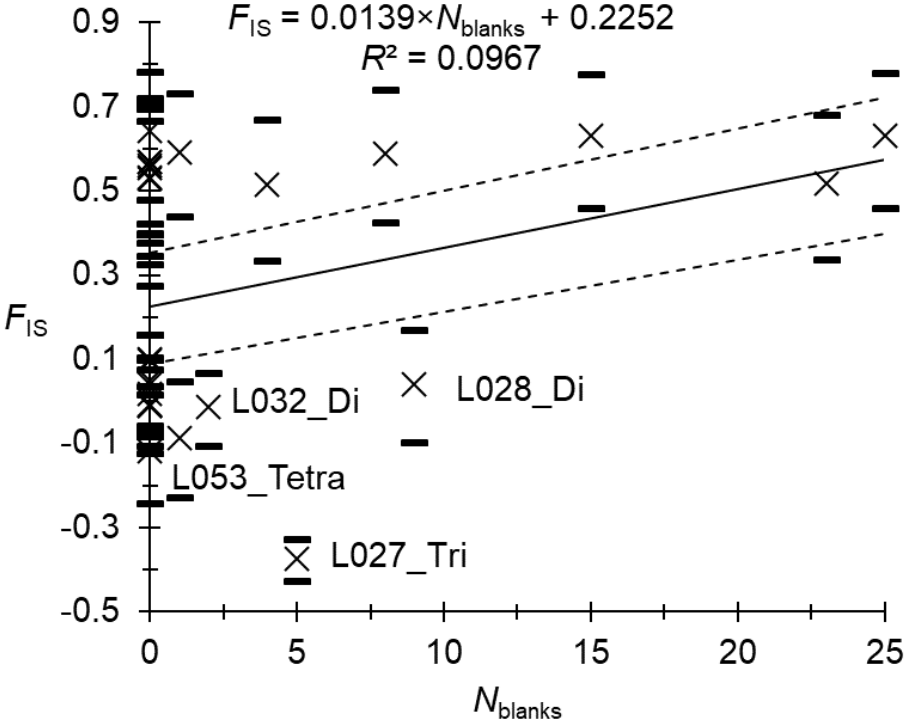
Regression of *F*_IS_ as a function of the number of missing genotypes (*N*_blanks_) at the different SSRseq loci (all polymorphisms) for *Glossina palpalis gambiensis* from the foci of Boffa and Dubreka (Guinea), computed within each cohort (C50 and C56). Regression of averages is represented with a black line, and 95% confidence intervals (5,000 bootstraps over individuals) with dotted lines. Each cross surrounded by two dashes correspond to each locus and its 95%CI. Loci with too many blanks (with significantly negative residuals) are pointed with their names. The regression model and its determination coefficient are presented.

After several attempts of the regression *F*_IS_∼*p*_nulls_, we decided to recode missing data as 999999 only for loci L045-Tri and L047-Tetra. The graphic of the regression *F*_IS_∼*p*_nulls_ is presented in the Figure 4. Following the criteria expounded above, we removed 12 loci from this regression. More information for each locus can be checked in the supplementary file S3. With the remaining 14 loci, the regression displayed high performances, with a *R*^2^=0.9794 and an intercept *F*_IS-0_=-0.0177 in 95%CI=[-0.1321, 0.0902], very close to the value obtained with old autosomal microsatellite markers (*F*_IS-0_=-0.0381 in 95%CI=[-0.1794, 0.0936]) (Kagbadouno et al., 2024), but with a narrower confidence interval. These intercepts would represent the *F*_IS_ that would be measured on average with loci free of null alleles in a pangamic dioecious population with effective population size (De Meeûs & Noûs, 2023): *N*_*e*_=26 in 95%CI=[3, Infinite].

**Figure 4.**
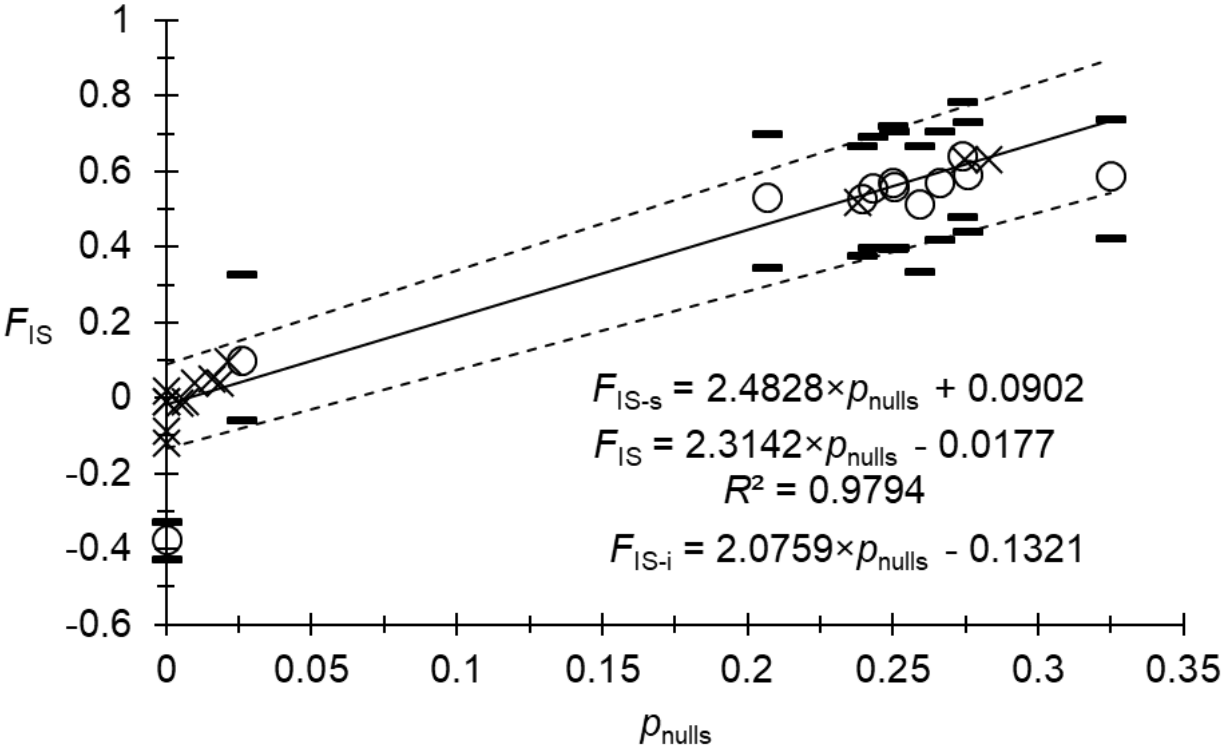
Regression of *F*_IS_ as a function of null allele frequencies (*p*_nulls_) (weighted averages across cohorts) at the different SSRseq loci (all polymorphisms) for *Glossina palpalis gambiensis* from the foci of Boffa and Dubreka (Guinea), computed within each cohort (C50 and C56). Regression of averages (black crosses) is represented with a black line, and of 95% confidence intervals (5,000 bootstraps over individuals) with dotted lines. Equations (with slope and intercept), for average and confidence intervals, and determination coefficient (*R*^2^), for average, are also indicated. Problematic loci, as defined in the text (quality testing paragraph), are pointed with empty circles and their 95%CI (black dashes) and were excluded from the regression computations.

We then reanalyzed this new dataset with the remaining 14 loci in each cohort. We observed a decrease of the number of significant LD tests to 3 (3%), none of which remained significant after BY correction (all *p*_BH_>0.7).

##### Subdivision measures and testing

The correspondence between *F*_ST_’ and *G*_ST_” was very tight (slope *b*=0.9689, intercept *a*=0.0013, *R*^2^=0.9835). Between cohorts, we measured a non-significant *F*_ST-ENA_’=0.0022 in 95%CI=[-0.0113, 0.0171] (*p*-value=0.3048). This is a comparable result as the one obtained with old loci, which nevertheless provided much smaller values and wider 95%CI (*F*_ST-ENA_’=-0.0161 in 95%CI)[-0.0246, -0.0057] (*p*-value=0.9889).

#### Allele size polymorphism

##### Detection of significant levels of subdivision by the Wahlund effect procedure

The comparison of results obtained between sampling designs were similar to those with all polymorphism (Figure 5). Unless specified otherwise, we kept cohorts as subsample units for the analyses to come.

**Figure 5.**
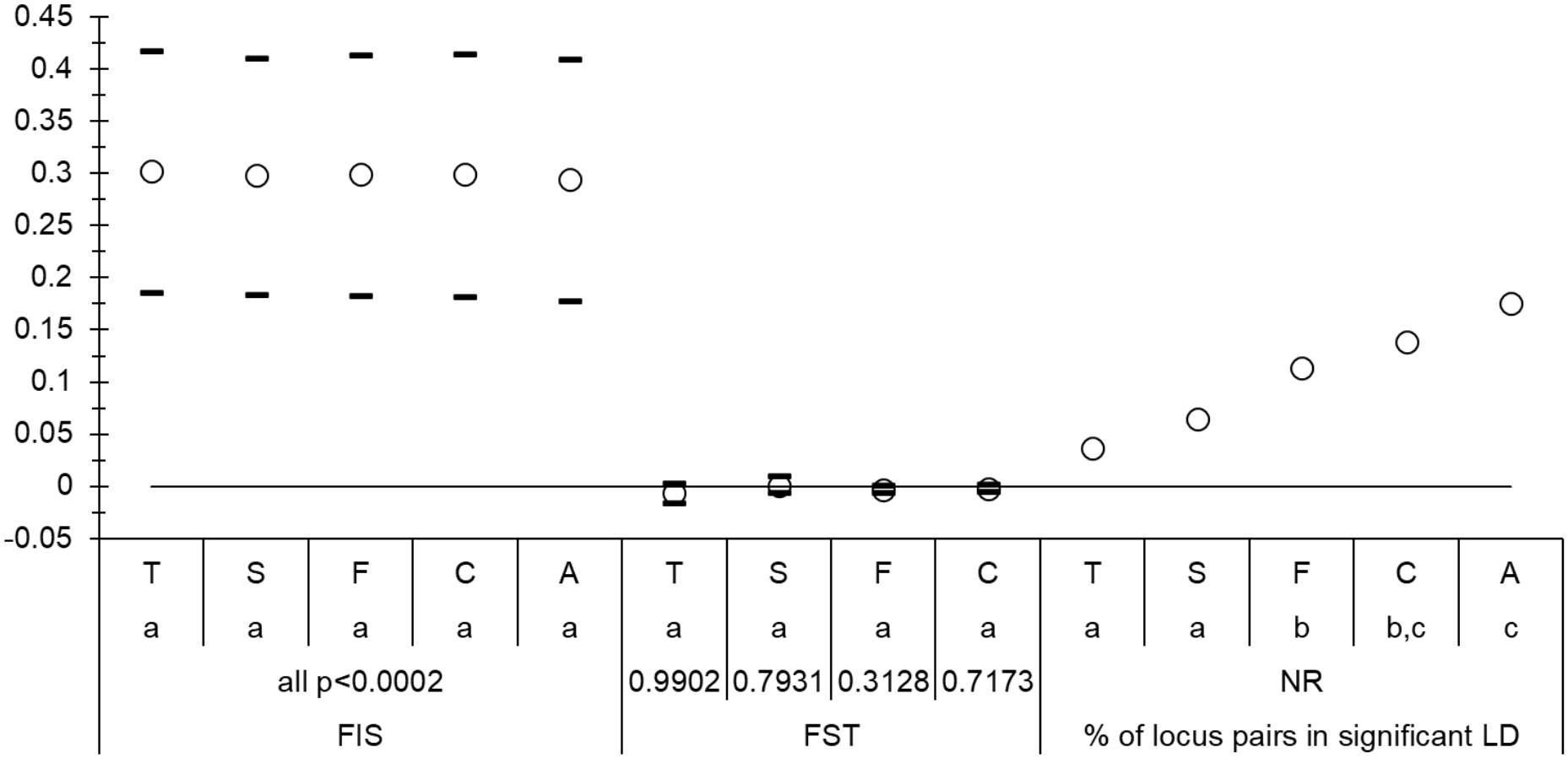
SSseq loci with allele size polymorphism for *Glossina palpalis gambiensis* from Boffa and Dubreka (Guinea). Comparisons of different sampling designs regarding *F*_IS_, *F*_ST_ and the proportions of locus pairs in significant linkage (paired Fisher’s exact tests). T, S, F, C, and A correspond to the sampling designs: Traps, Sites, Focus, Cohorts, and All in one sample respectively. Small captions (a, b,c) correspond to groups of non-significant paired tests (sharing one letter). Paired tests were all one-sided and adjusted for repetitions with the BY procedure.

##### Quality testing of the different loci

We observed 45 locus pairs in significant LD (14%), with 95%CI of the average per locus between 3 and 4. Again, we observed an important, highly significant and strongly variable *F*_IS_=0.299 in 95%CI=[0.181, 0.414] (*p*-value<0.0002). Neither *F*_IS_ or *H*_S_ differed significantly between the two cohorts. Null alleles were suggested by the *r*_SE_=30.5 and a positive, though not significant relationship between *F*_IS_ and *N*_blanks_. From this relationship, drawing the corresponding regression, we could deduce that for four loci, L027_Tri, L028_Di, L032_Di, and L053_Tetra, most missing data did not correspond to null homozygotes (same loci as for all polymorphisms). We consequently recoded all missing data, except for these four loci, as homozygous for the null allele and used FreeNA to compute null allele frequencies (*p*_nulls_). We needed to undertake several regressions *p*_nulls_∼*N*_blanks_ and recode missing genotypes as “0”, at some more loci, before this regression was maximized. Finally, missing data were recoded as 999999 only for loci L045-Tri, and L047-Tetra and missing data were left as such (000000) for all other loci for FreeNA analyses. We kept 15 loci, as presented in the supplementary file S3 (one more than with all polymorphisms, but see below). With these loci, the regression *F*_IS_∼*p*_nulls_ displayed very good performances (Figure 6), with a 95%CI wider than with all polymorphisms, but still smaller than with old loci. The intercepts were compatible with an effective population size of *N*_*e*_=28 in 95%CI=[3, Infinite].

**Figure 6.**
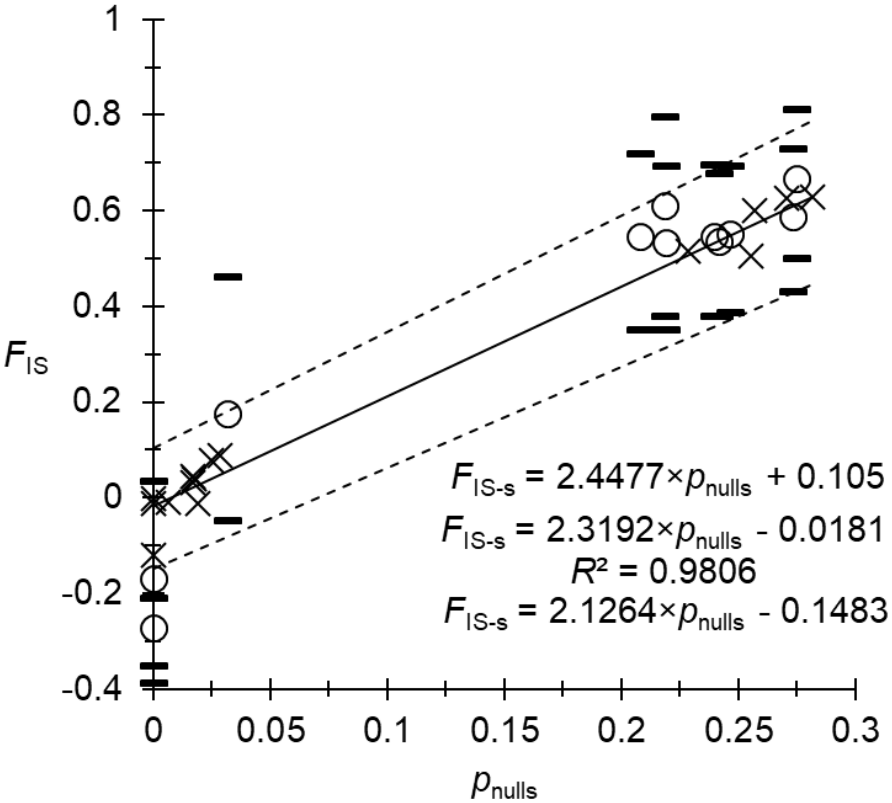
Regression of *F*_IS_ as a function of null allele frequencies (pnulls) (weighted averages across cohorts) at the different SSRseq loci (allele size polymorphism) for Glossina palpalis gambiensis from Boffa and Dubreka (Guinea), in cohorts C50 and C56 (two subsamples). Regression of averages is represented with a black line, and its 95% confidence intervals (5,000 bootstraps over individuals) with dotted lines. Equations (with slope and intercept), for average and confidence intervals, and determination coefficient (R^2^), for average, are also indicated. Problematic loci, represented with empty circles and black dashes, were excluded of the regression.

We tested for stuttering and SAD only for loci with a *F*_IS_>0. Only one locus (L035) displayed a significant stuttering signature (*p*_BH_<0.0001), but this locus was perfectly explained by null alleles. Indeed, according to null allele frequency, we expected an incidence of 7.7 expected null homozygotes (missing genotypes), while we observed 8 of those in the data (*p*_BH_=1). We observed no significant SAD signature (all *p*_BH_>0.26).

With the 15 retained loci, 12 pairs of loci (11%) displayed a significant LD, none of which stayed significant after BY correction (minimum *p*_BY_=0.11).

Finally, kept and excluded loci for SSRseq with all polymorphisms and allele size polymorphism are summarised in the Supplementary File S3.

##### Subdivision measures and testing

Between C50 and C56, the adequacy between *F*_ST_’ and *G*_ST_” was rather good (slope *b*=0.9326, intercept *a*=0.0015, *R*^2^=0.9501). Correcting for null alleles lead to *F*_ST-ENA_’=-0.0024 in 95%CI=[-0.0138, 0.0112] (*p*-value=0.6489).

In the Figure 7, the comparisons between the three markers kinds suggested that old markers produced more variable results and underestimated subdivision measure suggestive of homogenizing selection (all estimates below zero) between cohorts. Geographic subdivision between Boffa and Dubreka also appeared more centered on 0 with narrower 95%CI for SSRseq as compared to the results observed with old loci. Excluding GPCAG, which is suspected to respond to vector control with significant changes of its allele frequencies (Berté et al., 2019), did not change these observations (*F*_ST_=-0.003 in 95%CI=[-0.007, 0.004], p-value=0.9589). The low or absence of subdivision between the two foci implies free dispersal of tsetse flies in the whole mangrove from Boffa to Dubreka, i.e. along 50 km (distance between the two most remote sites) to 100 km for the maximum length of the mangrove containing Boffa and Dubreka.

**Figure 7.**
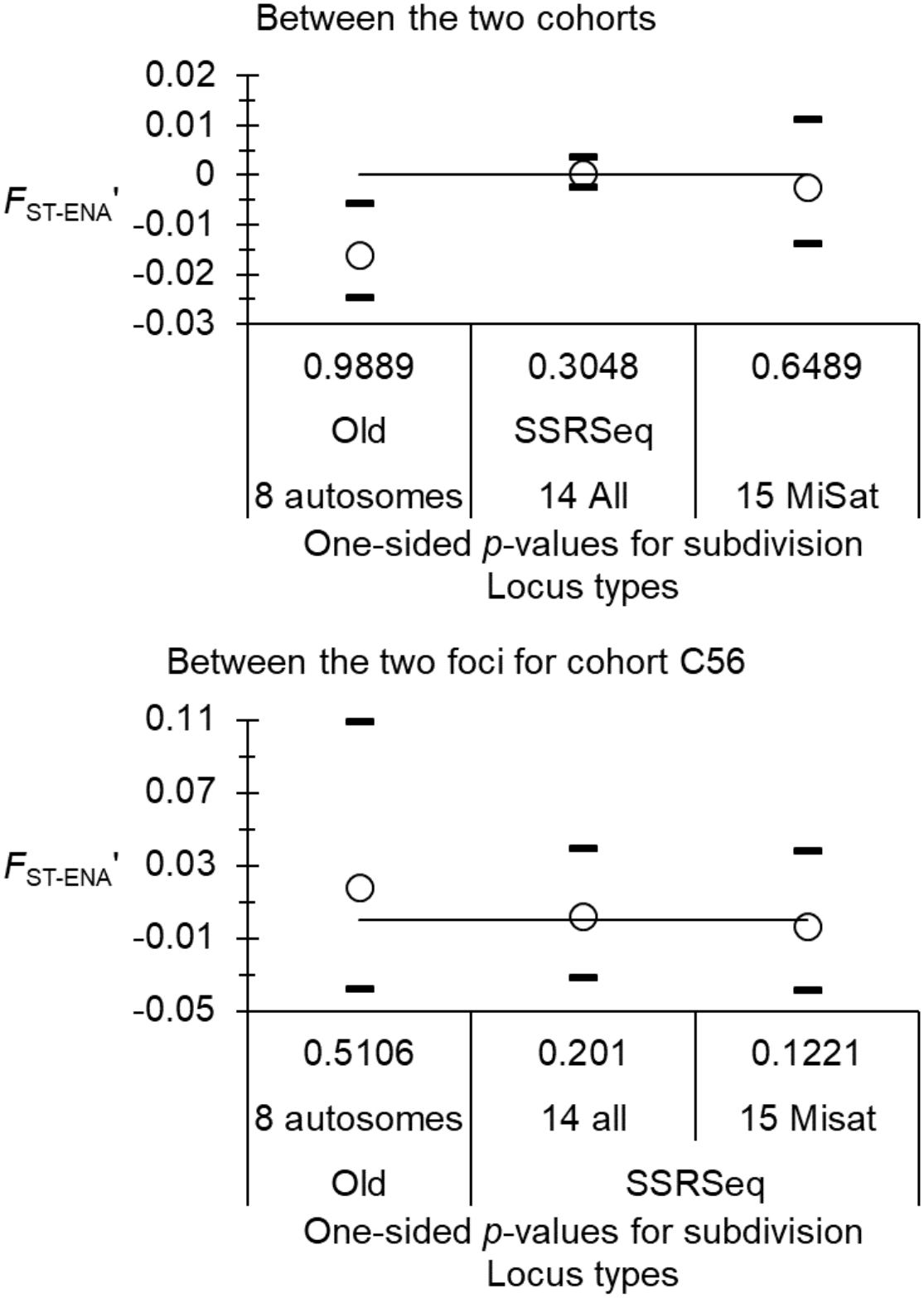
Comparisons between subdivision measures (empty circles), corrected for null alleles and polymorphism for the eight autosomal old loci, the 14 SSRseq best loci with all polymorphisms and the 15 best SSRseq loci with allele size variation only (MiSat) and their 95% confidence intervals obtained with 5,000 bootstraps over loci (black dashes) of *Glossina palpalis gambiensis* from Guinea.

##### Effective population sizes

We estimated the total surface of this mangrove at *S*_mangrove_≈2192 km^2^. The minimum surface defined by traps that captured at least one tsetse fly was *S*_traps_≈384 km^2^. Effective population sizes are presented in the Figure 8. Results obtained with harmonic means provided higher values than arithmetic ones, as predicted (Waples, 2024), but only with old markers, and not significantly so. Alternatively, SSRseq loci all provided significantly higher estimates with arithmetic means than with harmonic means. Temporal methods provided much higher estimates, as a result of the weak differentiation detected between the two cohorts, especially with old markers (by two to three orders of magnitude). Here again, Waples’ method provided the smallest estimate. We preferred not use this method further. Effective population sizes computed with SSRseq appeared the less variable, particularly so with allele size-based polymorphism. After a glance to Figure 8, it seemed reasonable to estimate that in the Boffa-Dubreka mangrove, the minimum effective population size approximately varies from 100 to 1,000 individuals.

**Figure 8.**
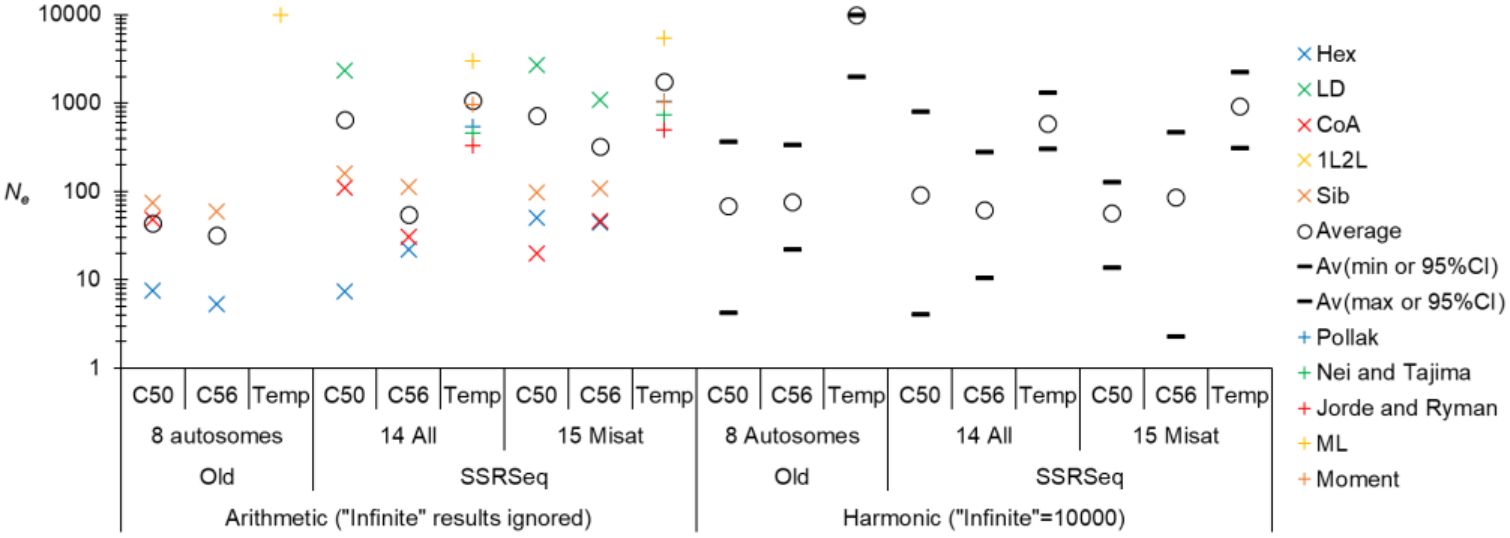
Effective population size estimated with single subsample (Single) and temporal (Temp) methods for *Glossina palpalis gambiensis* of the Boffa-Dubreka mangrove (Guinea), at cohorts C50, C56 and overall (temporal methods). Results obtained with eight autosomal old markers, the 14 best SSRseq loci with all polymorphisms (All) and the 15 best SSRseq loci with allele size polymorphism (MiSat) are presented. In the left (Arithmetic), individual values of each method are represented by crosses and pluses of different colors) and means by empty circles and averaged minimum and maximum values (min or max) (black dashes) (averages weighted with the number of usable values (“Infinite” outputs ignored. On the right (Harmonic) “Infinite” outputs were set at 10,000 and the harmonic mean (empty circles) computed across single sample or temporal methods with their 95% confidence intervals (black dashes).

We could then estimate effective population densities in the total mangrove as *D*_*e*-m_=[0.05, 0.5] individuals per km^2^ and, with *S*_traps_, *D*_*e*-t_=[0.25, 2.6] individuals per km^2^. These values are among the smallest found as compared to other studies (De Meeûs et al., 2019).

##### Conclusions on SSRseq in Gpg

Vector control had no visible effect on the genetic composition of this population. Nevertheless, it seemed that genetic drift had less impact over time than over space (though not significantly so, Figures 7 and 8), which may reflect the effect of vector control combined with recolonization of sites by tsetse flies from very variably distant locations. Vector control efficiently protects human hosts from infection (Courtin et al., 2015), but was not able to reduce significantly the demography of this vector in the Guinean Mangrove, as a result of its very large population (geographically and demographically), occupation of inaccessible zones, and ability to disperse at fairly long distances. For this species, in this geographic zone, SSRseq provided more reliable results than old markers. The comparison between the two datasets with all or allele size polymorphisms did not reveal clear differences of performances. Nonetheless, allele size polymorphism permitted retaining one more locus after the quality testing procedure (but see the Conclusions section). Allele size variations also allowed testing for stuttering and/or SAD (very difficult or even impossible to test with all polymorphisms), and to find that SSRseq markers did not show any evidence of such amplification issues.

### SSRseq loci in Gpp from the Bonon focus in Côte d’Ivoire

#### Detection of significant levels of subdivision by the Wahlund effect procedure

According to Figure 9, only traps displayed a significant subdivision signal, for all parameters with all polymorphisms. This was not confirmed with allele size polymorphism (Figure 10), though *F*_ST_ was still maximum with traps, with a significant subdivision signature (*p*-value=0.0348).

**Figure 9.**
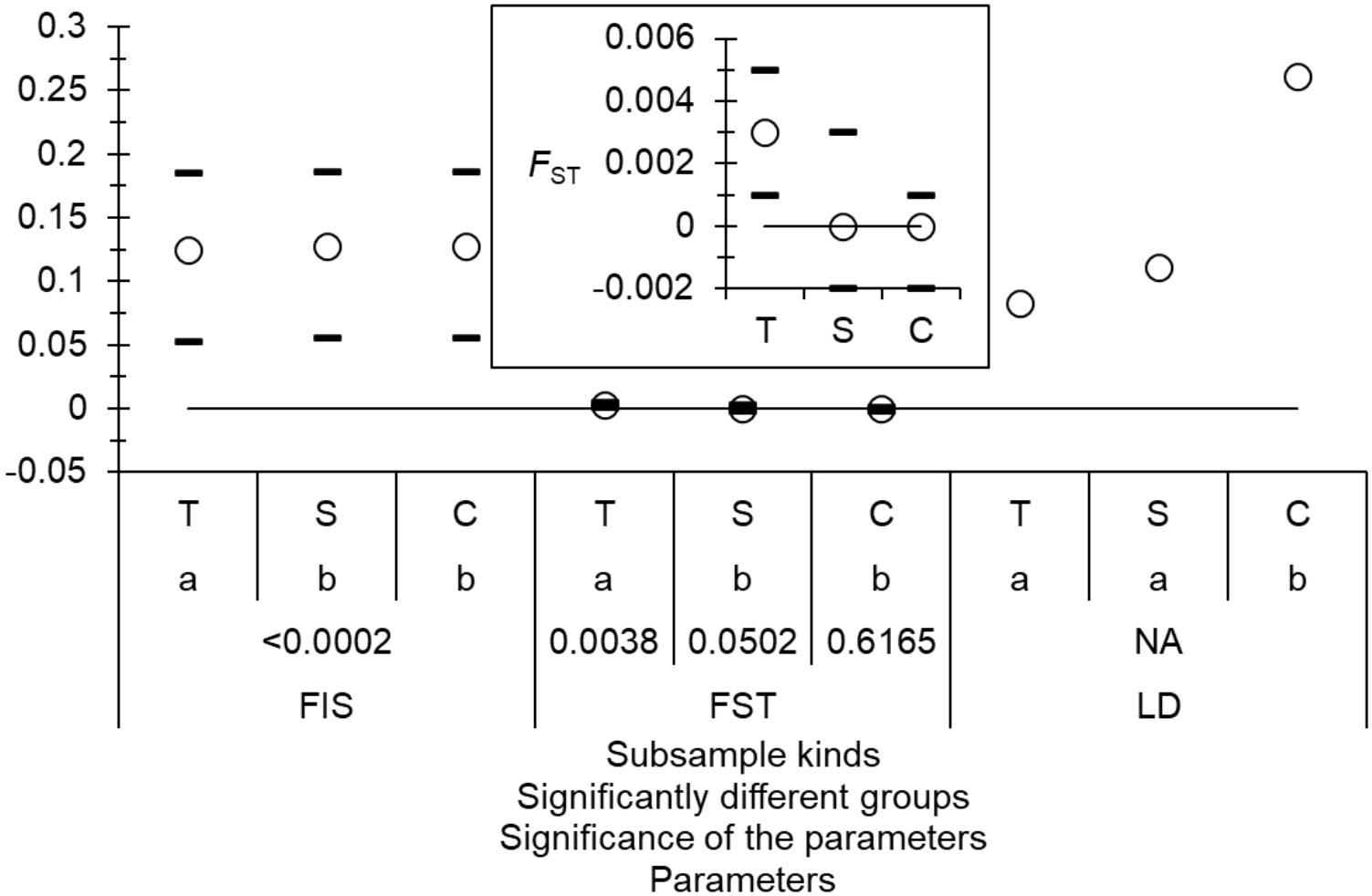
SSRseq loci with all polymorphisms for *Glossina palpalis palpalis* from Bonon (Côte d’Ivoire). Comparisons of different sampling designs regarding *F*_IS_, *F*_ST_ and the proportions of locus pairs in significant linkage (paired tests). T, S, and C correspond to the sampling designs: Traps, Sites, and Cohorts respectively. Small captions (a, b) correspond to groups of non-significant paired tests (sharing one letter). Paired tests were adjusted for repetitions with the BH procedure. Significance of *F*_IS_ and of subdivision are also indicated.

**Figure 10.**
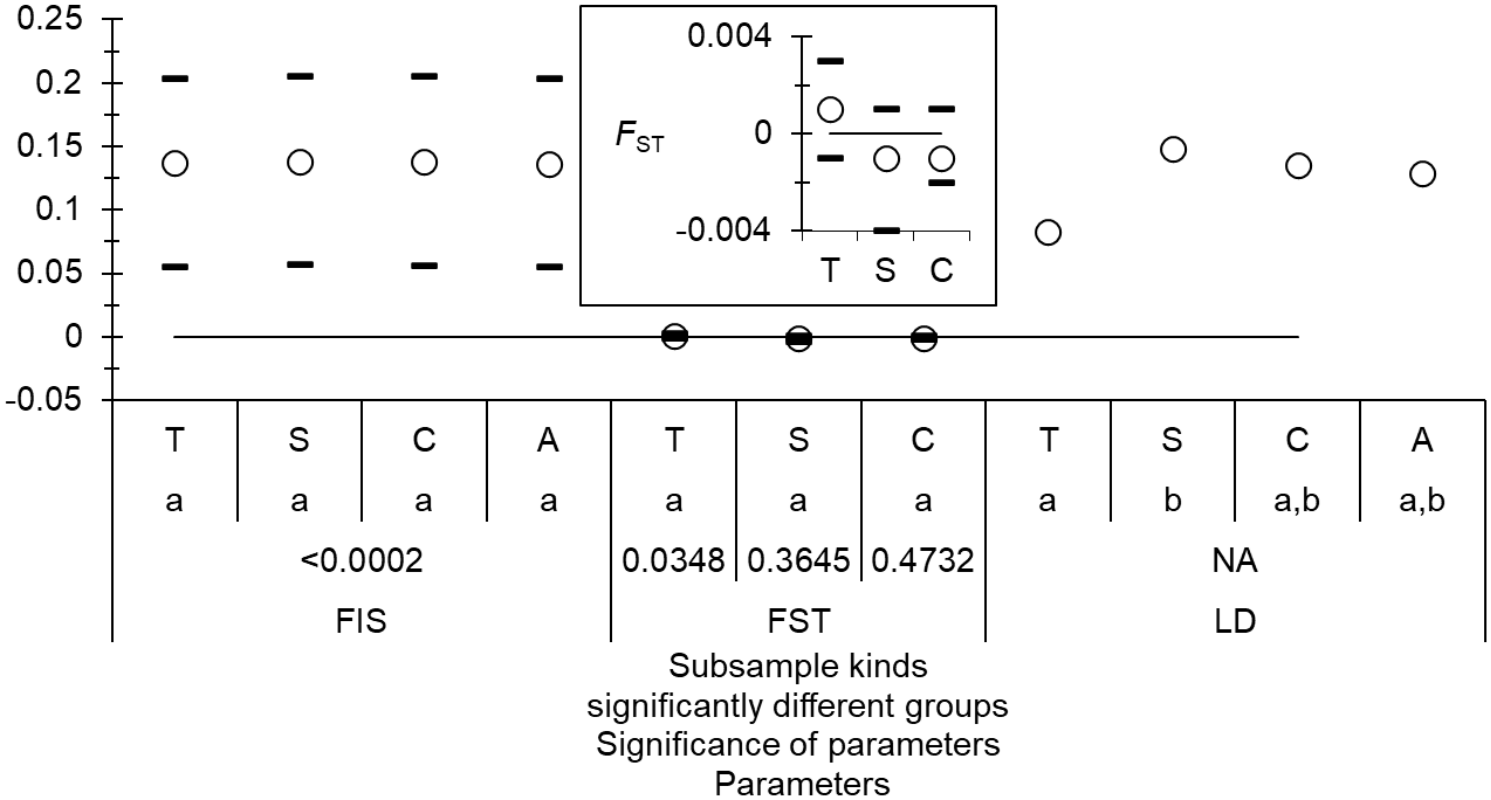
SSRseq loci with allele size polymorphism for *Glossina palpalis palpalis* from Bonon (Côte d’Ivoire). Comparisons of different sampling designs regarding *F*_IS_ (paired comparisons), *F*_IS_ and *F*_IT_ in the two cohorts, *F*_ST_ and the proportions of locus pairs in significant linkage (paired Fisher’s exact tests). T, S, C, and A correspond to the sampling designs: Traps, Sites, Cohorts, and All in one sample (computed as *F*_IT_ for *F*_IS_) respectively. Paired tests were adjusted for repetitions with the BY procedure and no comparison appeared significant. Significance of *F*_IS_ and subdivision are also indicated.

Consequently, unless specified otherwise, we only retained trap as representative of the subsample unit in the following analyses.

#### Quality testing of the different loci

We could not extensively test for LD to get precise enough *p*-values with 10,000 randomizations as the procedure took too much time and was aborted after four days. With 1,000 permutations, the process lasted 3 days. We could not control the false discovery rate, as the minimum reachable *p*-value=0.001 would not have remained significant at the BH level. We observed 27 significant tests (8%) (All and allele size polymorphisms). Four loci needed being removed: Locus L027 for the same reason as before (almost fixed heterozygosity), L032, L044 and 59 with too many missing data (25, 36 and 29 respectively).

Keeping all loci, within each trap-cohort combination, led to many unreliable results. This was due to the excess of missing data at some loci, combined to subsample sizes that could be very small in several instances (1-4 individuals). Before quality testing of loci, we decided to exclude all loci with more than 1% missing data, keeping loci with 0, 1 or 2 missing genotypes. We thus kept nine loci: Loci L037, L039, L040, L041, L048, L049, L050, L051, and L054.

With the remaining 9 loci, only one (3%) and two (6%) locus pairs appeared in significant LD with all and allele size polymorphisms respectively, which did not stay significant after BH correction. There was a substantial and significant heterozygote deficit: *F*_IS_=0.11 in 95%CI=[0.04, 0.18] (*p*-value<0.0002) in both datasets against *F*_IS_=0.16 in 95%CI=[0.072, 0.248] in females with old loci. Several rounds of attempts were needed to obtain the best strategy to recode missing data for FreeNA analyses. Blank genotypes were all recoded as null as missing data for all polymorphisms and 999999 for locus 050 only for allele size polymorphism. Following this, the regression *F*_IS_*∼p*_nulls_ provided an almost perfect result. since the model explained 98% of *F*_IS_ variation and the intercept was slightly negative (*F*_IS-0_=-0.0067 or *F*_IS-0_=-0.0306 for all and allele size polymorphisms respectively) (Figures 13 and 14). These figures are compatible with a random mating dioecious population of size *N*_*e*_=75 or *N*_*e*_=16 individuals for all or allele size polymorphisms respectively.

Finally, kept and excluded loci for SSRseq with all polymorphisms and allele size polymorphism are summarised in the Supplementary File S3.

#### Subdivision measures and testing

, We tested for the effect of vector control (VC) from T0 (before VC) and at different times, on *H*_S_, *F*_IS_ and *F*_ST_ (Friedman’s sum rank test over all, and Wilcoxon signed rank test between paired cohorts) and with Fisher exact tests for LD (all two-sided). For this we excluded loci with too many missing data (L027, L032, L044 and L59), but kept the 11 loci excluded after quality testing. The reason for that is that some of these might be under some kind of selection (if any), and possibly linked to vector control. The results obtained with the 22 loci are shown in Figures 11 (all polymorphisms) and 12 (allele size) and suggested some subtle effects of VC, probably resulting from the recolonization by individuals from more or less distant sites (except for *F*_ST-ENA_’ with allele size polymorphism). None of these differences could be detected with old loci, except with LD and only with Friedman test (*p*-value=0.0271).

**Figure 11.**
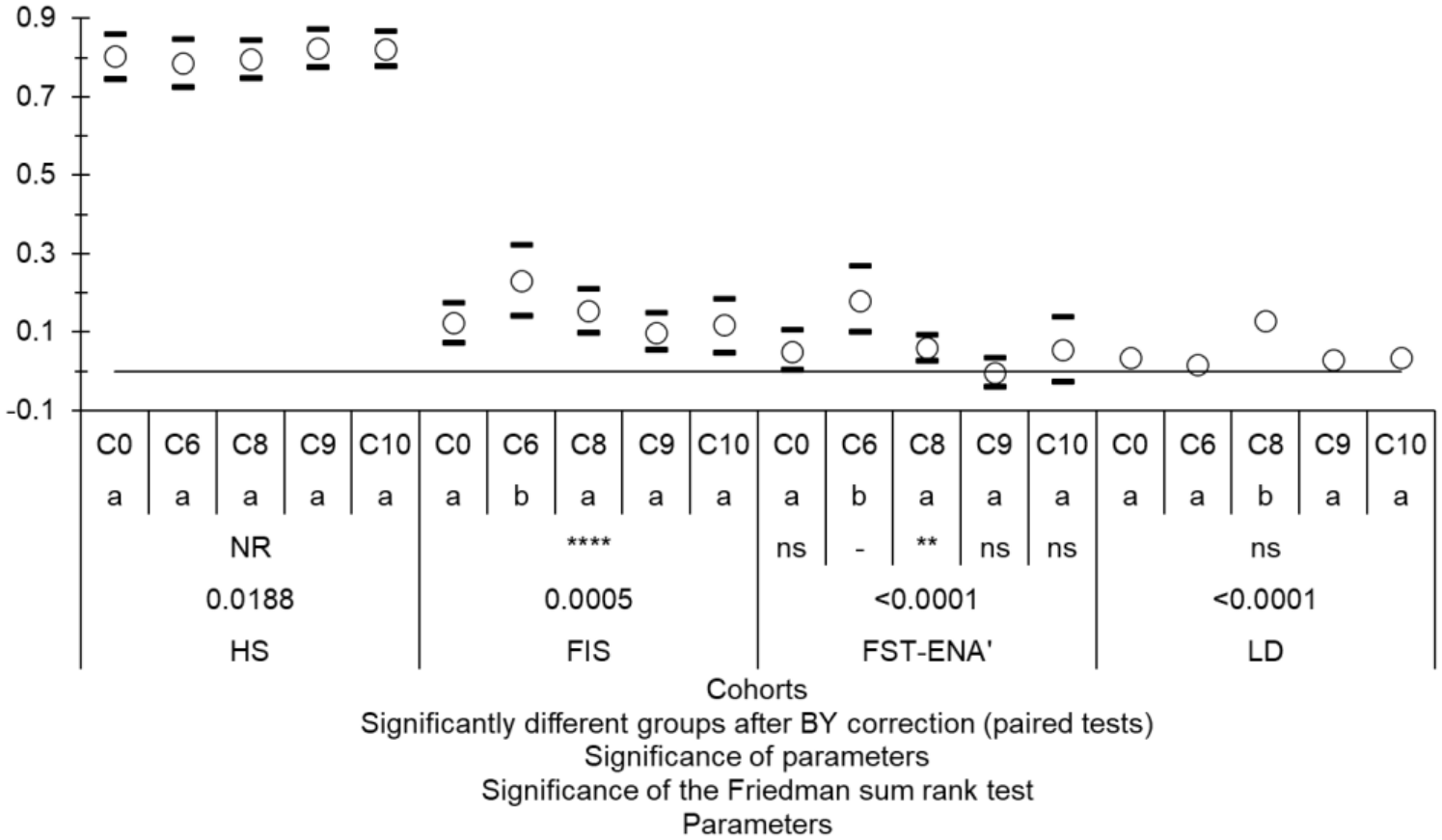
Comparisons of population genetics parameters between different cohorts (C0 before control, and C6-C10) of *Glossina palpalis palpalis* from Bonon (Côte d’Ivoire) obtained for the best 22 SSRseq loci with all polymorphisms. Significant paired differences with BH adjustment are indicated with different letters. Paired tests were signed rank test for all but the proportion of significant linkage disequilibrium tests (LD, Fisher exact tests). The significance of departures from 0 are also indicated when relevant, as is the result of the Friedman test overall cohorts.

The nine kept and the 17 excluded loci for SSRseq with all polymorphisms and allele size polymorphism are summarised in the Supplementary File S3.

With the nine loci kept after quality testing process, temporal subdivision appeared weak and not significant between any pair of cohorts for the same trap (minimum *p*_BH_=0.088), and the correlation between *D*_CSE-FreeNA_ and the temporal distances appeared small (*r*=0.1101 and *r*=0.1593 for all and allele size polymorphisms respectively) and not significant (Mantel tests, *p*-values=0.22).

We found a significant isolation by distance at C0, with slope *b*≈0.01 in 95%CI≈[0.005, 0.015] over the three marker kinds (old and SSRseq loci with all and allele size polymorphisms), leading to a neighborhood size *Nb*≈100 in 95%CI≈[67, 200], and a number of immigrants from neighboring subpopulations (around traps) *N*_*e*_*m*≈16 in 95%CI≈[11, 32]. After the beginning of control, we observed a progressive disappearance of the signature of isolation by distance (Figure 15). This was particularly clear from SSRseq with allele size, less so with all polymorphisms and not at all with old markers for which the signature disappeared at C6, and reappeared at C8. This suggested a recolonization of subpopulations with flies from anywhere in the trapping zone or even from the surrounding areas. This could be in line with the increase of locus pairs in significant LD (see Figures 11 and 12). This also suggested a higher sensitivity of SSRseq markers, with a small advantage for those with allele size polymorphism.

**Figure 12.**
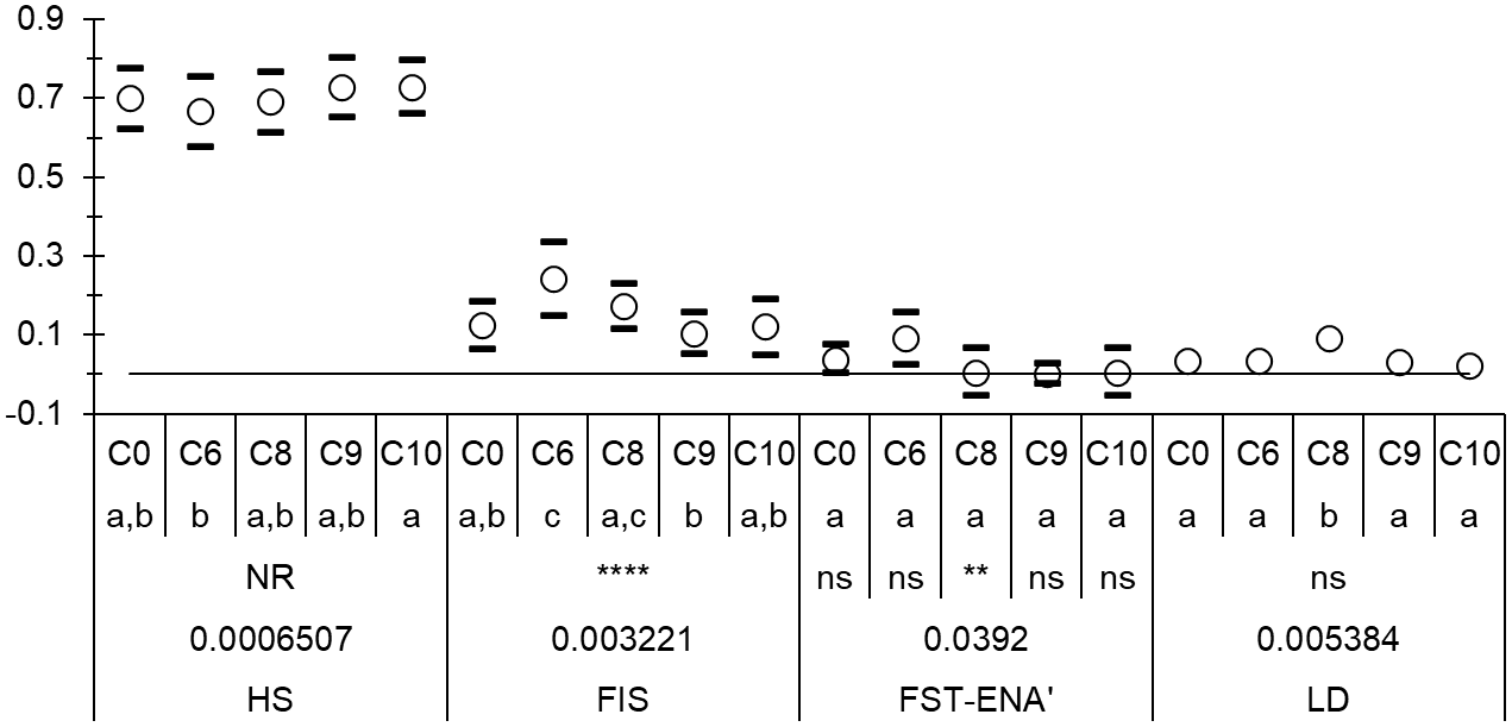
Comparisons of population genetics parameters between different cohorts (C0 before control, and C6-C10) of *Glossina palpalis palpalis* from Bonon (Côte d’Ivoire) obtained for the best 22 SSRseq loci with allele size polymorphism. Significant paired differences with BH adjustment are indicated with different letters. Paired tests were signed rank test for all but the proportion of significant linkage disequilibrium tests (LD, Fisher exact tests). The significance of departures from 0 are also indicated when relevant, as is the result of the Friedman test overall cohorts.

**Figure 13.**
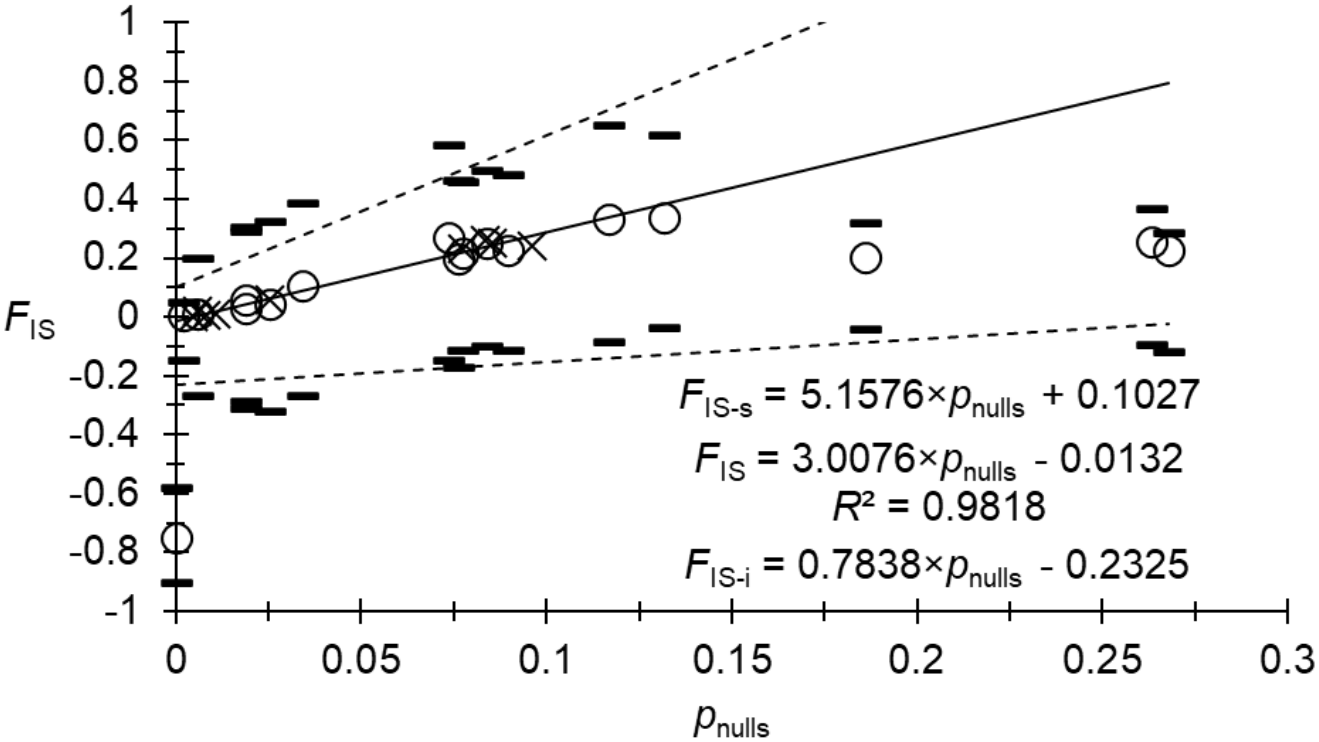
Regression of *F*_IS_ as a function of null allele frequencies (*p*_nulls_) (weighted averages across cohort-trap subsamples) at the different SSRseq loci (all polymorphisms) for *Glossina palpalis palpalis* from Bonon (Côte d’Ivoire). Regression of averages is represented with a black line, with its 95% confidence intervals (5,000 bootstraps over individuals) with dotted lines. Equations (with slope and intercept), for average and confidence intervals, and determination coefficient (*R*^2^), for average, are also indicated. Problematic loci, represented with empty circles and black dashes, were excluded from the regression.

**Figure 14.**
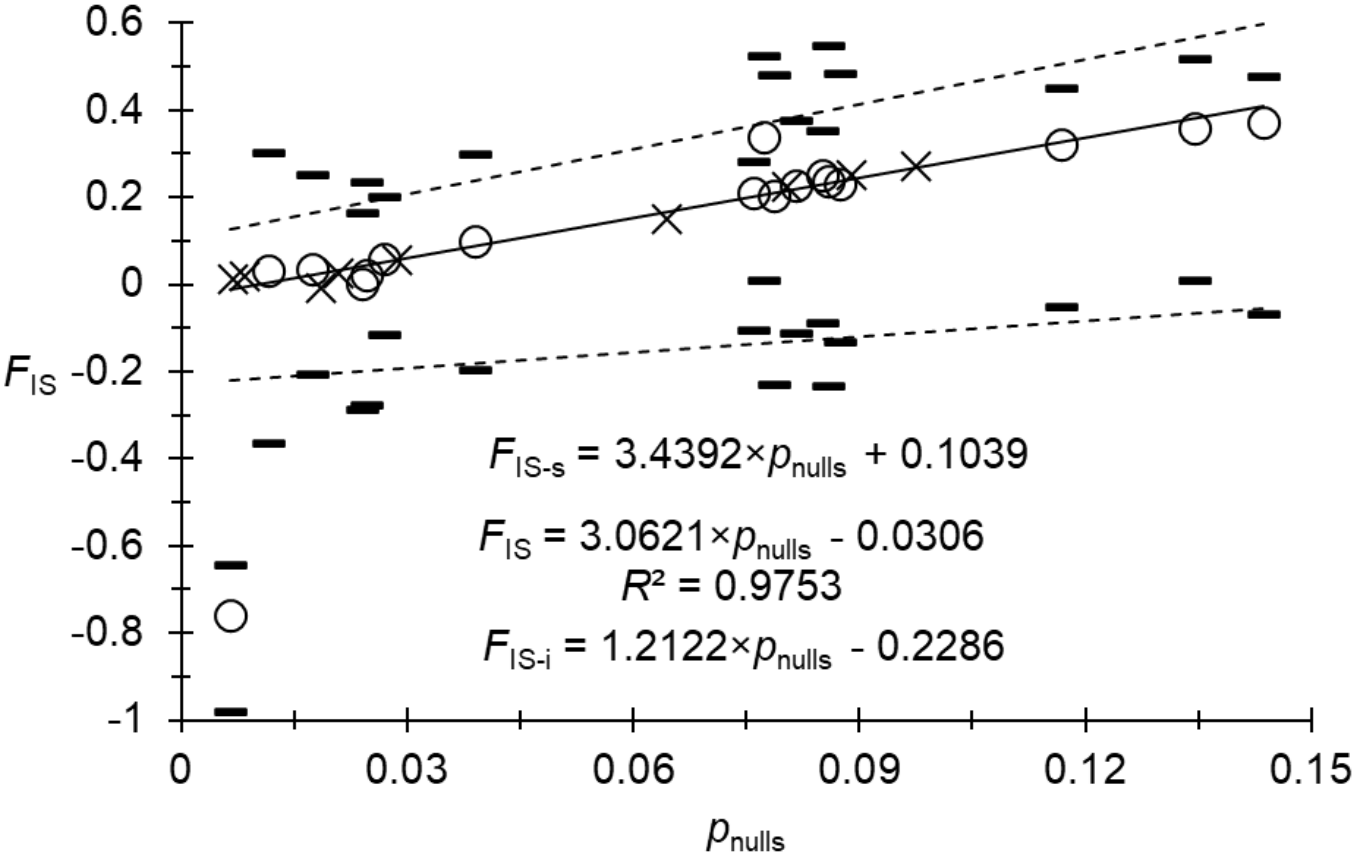
Regression of *F*_IS_ as a function of null allele frequencies (*p*_nulls_) (weighted averages across cohort-trap subsamples) at the different SSRseq loci (allele size polymorphism) for *Glossina palpalis palpalis* from Bonon (Côte d’Ivoire). Regression of averages is represented with a black line, with its 95% confidence intervals (5,000 bootstraps over individuals) with dotted lines. Equations (with slope and intercept), for average and confidence intervals, and determination coefficient (*R*^2^), for average, are also indicated. Problematic loci, represented with empty circles and black dashes, were excluded from the regression.

**Figure 15.**
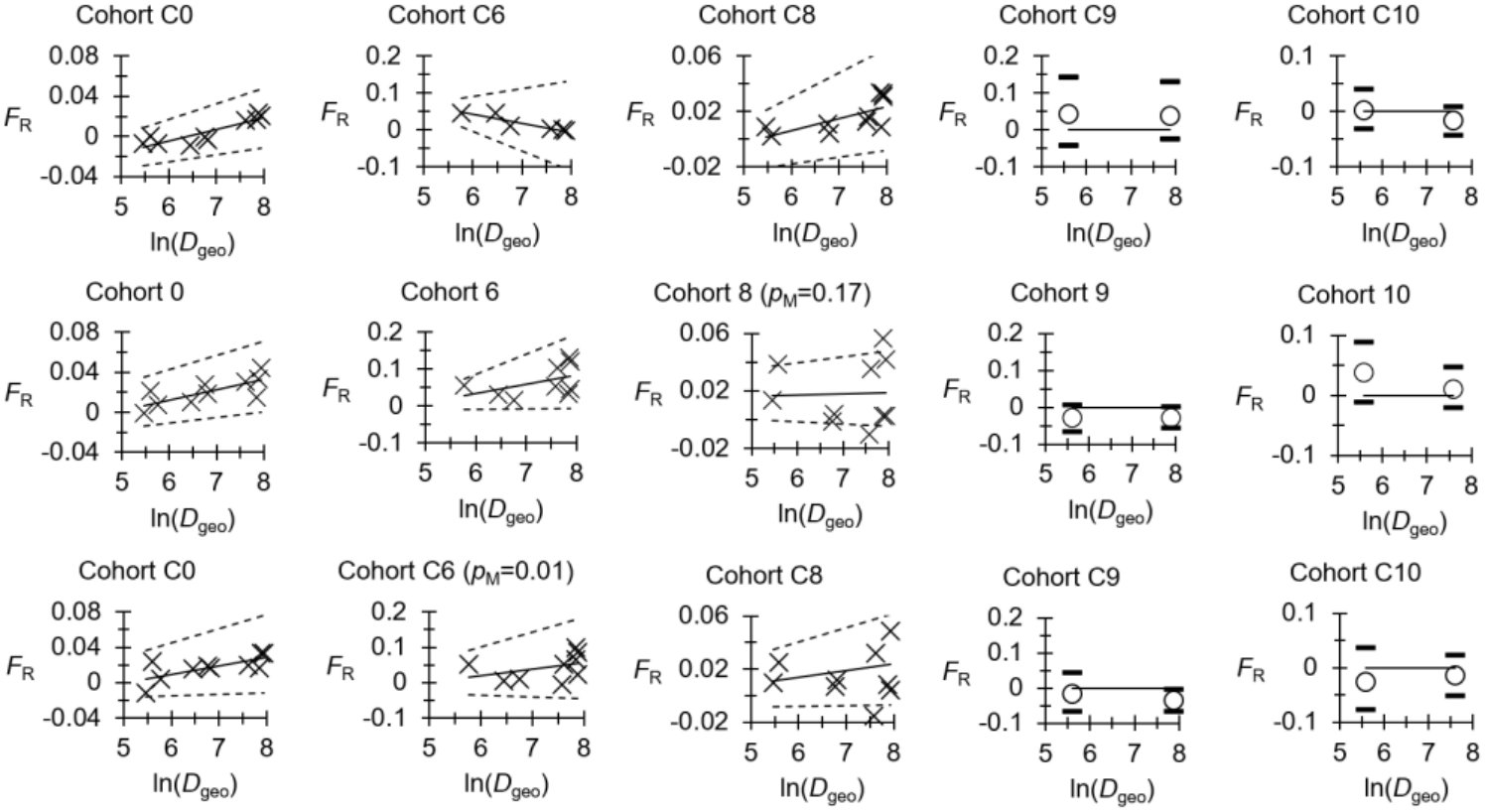
Results of Rousset’s model for isolation by distance between the natural logarithm of geographic distances (in m) (ln(*D*_geo_) and Roussets genetic distance corrected for null alleles *F*_R_=*F*_ST-ENA_/(1-*F*_ST-ENA_) (plain line) and its 95% confidence intervals (dotted lines), for *Glossina palpalis palpalis* from Bonon (Côte d’Ivoire) using the nine old loci on females (top), nine SSRseq loci with all polymorphisms (middle) and SSRseq loci with allele size polymorphism (bottom). Regression models were undertaken when a reasonable cloud of points was available. Otherwise, we only represented *F*_R_ (empty circles) and its 95% confidence interval (black dashes), averaged across close by traps. The slope (*b*) and the width of its confidence interval (Δ_*b*_) at C0 happened to be similar for each marker kind (*b*≈0.01 in 95%CI≈[0.005, 0.015]).

#### Effective population sizes

A synthesis of effective population size estimates is provided in the Figure 16. A tendency for decrease after the beginning of control is apparent with SSRseq loci with allele size polymorphism only, but not significantly so. Old loci and SSRseq loci with all polymorphisms tended to provide more variable results as compared to SSRseq loci with allele size polymorphism, which, in particular, offered the best consistency between single sample methods at C0 and temporal methods, in line with the absence of a significant genetic signature of VC activities. Globally, we infered that this tsetse population roughly displays a minimum effective population size between 100 (the neighborhood size) and 400 individuals.

**Figure 16.**
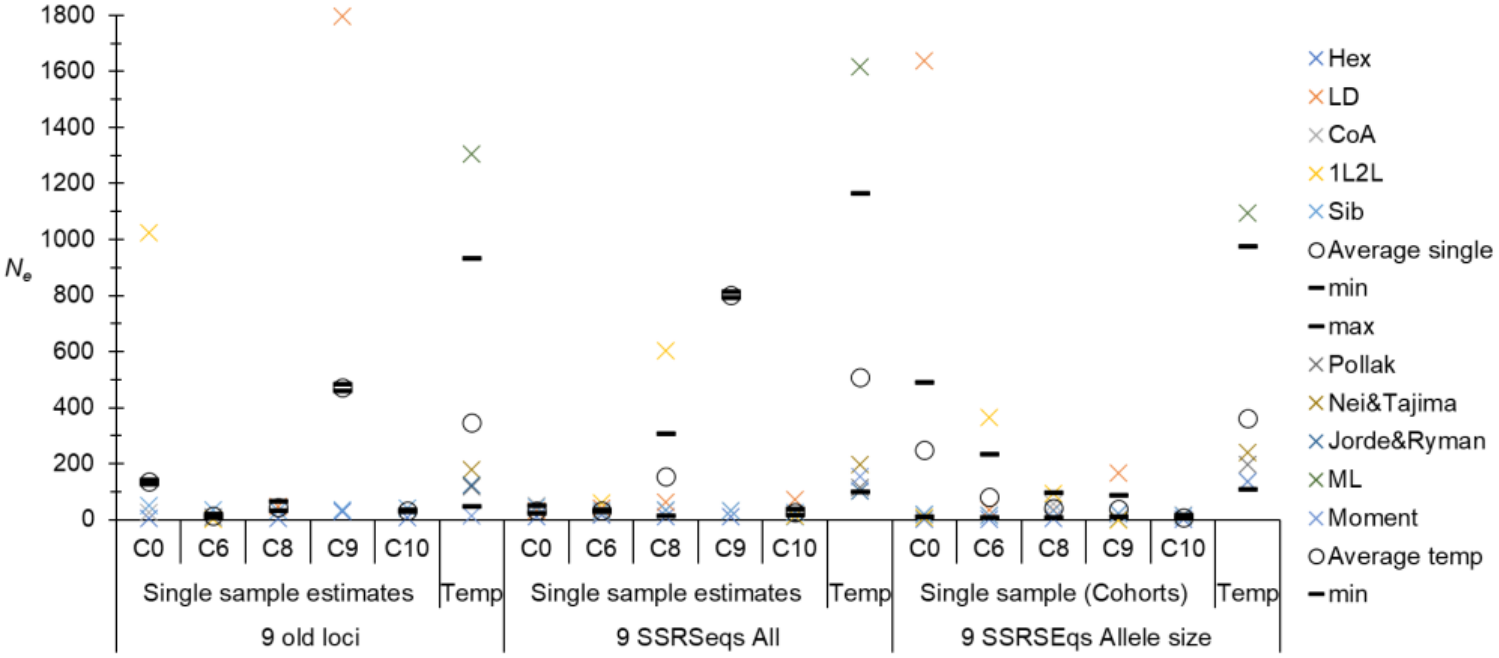
Effective population size estimates with different methods (crosses of different colors) and averages (weighted with the number of usable values) (empty circles) and average minimum and maximum values (black dashes), for different cohorts (C0 to C10) and for temporal methods (Temp). Estimates are provided with the nine best loci for females with old loci, SSRseq with all polymorphisms (All) and with allele size polymorphism.

#### Dispersal

Taking the two most distant traps provided a maximum distance of *D*_geo-max_=2.8 km. This never outputted a significant subdivision, meaning that tsetse flies easily travel that far, even if we could detect an isolation by distance signature at that scale. We here considered this distance as the diameter of the disc that contains all traps used in this study, with a surface *S*_min_=*π*×(*D*_geo-max_/2)^2^=6,144 km^2^. If we consider that SSRseq with allele size polymorphism provided the most reliable results, we could estimate, for C0, an average dispersal distance of *δ*=1 km/generation in 95%CI=[0.6, 2], and a maximum effective population density *D*_*e*_=*N*_*e*_*/S*_min_=55 flies/km^2^ in minimax=[2, 109]. Such figures are in the same order of magnitude as those observed with old loci: *δ*=1.1 km in 95%CI=[1, 1.4] and *D*_*e*_=22 in minimax=[20, 23]. Note that these quantities are a little different from those published in Berté et al (Berté et al., 2019) as we here used a dataset with more loci, only for C0 and with another surface for the population, which did not change much our estimates except for the densities.

## Conclusions

Generally speaking, SSRseq loci appeared much more polymorphic in these two tsetse fly species than what has be found in other organisms (Lepais et al., 2020; Soliani et al., 2025) (Figure 17). The origin of such differences is unclear. It may come from much bigger effective population sizes in tsetse flies as compared to other species, which is not very intuitive, given the life cycle of this viviparous insect (Solano et al., 2010) (but see below). Alternatively, a nonexclusive other explanation may be that the genome of tsetse flies displays mutation rates that are much higher than for the other organisms presented in the Figure 17. This may provide a higher adaptive flexibility to these vectors of important diseases.

**Figure 17.**
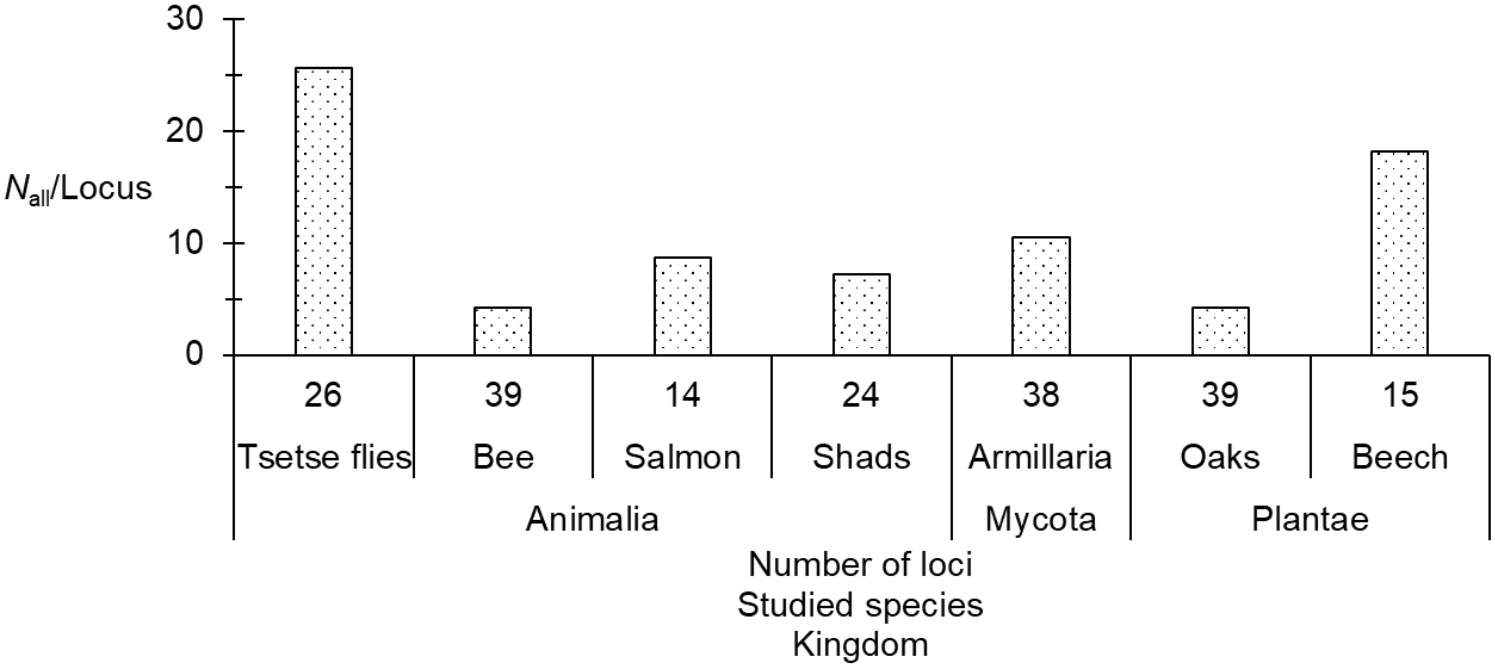
Average number of alleles found per SSRseq locus (all polymorphisms) for different species: Two species of tsetse flies (present paper); one species of a Caribbean bee, the Atlantic salmon (fish), two species of shads (fishes), one fungus and two species of Iberian oaks (Lepais et al., 2020); and one South-American beech (Soliani et al., 2025).

After pre-selection, only 26 loci were kept, with a minority of dinucleotide, due to their lack of reliability (sequencing errors, stuttering and/or null alleles). After data analyses, we had to exclude several loci that presented various problems: evidence of selection, excessive number of missing data, excessive frequencies of null alleles or something else that we were not able to identify but clearly gave outlier results. Loci under selection were detected as such because of subdivision between cohorts during a vector control campaign (VCC). These may then be responding to devices used in VCC. Some of these loci are located in coding genes that may be involved in resistance, but many others are not in or close to a known gene (Supplementary File S3). One locus, L027 (a trinucleotide locus) was found in a huge excess of heterozygotes that may suggest overdominant selection. According to the annotated genome of *Glossina fuscipes* (the first annotated genome we found), Locus 027 is located in an intron of a gene implied in the regulation of gene expression. For this locus, one allele, allele 105 in both tsetse species, was almost present in all individuals. This may also suggest that this locus probably displayed a fake polymorphism. Further studies are needed to understand precisely what underlies the patterns exhibited by this locus. Another interesting locus was Locus 047 (a tetranucleotide), which appeared more often than the others in LD with another locus, in particular in *G. p. gambiensis*, and presented an obvious tendency for homogenizing selection in that species, though marginally not significantly so with allele size polymorphism. This locus also presented more missing data than the other loci, especially so in *G. p. palpalis*. Locus 047 is located close to a gene that controls for odor avoidances, in particular phthalates that are present in plastics and insecticides, and may thus be related with vector control. We did not have samples at T0 in Guinea (before vector control measures), as our first sampling time there was 50 generations after the beginning of control. On the contrary, in Bonon, Côte d’Ivoire,T0 flies were available, but the most distant sample in time was only 12 generations after the beginning of control. These differences may partly explain the differences observed at locus 047 between these two sites, in particular regarding subdivision between cohorts. Another factor is the subdividing importance of traps in Bonon for *G. p. palpalis*, while *G. p. gambiensis* did not display any genetic subdivision signature at the scale of the two Guinean foci explored in the present survey.

More loci needed being removed in *G. p. palpalis* as compared to *G. p. gambiensis* as more problems of reliability of parameter estimates appeared in the first species. This is partly due to the fact that traps appeared as a relevant subdivision (though with a very weak signal) in Bonon, which lead to moderate to very small subsamples, and not in the Guinean mangrove. In Guinea, the cohort could be kept as the subsample unit, guaranteeing much bigger subsamples. Nevertheless, missing data and null alleles were more common in *G. p. palpalis* in particular in dinucleotide loci that had to be all removed, with some other loci. Both tsetse species, with an equal amount of DNA, were used to build the SSRseq library, so this difference can only be attributed to chance or to a better quality of the DNA from Guinean tsetse flies. Nevertheless, for further studies using such loci, and because of the large amount of available and highly polymorphic loci, we would advise keeping loci with less than 1% missing data, which led to suitable results in *G. p. palpalis*.

Generally, these new SSRseq markers provided comparable but more accurate and stable results to what was found with old markers. We selected 9 to 14 loci, all on autosomes, without short allele dominance, stuttering and rarer null alleles. It confirmed the high connectivity, and thus dispersal capabilities, of tsetse flies at the scales investigated. The very high level of polymorphism maintained, suggested higher effective population sizes than what could be estimated. Using equation (7.8e) (page 380) in Hedrick’s book (Hedrick, 2005a), and mutation rates *u*=0.001 in minimax=[0.0003, 0.002] on average (Ellegren, 2004), with the number of alleles found at the best SSRseq loci with allele size polymorphism (*n*_a_≈12 for both species), we have estimated that, for both species, *N*_*e*_≈3000 in minimax=[1400, 9200], or, alternatively, mutation rate was *u*≈0.004 in minimax≈[0.001, 0.147] in *G. p. gambiensis*, and *u*≈0.01 in minimax≈0.006, 0.25] in *G. p. palpalis*. Real effective population sizes and mutation rates probably lies between these two extremes.

Finally, allele size polymorphism provided slightly more accurate results, for a reasonable cost (around 14€ per individual at 26 loci). This would be less expensive than for old microsatellite (12.60 € per individual) because SSRseq results are delivered as such without any supplementary bench work. According to what we could find, 40 SNPs would cost around 5.2 € per sample (Hipp, 2016). Since, at the very least, 200 SNPs are required for robust population genetics analyses (Séré et al., 2017) (and generally more), this would sum to 26 € per sample, without counting the time spent in bioinformatic processing and attached salary or invoice costs. SSRseqs thus represent a much cheaper and efficient alternative. For further studies with SSRseq loci, to save time, we thus advise using allele size polymorphism on loci with less than 1% missing data. A potentially feasible and valuable next step is to transfer the SSRseq loci developed during the present study to other species of tsetse flies of the palpalis group (e.g. *G. fuscipes* and *G. tachinoides*).

## Appendices

### Appendix 1: Controlling the false discovery rate in multiple tests series

Systematic comparisons between all pairs of subsamples will produce a *k* tests series with some degree of dependency, where a number of significant *p*-values is expected to occur simply by chance. This would suggest using Benjamini and Yekutieli (BY) (Benjamini & Yekutieli, 2001) adjustment. Nevertheless, after reading the relevant literature more thoroughly, it appears that BY is highly conservative, and maybe prohibitively so (Reiner et al., 2003), especially for tests on proportions (De Meeûs, 2014). Moreover, for positively or moderately negatively dependent test statistics, BY presents no real FDR advantages as compared to Benjamini and Hochberg’s method (Benjamini & Hochberg, 2000) (BH), which is then better to be used instead (Benjamini & Yekutieli, 2001; Reiner et al., 2003). Negatively dependent tests means that the strength of H1 (with a better chance of small *p*-values) of some tests in the series implies that other tests in the series will tend to have also small *p*-values, while other sets of tests are undertaken under H0 (with an expected uniform distribution of *p*-values centered around 0.5). Typically, in a post-hoc multiple testing after an analysis of variance or after multiple comparisons of means (or medians), we expect such moderate negatively correlated *p*-values. Nevertheless, in some instances, such correlation may be strong enough to violate the assumptions needed to apply BH. This would be the case, for population genetics analyses, when different sets of loci are in strong LD (due to severe selection and/or physical linkage) and others are not. This is also what would be observed during differentiation tests between different pairs of subsamples taken from a population under isolation by distance, with a lot of pairs highly significant because from remote sites, and other pairs not so different (only by chance) because computed between close by sites. In such instances, detecting the true significant tests would require the use of BY. In all other cases, BH is expected to give robust and more powerful results.

### Appendix 2: Equilibrium values in an *n* Island model

Let us assume a population of *n* subpopulations of *N* diploid individuals, with a selfing rate *s*, a mutation rate *u* among *k* possible alleles, and a migration rate *m*. Let us define several parameters:

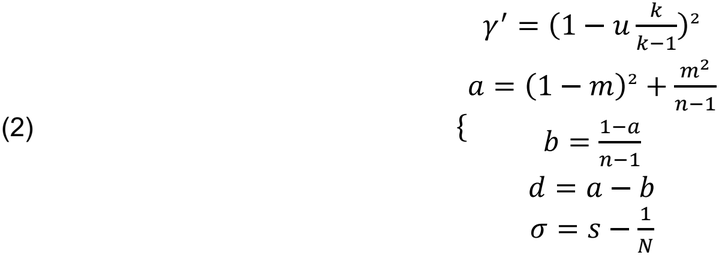

⬄

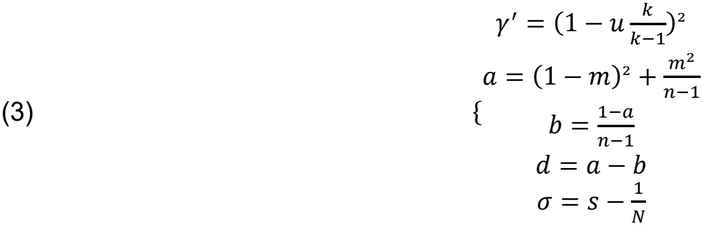

According to Rousset (Rousset, 1996), at drift-mutation-migration equilibrium, Wright’s *F*_ST_ writes:

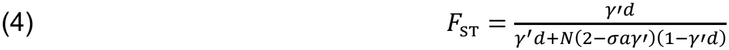

If we assume that the number of alleles is big and mutation rate is small enough, we can see that *γ*’→1. Moreover, if we consider local panmixia, then *s*=1/*N* and *σ*=0. We can thus write:

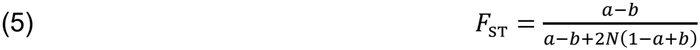

⬄

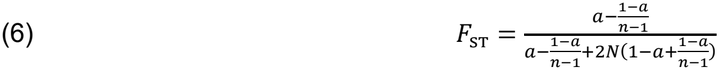

⬄

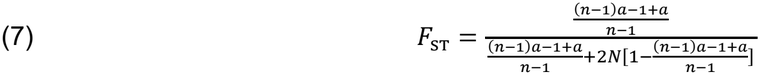

⬄

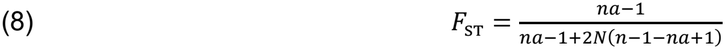

⬄

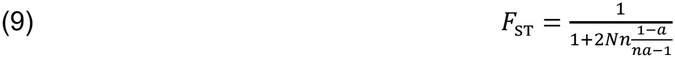

⬄

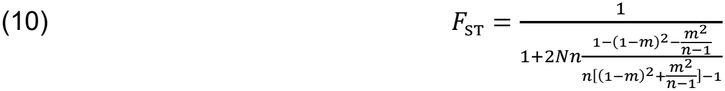

⬄

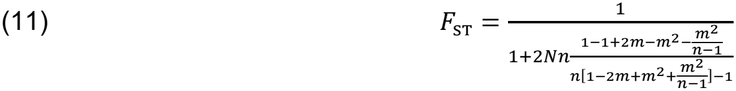

⬄

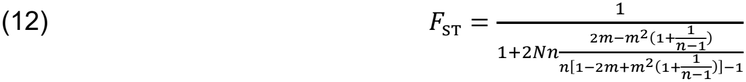

⬄

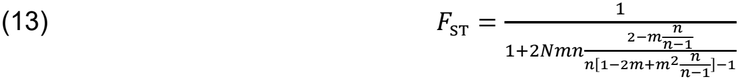

If we now assume that *m*<<1, we can write:

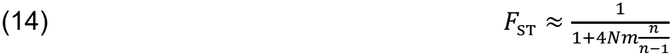

which is the same formula as in the infinite Island model, but with an increased migration rate by a factor *n*/(*n*-1). From there, it is easy to see that, for any population for which each subpopulation displays an effective population size *N*_*e*_:

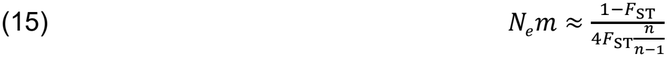

QED.

Figure A1 displays the different values for *N*_*e*_*m* as a function of *F*_ST_ for *n*=3. Negative estimates of *F*_ST_ will also translate into negative number of immigrants. This is interpreted as free migration between subsamples if not too negative. Strongly negative values can only be observed in case of homogenizing selection.

**Figure A1.**
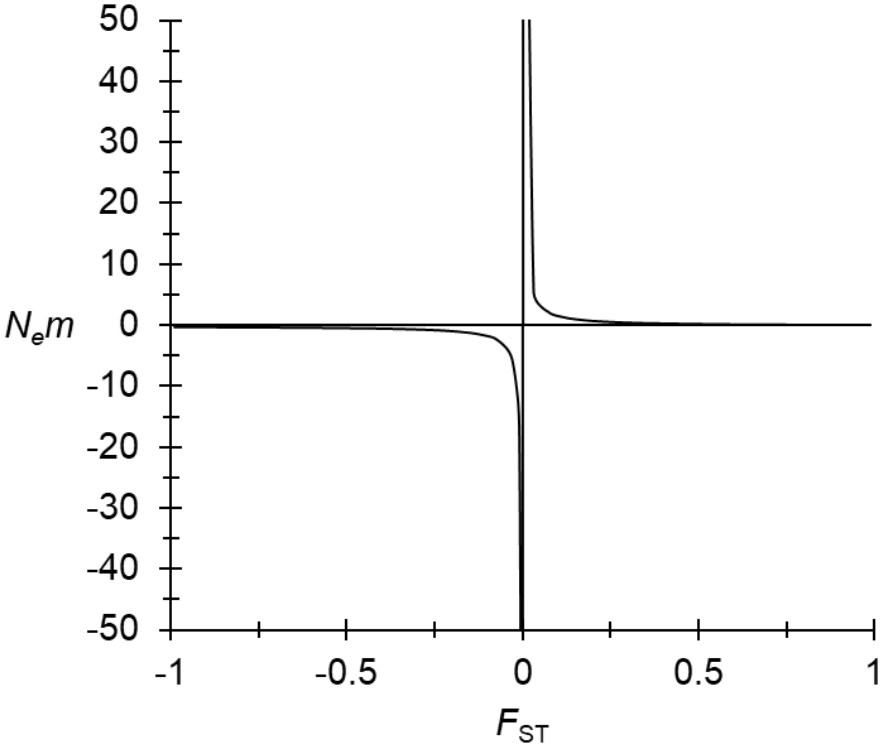
Values of number of immigrants in a three Island model as a function of *F*_ST_, at equilibrium between mutation, migration and drift.

### Appendix 3: Script for computations of geographic distances with geosphere for R

If DataLongLat1.txt contain the GPS coordinates in two columns (1^st^ Longitude, 2^nd^ Latitude), in decimal degrees for the 1^st^ subsamples of each subsample pair, and DataLongLat2.txt, the same but for the 2^nd^ subsamples, then, in R after having loaded the package geosphere and changed directory to the one wher these datasets are, geographic distances can be computed with the following commands (> means you are in the R console):

>LongLat1 <-read.table(“DataLongLat1.txt”, header=TRUE)

>LongLat2 <-read.table(“DataLongLat2.txt”, header=TRUE)

>distGeo(LongLat1,LongLat2)

Alternatively, a file named “DistGeo.txt”, containing all geographic distances in one column, can be created by typing:

>TabDgeo<-data.frame(distGeo(LongLat1,LongLat2))

>write.table(TabDgeo,”DistGeo.txt”, col=NA, sep=“\t”, dec=“.”)

## Acknowledgements

This work was initially financed by recurrent funding of the IRD. The development of the microsatellites and the sequence-based microsatellite genotyping protocol were performed at the PGTB (doi:10.15454/1.5572396583599417E12) with the help of Zoé Compagnie, Emilie Chancerel and Erwan Guichoux. This project has received funding from the European Union’s Horizon 2020 research and innovation programme under grant agreement n°101000467, acronym ‘‘COMBAT’’ (Controlling and Progressively Minimizing the Burden of Animal Trypanosomosis) (Boulangé et al., 2022). The authors wish to thank the recommender and three anonymous reviewers whose comments considerably helped improving this paper.

## Funding

This work was initially financed by recurrent funding of the IRD. This project has received funding from the European Union’s Horizon 2020 research and innovation program under grant agreement n°101000467, acronym ‘‘COMBAT’’ (Controlling and Progressively Minimizing the Burden of Animal Trypanosomosis) (Boulangé et al., 2022).

## Conflict of interest disclosure

The authors declare that they have no financial conflict of interest with the content of this article. TdM is recommender for PCI Evol Biol, PCI Ecology and PCI Infections and co-founder of PCI infections.

## Data, scripts, code, and supplementary information availability

The raw and cured datasets are available in supplementary files S1 and S2. R scripts are given in the text or in the appendix. Short reads supporting the study has been submitted to European Nucleotid Archive (European Nucleotid Archive run accession number) for shotgun whole genome sequencing used for microsatellite discovery and primer developments(ERR17897884), and for SSRseq amplicon sequencing (ERR18167661-ERR18168100).

## Notes

### Competing Interest Statement

The authors have declared no competing interest.

### Summary of Updates

Following the extended review by the recommender of PCI Zoology, to which the preprint was submitted for evaluation, we have modified the manuscript and the supplementary material accordingly.

