## Supplementary File S3 for "Development of microsatellite markers with the SSRseq method on *Glossina palpalis gambiensis*, and *G. p. palpalis*, and analysis of corresponding field samples from three sleeping sickness foci: Boffa, Dubreka (Guinea), and Bonon (Côte d’Ivoire)"

### Slide 1
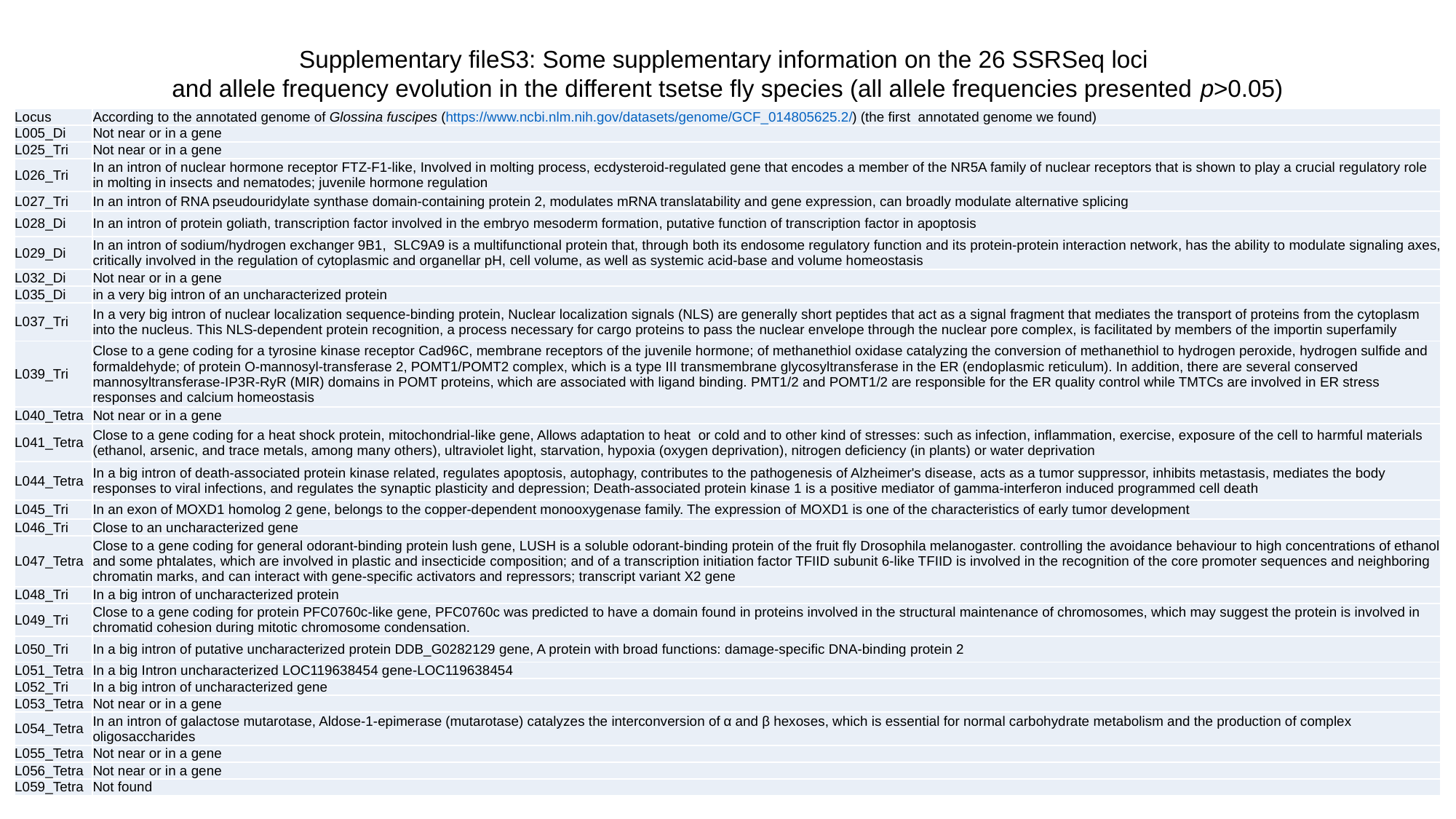

Supplementary fileS3: Some supplementary information on the 26 SSRSeq loci
and allele frequency evolution in the different tsetse fly species (all allele frequencies presented p>0.05)
| Locus | According to the annotated genome of Glossina fuscipes (https://www.ncbi.nlm.nih.gov/datasets/genome/GCF\_014805625.2/) (the first annotated genome we found) |
| --- | --- |
| L005\_Di | Not near or in a gene |
| L025\_Tri | Not near or in a gene |
| L026\_Tri | In an intron of nuclear hormone receptor FTZ-F1-like, Involved in molting process, ecdysteroid-regulated gene that encodes a member of the NR5A family of nuclear receptors that is shown to play a crucial regulatory role in molting in insects and nematodes; juvenile hormone regulation |
| L027\_Tri | In an intron of RNA pseudouridylate synthase domain-containing protein 2, modulates mRNA translatability and gene expression, can broadly modulate alternative splicing |
| L028\_Di | In an intron of protein goliath, transcription factor involved in the embryo mesoderm formation, putative function of transcription factor in apoptosis |
| L029\_Di | In an intron of sodium/hydrogen exchanger 9B1, SLC9A9 is a multifunctional protein that, through both its endosome regulatory function and its protein-protein interaction network, has the ability to modulate signaling axes, critically involved in the regulation of cytoplasmic and organellar pH, cell volume, as well as systemic acid-base and volume homeostasis |
| L032\_Di | Not near or in a gene |
| L035\_Di | in a very big intron of an uncharacterized protein |
| L037\_Tri | In a very big intron of nuclear localization sequence-binding protein, Nuclear localization signals (NLS) are generally short peptides that act as a signal fragment that mediates the transport of proteins from the cytoplasm into the nucleus. This NLS-dependent protein recognition, a process necessary for cargo proteins to pass the nuclear envelope through the nuclear pore complex, is facilitated by members of the importin superfamily |
| L039\_Tri | Close to a gene coding for a tyrosine kinase receptor Cad96C, membrane receptors of the juvenile hormone; of methanethiol oxidase catalyzing the conversion of methanethiol to hydrogen peroxide, hydrogen sulfide and formaldehyde; of protein O-mannosyl-transferase 2, POMT1/POMT2 complex, which is a type III transmembrane glycosyltransferase in the ER (endoplasmic reticulum). In addition, there are several conserved mannosyltransferase-IP3R-RyR (MIR) domains in POMT proteins, which are associated with ligand binding. PMT1/2 and POMT1/2 are responsible for the ER quality control while TMTCs are involved in ER stress responses and calcium homeostasis |
| L040\_Tetra | Not near or in a gene |
| L041\_Tetra | Close to a gene coding for a heat shock protein, mitochondrial-like gene, Allows adaptation to heat or cold and to other kind of stresses: such as infection, inflammation, exercise, exposure of the cell to harmful materials (ethanol, arsenic, and trace metals, among many others), ultraviolet light, starvation, hypoxia (oxygen deprivation), nitrogen deficiency (in plants) or water deprivation |
| L044\_Tetra | In a big intron of death-associated protein kinase related, regulates apoptosis, autophagy, contributes to the pathogenesis of Alzheimer's disease, acts as a tumor suppressor, inhibits metastasis, mediates the body responses to viral infections, and regulates the synaptic plasticity and depression; Death-associated protein kinase 1 is a positive mediator of gamma-interferon induced programmed cell death |
| L045\_Tri | In an exon of MOXD1 homolog 2 gene, belongs to the copper-dependent monooxygenase family. The expression of MOXD1 is one of the characteristics of early tumor development |
| L046\_Tri | Close to an uncharacterized gene |
| L047\_Tetra | Close to a gene coding for general odorant-binding protein lush gene, LUSH is a soluble odorant-binding protein of the fruit fly Drosophila melanogaster. controlling the avoidance behaviour to high concentrations of ethanol and some phtalates, which are involved in plastic and insecticide composition; and of a transcription initiation factor TFIID subunit 6-like TFIID is involved in the recognition of the core promoter sequences and neighboring chromatin marks, and can interact with gene-specific activators and repressors; transcript variant X2 gene |
| L048\_Tri | In a big intron of uncharacterized protein |
| L049\_Tri | Close to a gene coding for protein PFC0760c-like gene, PFC0760c was predicted to have a domain found in proteins involved in the structural maintenance of chromosomes, which may suggest the protein is involved in chromatid cohesion during mitotic chromosome condensation. |
| L050\_Tri | In a big intron of putative uncharacterized protein DDB\_G0282129 gene, A protein with broad functions: damage-specific DNA-binding protein 2 |
| L051\_Tetra | In a big Intron uncharacterized LOC119638454 gene-LOC119638454 |
| L052\_Tri | In a big intron of uncharacterized gene |
| L053\_Tetra | Not near or in a gene |
| L054\_Tetra | In an intron of galactose mutarotase, Aldose-1-epimerase (mutarotase) catalyzes the interconversion of α and β hexoses, which is essential for normal carbohydrate metabolism and the production of complex oligosaccharides |
| L055\_Tetra | Not near or in a gene |
| L056\_Tetra | Not near or in a gene |
| L059\_Tetra | Not found |

### Slide 2
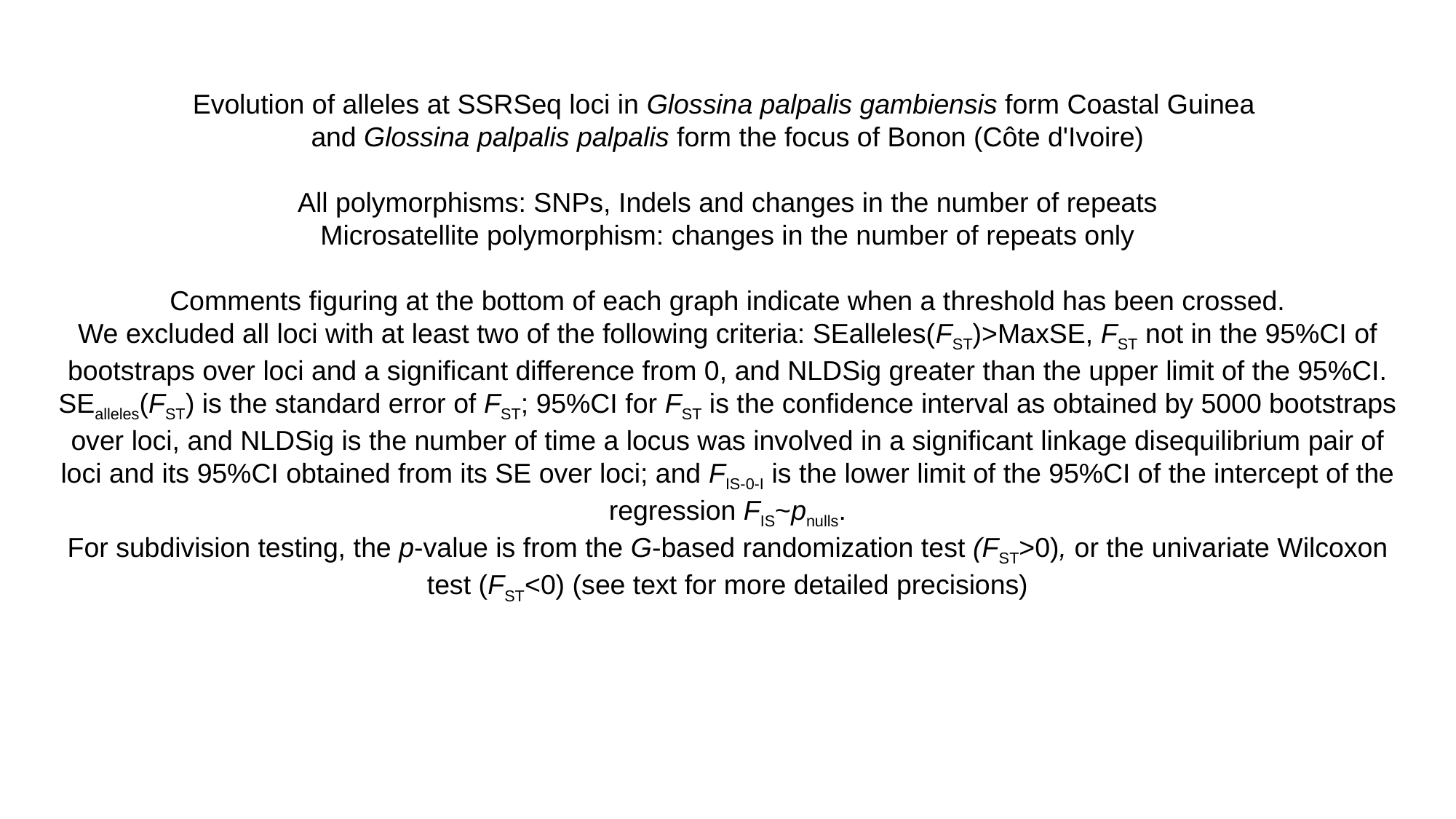

Evolution of alleles at SSRSeq loci in Glossina palpalis gambiensis form Coastal Guinea
and Glossina palpalis palpalis form the focus of Bonon (Côte d'Ivoire)
All polymorphisms: SNPs, Indels and changes in the number of repeats
Microsatellite polymorphism: changes in the number of repeats only
Comments figuring at the bottom of each graph indicate when a threshold has been crossed.
We excluded all loci with at least two of the following criteria: SEalleles(F_ST_)>MaxSE, F_ST_ not in the 95%CI of bootstraps over loci and a significant difference from 0, and NLDSig greater than the upper limit of the 95%CI.
SE_alleles_(F_ST_) is the standard error of F_ST_; 95%CI for F_ST_ is the confidence interval as obtained by 5000 bootstraps over loci, and NLDSig is the number of time a locus was involved in a significant linkage disequilibrium pair of loci and its 95%CI obtained from its SE over loci; and F_IS-0-I_ is the lower limit of the 95%CI of the intercept of the regression F_IS_~p_nulls_.
For subdivision testing, the p-value is from the G-based randomization test (F_ST_>0), or the univariate Wilcoxon test (F_ST_<0) (see text for more detailed precisions)

### Slide 3
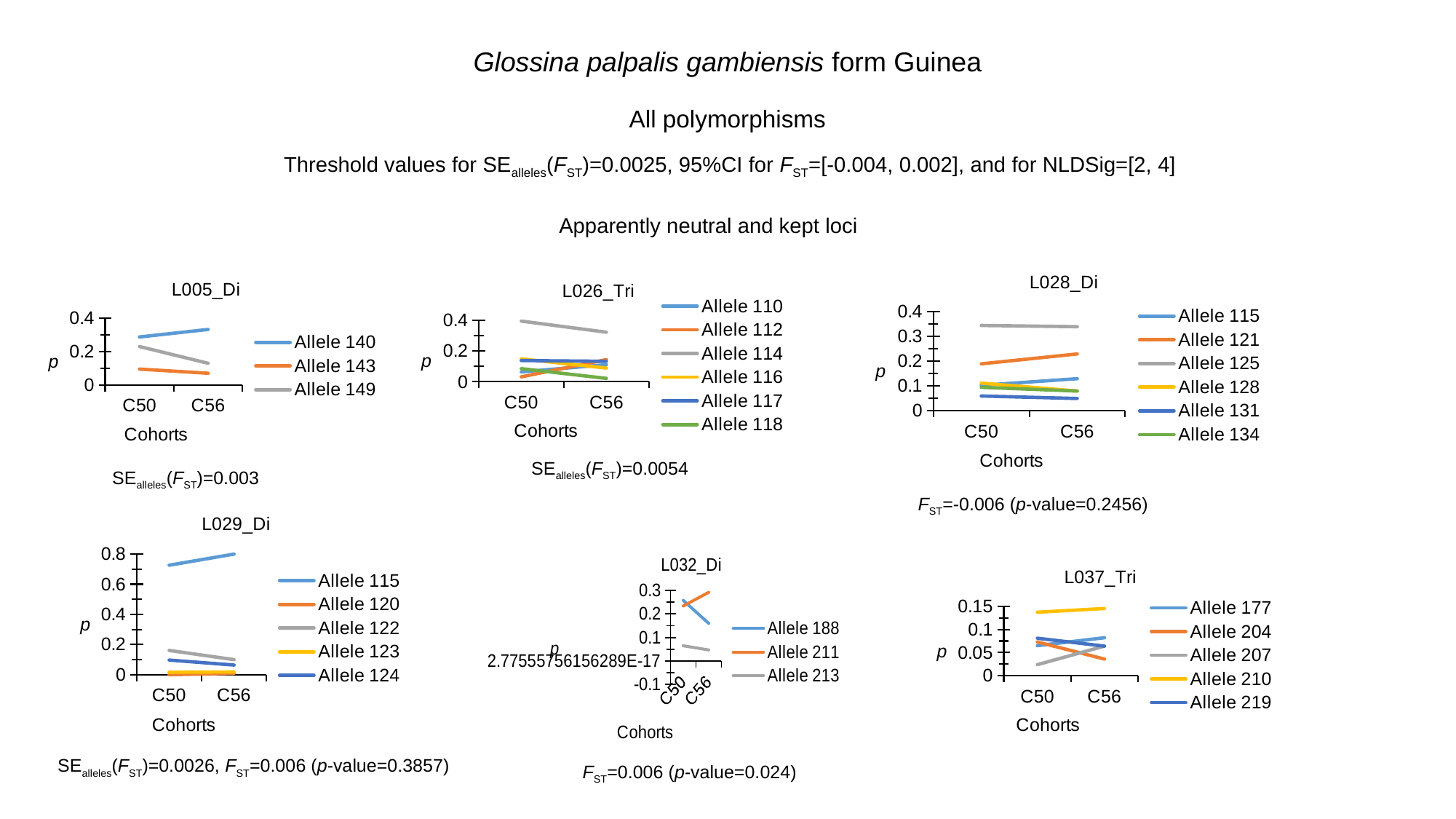

Glossina palpalis gambiensis form Guinea
All polymorphisms
Threshold values for SE_alleles_(F_ST_)=0.0025, 95%CI for F_ST_=[-0.004, 0.002], and for NLDSig=[2, 4]
Apparently neutral and kept loci
#### Chart: L028_Di
| Category | Allele 115 | Allele 121 | Allele 125 | Allele 128 | Allele 131 | Allele 134 |
|---|---|---|---|---|---|---|
| C50 | 0.103 | 0.19 | 0.345 | 0.112 | 0.06 | 0.095 |
| C56 | 0.13 | 0.23 | 0.34 | 0.08 | 0.05 | 0.08 |
#### Chart: L005_Di
| Category | Allele 140 | Allele 143 | Allele 149 |
|---|---|---|---|
| C50 | 0.288 | 0.096 | 0.231 |
| C56 | 0.333 | 0.071 | 0.131 |
#### Chart: L026_Tri
| Category | Allele 110 | Allele 112 | Allele 114 | Allele 116 | Allele 117 | Allele 118 |
|---|---|---|---|---|---|---|
| C50 | 0.064 | 0.032 | 0.394 | 0.149 | 0.138 | 0.085 |
| C56 | 0.111 | 0.144 | 0.322 | 0.089 | 0.133 | 0.022 |SE_alleles_(F_ST_)=0.0054
SE_alleles_(F_ST_)=0.003
F_ST_=-0.006 (p-value=0.2456)
#### Chart: L029_Di
| Category | Allele 115 | Allele 120 | Allele 122 | Allele 123 | Allele 124 |
|---|---|---|---|---|---|
| C50 | 0.726 | 0.0 | 0.161 | 0.016 | 0.097 |
| C56 | 0.8 | 0.009 | 0.1 | 0.018 | 0.064 |
#### Chart: L032_Di
| Category | Allele 188 | Allele 211 | Allele 213 |
|---|---|---|---|
| C50 | 0.258 | 0.234 | 0.065 |
| C56 | 0.16 | 0.292 | 0.047 |
#### Chart: L037_Tri
| Category | Allele 177 | Allele 204 | Allele 207 | Allele 210 | Allele 219 |
|---|---|---|---|---|---|
| C50 | 0.065 | 0.073 | 0.024 | 0.137 | 0.081 |
| C56 | 0.082 | 0.036 | 0.064 | 0.145 | 0.064 |SE_alleles_(F_ST_)=0.0026, F_ST_=0.006 (p-value=0.3857)
F_ST_=0.006 (p-value=0.024)

### Slide 4
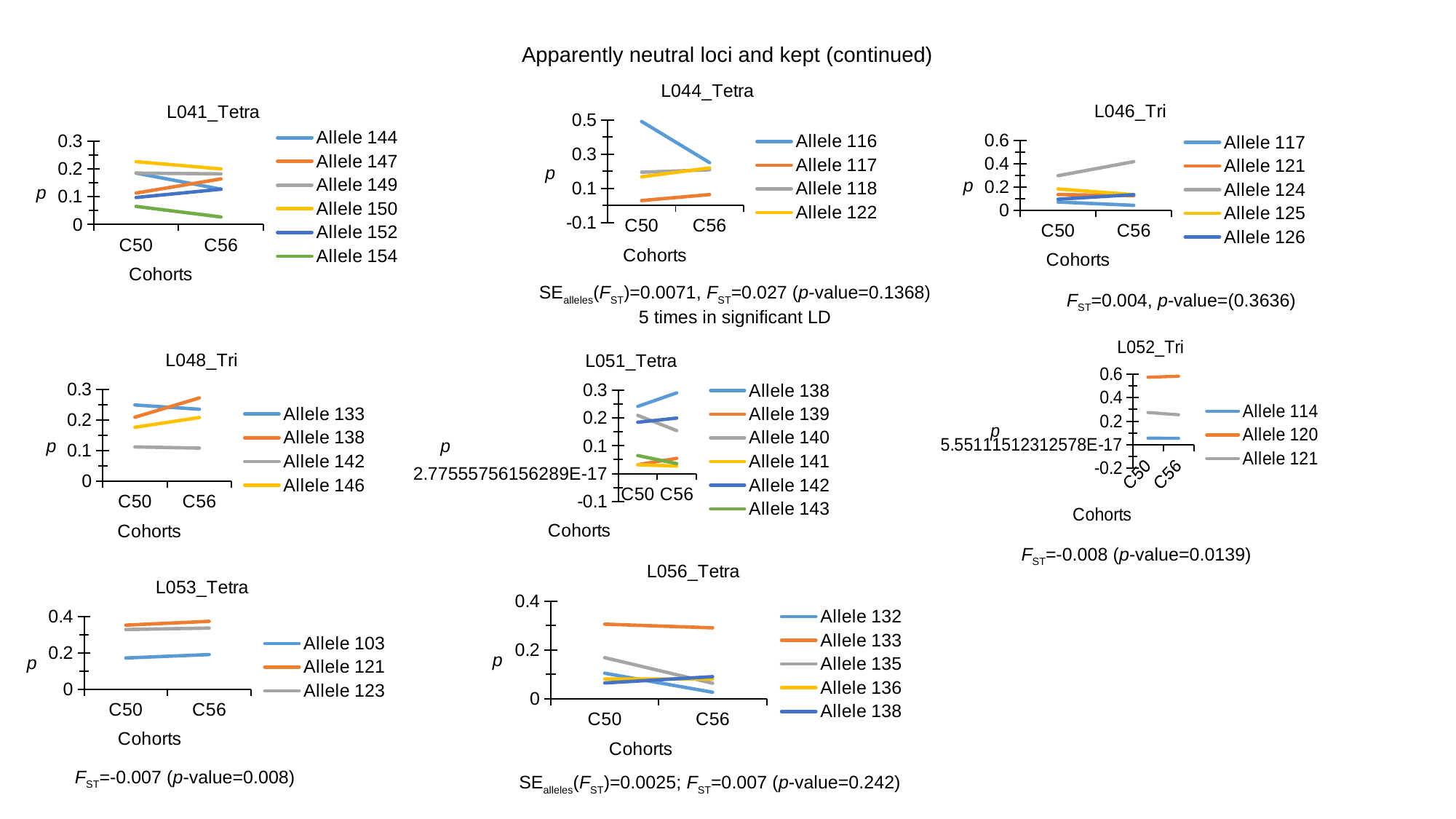

Apparently neutral loci and kept (continued)
#### Chart: L044_Tetra
| Category | Allele 116 | Allele 117 | Allele 118 | Allele 122 |
|---|---|---|---|---|
| C50 | 0.491 | 0.028 | 0.194 | 0.167 |
| C56 | 0.25 | 0.063 | 0.208 | 0.219 |
#### Chart: L046_Tri
| Category | Allele 117 | Allele 121 | Allele 124 | Allele 125 | Allele 126 |
|---|---|---|---|---|---|
| C50 | 0.073 | 0.137 | 0.298 | 0.185 | 0.097 |
| C56 | 0.045 | 0.127 | 0.418 | 0.136 | 0.136 |
#### Chart: L041_Tetra
| Category | Allele 144 | Allele 147 | Allele 149 | Allele 150 | Allele 152 | Allele 154 |
|---|---|---|---|---|---|---|
| C50 | 0.185 | 0.113 | 0.185 | 0.226 | 0.097 | 0.065 |
| C56 | 0.127 | 0.164 | 0.182 | 0.2 | 0.127 | 0.027 |SE_alleles_(F_ST_)=0.0071, F_ST_=0.027 (p-value=0.1368)
5 times in significant LD
F_ST_=0.004, p-value=(0.3636)
#### Chart: L052_Tri
| Category | Allele 114 | Allele 120 | Allele 121 |
|---|---|---|---|
| C50 | 0.056 | 0.573 | 0.274 |
| C56 | 0.055 | 0.582 | 0.255 |
#### Chart: L048_Tri
| Category | Allele 133 | Allele 138 | Allele 142 | Allele 146 |
|---|---|---|---|---|
| C50 | 0.25 | 0.21 | 0.113 | 0.177 |
| C56 | 0.236 | 0.273 | 0.109 | 0.209 |
#### Chart: L051_Tetra
| Category | Allele 138 | Allele 139 | Allele 140 | Allele 141 | Allele 142 | Allele 143 |
|---|---|---|---|---|---|---|
| C50 | 0.242 | 0.032 | 0.21 | 0.032 | 0.185 | 0.065 |
| C56 | 0.291 | 0.055 | 0.155 | 0.027 | 0.2 | 0.036 |F_ST_=-0.008 (p-value=0.0139)
#### Chart: L056_Tetra
| Category | Allele 132 | Allele 133 | Allele 135 | Allele 136 | Allele 138 |
|---|---|---|---|---|---|
| C50 | 0.105 | 0.306 | 0.169 | 0.081 | 0.065 |
| C56 | 0.027 | 0.291 | 0.064 | 0.082 | 0.091 |
#### Chart: L053_Tetra
| Category | Allele 103 | Allele 121 | Allele 123 |
|---|---|---|---|
| C50 | 0.172 | 0.352 | 0.328 |
| C56 | 0.191 | 0.373 | 0.336 |F_ST_=-0.007 (p-value=0.008)
SE_alleles_(F_ST_)=0.0025; F_ST_=0.007 (p-value=0.242)

### Slide 5
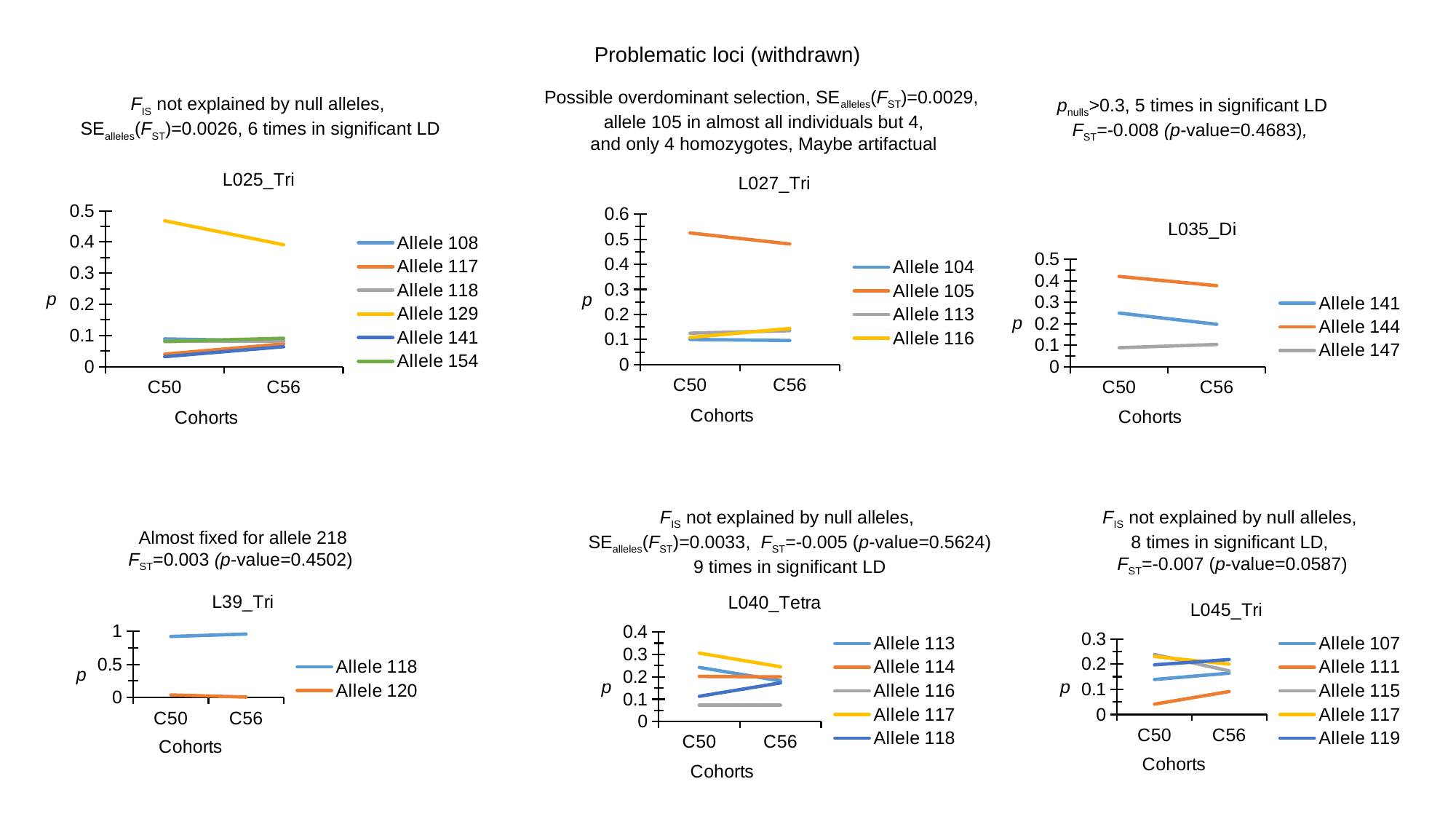

Problematic loci (withdrawn)
Possible overdominant selection, SE_alleles_(F_ST_)=0.0029,
allele 105 in almost all individuals but 4,
and only 4 homozygotes, Maybe artifactual
F_IS_ not explained by null alleles,
SE_alleles_(F_ST_)=0.0026, 6 times in significant LD
p_nulls_>0.3, 5 times in significant LD
F_ST_=-0.008 (p-value=0.4683),
#### Chart: L025_Tri
| Category | Allele 108 | Allele 117 | Allele 118 | Allele 129 | Allele 141 | Allele 154 |
|---|---|---|---|---|---|---|
| C50 | 0.089 | 0.04 | 0.081 | 0.468 | 0.032 | 0.081 |
| C56 | 0.082 | 0.073 | 0.082 | 0.391 | 0.064 | 0.091 |
#### Chart: L027_Tri
| Category | Allele 104 | Allele 105 | Allele 113 | Allele 116 |
|---|---|---|---|---|
| C50 | 0.1 | 0.525 | 0.125 | 0.108 |
| C56 | 0.096 | 0.481 | 0.135 | 0.144 |
#### Chart: L035_Di
| Category | Allele 141 | Allele 144 | Allele 147 |
|---|---|---|---|
| C50 | 0.25 | 0.42 | 0.089 |
| C56 | 0.198 | 0.377 | 0.104 |F_IS_ not explained by null alleles,
8 times in significant LD,
F_ST_=-0.007 (p-value=0.0587)
F_IS_ not explained by null alleles,
SE_alleles_(F_ST_)=0.0033, F_ST_=-0.005 (p-value=0.5624)
9 times in significant LD
Almost fixed for allele 218
F_ST_=0.003 (p-value=0.4502)
#### Chart: L39_Tri
| Category | Allele 118 | Allele 120 |
|---|---|---|
| C50 | 0.919 | 0.04 |
| C56 | 0.955 | 0.009 |
#### Chart: L040_Tetra
| Category | Allele 113 | Allele 114 | Allele 116 | Allele 117 | Allele 118 |
|---|---|---|---|---|---|
| C50 | 0.242 | 0.202 | 0.073 | 0.306 | 0.113 |
| C56 | 0.182 | 0.2 | 0.073 | 0.245 | 0.173 |
#### Chart: L045_Tri
| Category | Allele 107 | Allele 111 | Allele 115 | Allele 117 | Allele 119 |
|---|---|---|---|---|---|
| C50 | 0.139 | 0.041 | 0.238 | 0.23 | 0.197 |
| C56 | 0.164 | 0.091 | 0.173 | 0.2 | 0.218 |

### Slide 6
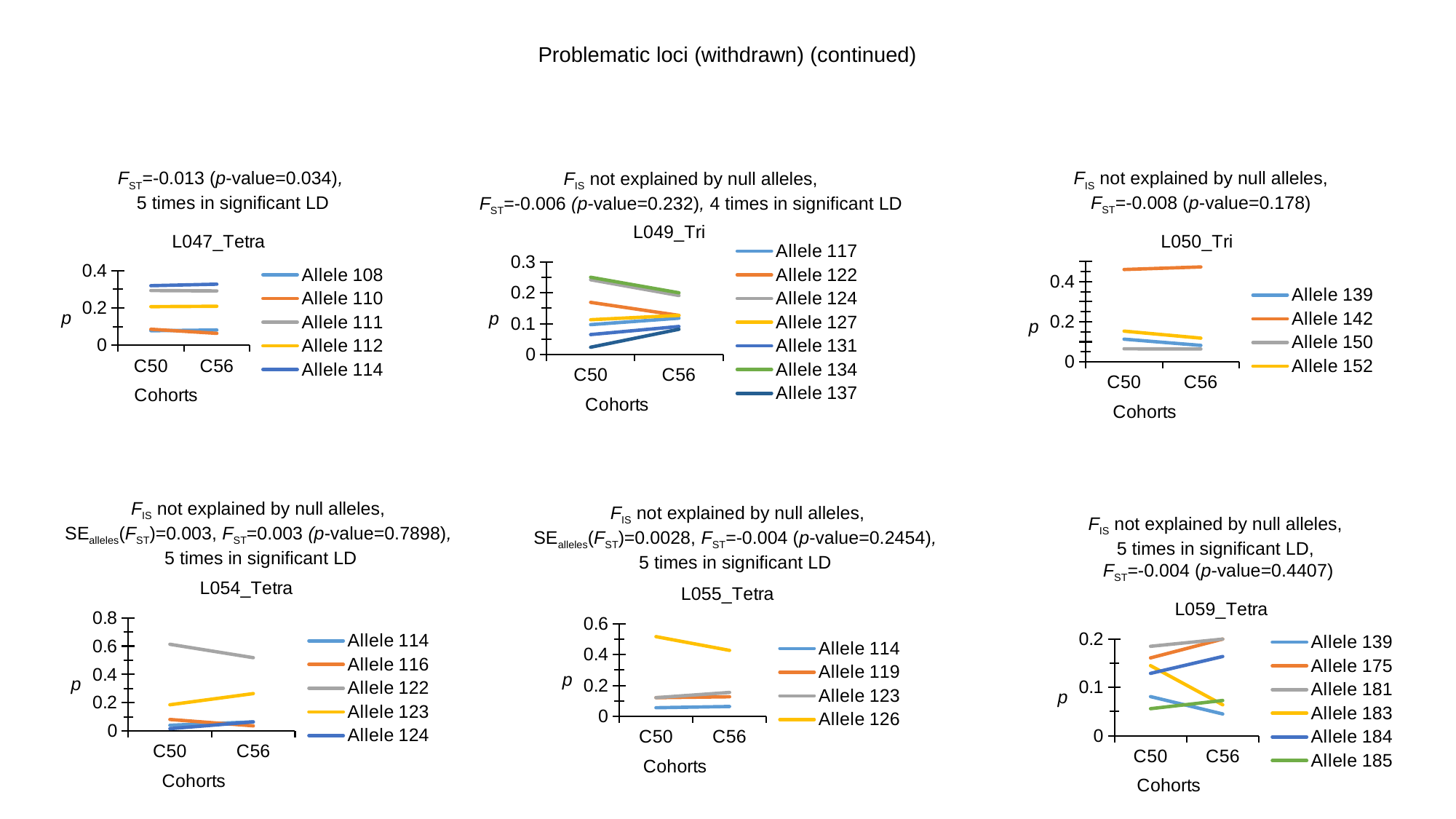

Problematic loci (withdrawn) (continued)
F_ST_=-0.013 (p-value=0.034),
5 times in significant LD
F_IS_ not explained by null alleles,
F_ST_=-0.008 (p-value=0.178)
F_IS_ not explained by null alleles,
F_ST_=-0.006 (p-value=0.232), 4 times in significant LD
#### Chart: L049_Tri
| Category | Allele 117 | Allele 122 | Allele 124 | Allele 127 | Allele 131 | Allele 134 | Allele 137 |
|---|---|---|---|---|---|---|---|
| C50 | 0.097 | 0.169 | 0.242 | 0.113 | 0.065 | 0.25 | 0.024 |
| C56 | 0.118 | 0.127 | 0.191 | 0.127 | 0.091 | 0.2 | 0.082 |
#### Chart: L047_Tetra
| Category | Allele 108 | Allele 110 | Allele 111 | Allele 112 | Allele 114 |
|---|---|---|---|---|---|
| C50 | 0.078 | 0.086 | 0.293 | 0.207 | 0.319 |
| C56 | 0.082 | 0.064 | 0.291 | 0.209 | 0.327 |
#### Chart: L050_Tri
| Category | Allele 139 | Allele 142 | Allele 150 | Allele 152 |
|---|---|---|---|---|
| C50 | 0.113 | 0.46 | 0.065 | 0.153 |
| C56 | 0.082 | 0.473 | 0.064 | 0.118 |F_IS_ not explained by null alleles,
SE_alleles_(F_ST_)=0.003, F_ST_=0.003 (p-value=0.7898),
5 times in significant LD
F_IS_ not explained by null alleles,
SE_alleles_(F_ST_)=0.0028, F_ST_=-0.004 (p-value=0.2454),
5 times in significant LD
F_IS_ not explained by null alleles,
5 times in significant LD,
F_ST_=-0.004 (p-value=0.4407)
#### Chart: L054_Tetra
| Category | Allele 114 | Allele 116 | Allele 122 | Allele 123 | Allele 124 |
|---|---|---|---|---|---|
| C50 | 0.04 | 0.081 | 0.613 | 0.185 | 0.016 |
| C56 | 0.064 | 0.036 | 0.518 | 0.264 | 0.064 |
#### Chart: L055_Tetra
| Category | Allele 114 | Allele 119 | Allele 123 | Allele 126 |
|---|---|---|---|---|
| C50 | 0.056 | 0.121 | 0.121 | 0.516 |
| C56 | 0.064 | 0.127 | 0.155 | 0.427 |
#### Chart: L059_Tetra
| Category | Allele 139 | Allele 175 | Allele 181 | Allele 183 | Allele 184 | Allele 185 |
|---|---|---|---|---|---|---|
| C50 | 0.081 | 0.161 | 0.185 | 0.145 | 0.129 | 0.056 |
| C56 | 0.045 | 0.2 | 0.2 | 0.064 | 0.164 | 0.073 |

### Slide 7
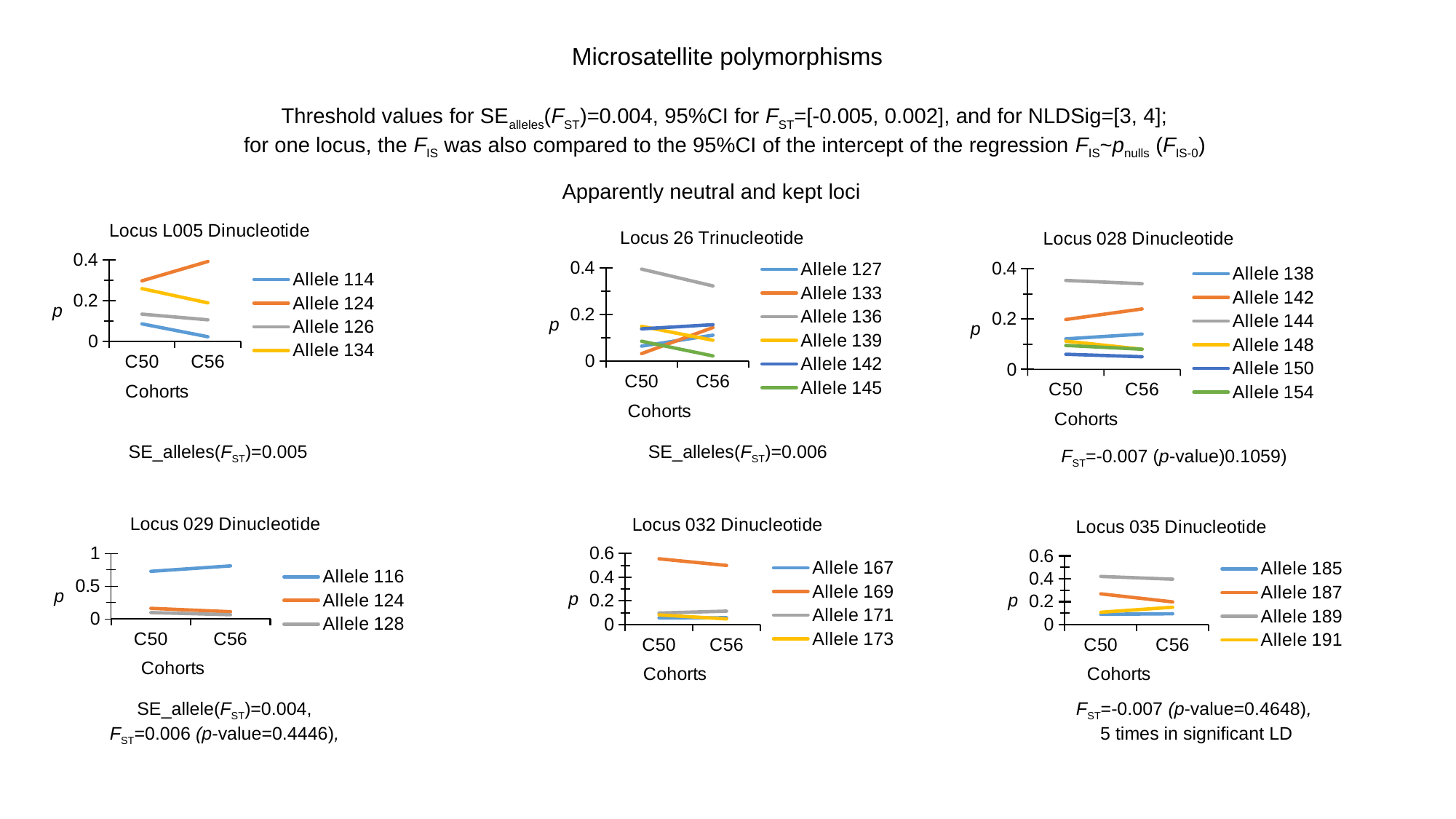

Microsatellite polymorphisms
Threshold values for SE_alleles_(F_ST_)=0.004, 95%CI for F_ST_=[-0.005, 0.002], and for NLDSig=[3, 4];
for one locus, the F_IS_ was also compared to the 95%CI of the intercept of the regression F_IS_~p_nulls_ (F_IS-0_)
Apparently neutral and kept loci
#### Chart: Locus L005 Dinucleotide
| Category | Allele 114 | Allele 124 | Allele 126 | Allele 134 |
|---|---|---|---|---|
| C50 | 0.087 | 0.298 | 0.135 | 0.26 |
| C56 | 0.024 | 0.393 | 0.107 | 0.19 |
#### Chart: Locus 26 Trinucleotide
| Category | Allele 127 | Allele 133 | Allele 136 | Allele 139 | Allele 142 | Allele 145 |
|---|---|---|---|---|---|---|
| C50 | 0.064 | 0.032 | 0.394 | 0.149 | 0.138 | 0.085 |
| C56 | 0.111 | 0.144 | 0.322 | 0.089 | 0.156 | 0.022 |
#### Chart: Locus 028 Dinucleotide
| Category | Allele 138 | Allele 142 | Allele 144 | Allele 148 | Allele 150 | Allele 154 |
|---|---|---|---|---|---|---|
| C50 | 0.121 | 0.198 | 0.353 | 0.112 | 0.06 | 0.095 |
| C56 | 0.14 | 0.24 | 0.34 | 0.08 | 0.05 | 0.08 |SE_alleles(F_ST_)=0.005
SE_alleles(F_ST_)=0.006
F_ST_=-0.007 (p-value)0.1059)
#### Chart: Locus 029 Dinucleotide
| Category | Allele 116 | Allele 124 | Allele 128 |
|---|---|---|---|
| C50 | 0.726 | 0.161 | 0.097 |
| C56 | 0.809 | 0.109 | 0.064 |
#### Chart: Locus 032 Dinucleotide
| Category | Allele 167 | Allele 169 | Allele 171 | Allele 173 |
|---|---|---|---|---|
| C50 | 0.056 | 0.556 | 0.097 | 0.081 |
| C56 | 0.057 | 0.5 | 0.113 | 0.047 |
#### Chart: Locus 035 Dinucleotide
| Category | Allele 185 | Allele 187 | Allele 189 | Allele 191 |
|---|---|---|---|---|
| C50 | 0.089 | 0.268 | 0.42 | 0.107 |
| C56 | 0.094 | 0.198 | 0.396 | 0.151 |SE_allele(F_ST_)=0.004,
F_ST_=0.006 (p-value=0.4446),
F_ST_=-0.007 (p-value=0.4648),
5 times in significant LD

### Slide 8
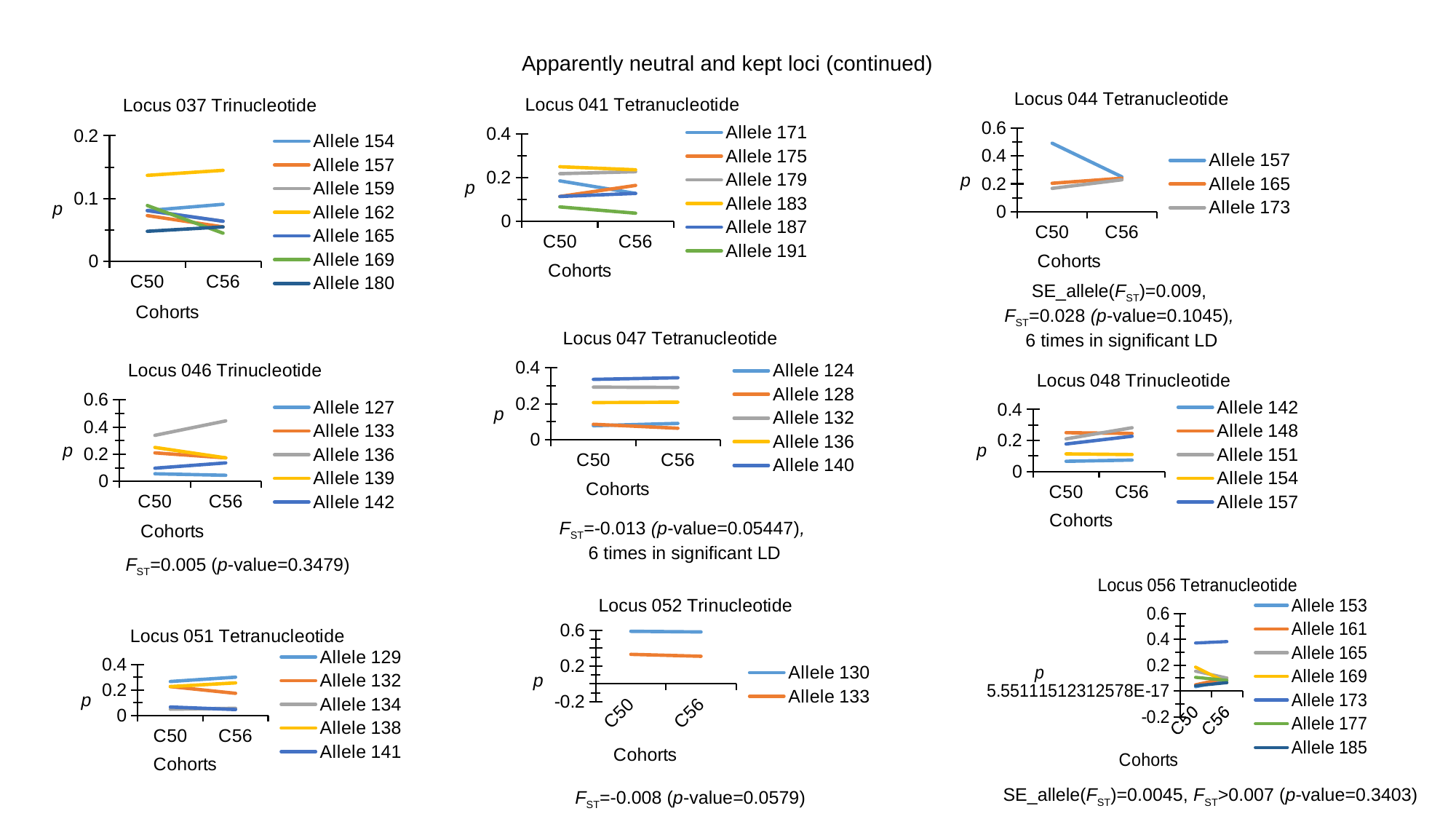

Apparently neutral and kept loci (continued)
#### Chart: Locus 044 Tetranucleotide
| Category | Allele 157 | Allele 165 | Allele 173 |
|---|---|---|---|
| C50 | 0.491 | 0.204 | 0.167 |
| C56 | 0.25 | 0.24 | 0.229 |
#### Chart: Locus 037 Trinucleotide
| Category | Allele 154 | Allele 157 | Allele 159 | Allele 162 | Allele 165 | Allele 169 | Allele 180 |
|---|---|---|---|---|---|---|---|
| C50 | 0.081 | 0.073 | 0.081 | 0.137 | 0.081 | 0.089 | 0.048 |
| C56 | 0.091 | 0.055 | 0.064 | 0.145 | 0.064 | 0.045 | 0.055 |
#### Chart: Locus 041 Tetranucleotide
| Category | Allele 171 | Allele 175 | Allele 179 | Allele 183 | Allele 187 | Allele 191 |
|---|---|---|---|---|---|---|
| C50 | 0.185 | 0.113 | 0.218 | 0.25 | 0.113 | 0.065 |
| C56 | 0.127 | 0.164 | 0.227 | 0.236 | 0.127 | 0.036 |SE_allele(F_ST_)=0.009,
F_ST_=0.028 (p-value=0.1045),
6 times in significant LD
#### Chart: Locus 047 Tetranucleotide
| Category | Allele 124 | Allele 128 | Allele 132 | Allele 136 | Allele 140 |
|---|---|---|---|---|---|
| C50 | 0.078 | 0.086 | 0.293 | 0.207 | 0.336 |
| C56 | 0.091 | 0.064 | 0.291 | 0.209 | 0.345 |
#### Chart: Locus 046 Trinucleotide
| Category | Allele 127 | Allele 133 | Allele 136 | Allele 139 | Allele 142 |
|---|---|---|---|---|---|
| C50 | 0.056 | 0.21 | 0.339 | 0.25 | 0.097 |
| C56 | 0.045 | 0.173 | 0.445 | 0.173 | 0.136 |
#### Chart: Locus 048 Trinucleotide
| Category | Allele 142 | Allele 148 | Allele 151 | Allele 154 | Allele 157 |
|---|---|---|---|---|---|
| C50 | 0.065 | 0.25 | 0.21 | 0.113 | 0.177 |
| C56 | 0.073 | 0.245 | 0.282 | 0.109 | 0.227 |F_ST_=-0.013 (p-value=0.05447),
6 times in significant LD
F_ST_=0.005 (p-value=0.3479)
#### Chart: Locus 056 Tetranucleotide
| Category | Allele 153 | Allele 161 | Allele 165 | Allele 169 | Allele 173 | Allele 177 | Allele 185 |
|---|---|---|---|---|---|---|---|
| C50 | 0.032 | 0.048 | 0.153 | 0.185 | 0.371 | 0.105 | 0.04 |
| C56 | 0.082 | 0.1 | 0.1 | 0.064 | 0.382 | 0.082 | 0.064 |
#### Chart: Locus 052 Trinucleotide
| Category | Allele 130 | Allele 133 |
|---|---|---|
| C50 | 0.589 | 0.331 |
| C56 | 0.582 | 0.309 |
#### Chart: Locus 051 Tetranucleotide
| Category | Allele 129 | Allele 132 | Allele 134 | Allele 138 | Allele 141 |
|---|---|---|---|---|---|
| C50 | 0.266 | 0.226 | 0.048 | 0.226 | 0.065 |
| C56 | 0.3 | 0.173 | 0.055 | 0.255 | 0.045 |SE_allele(F_ST_)=0.0045, F_ST_>0.007 (p-value=0.3403)
F_ST_=-0.008 (p-value=0.0579)

### Slide 9
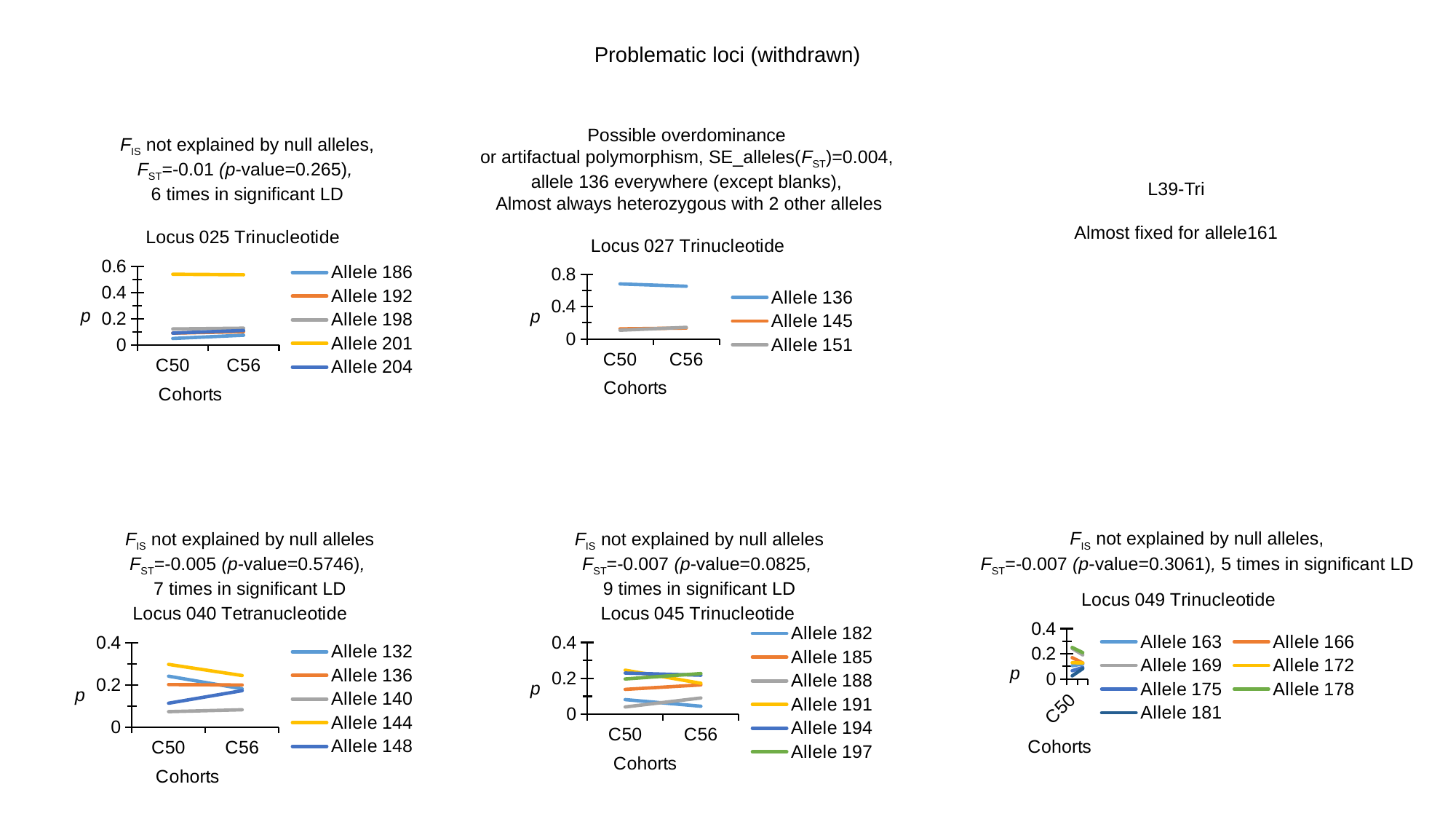

Problematic loci (withdrawn)
Possible overdominance
or artifactual polymorphism, SE_alleles(F_ST_)=0.004,
allele 136 everywhere (except blanks),
Almost always heterozygous with 2 other alleles
F_IS_ not explained by null alleles,
F_ST_=-0.01 (p-value=0.265),
6 times in significant LD
L39-Tri
Almost fixed for allele161
#### Chart: Locus 025 Trinucleotide
| Category | Allele 186 | Allele 192 | Allele 198 | Allele 201 | Allele 204 |
|---|---|---|---|---|---|
| C50 | 0.048 | 0.089 | 0.121 | 0.54 | 0.089 |
| C56 | 0.073 | 0.1 | 0.127 | 0.536 | 0.109 |
#### Chart: Locus 027 Trinucleotide
| Category | Allele 136 | Allele 145 | Allele 151 |
|---|---|---|---|
| C50 | 0.683 | 0.125 | 0.108 |
| C56 | 0.654 | 0.135 | 0.144 |F_IS_ not explained by null alleles
F_ST_=-0.005 (p-value=0.5746),
7 times in significant LD
F_IS_ not explained by null alleles,
F_ST_=-0.007 (p-value=0.3061), 5 times in significant LD
F_IS_ not explained by null alleles
F_ST_=-0.007 (p-value=0.0825,
9 times in significant LD
#### Chart: Locus 049 Trinucleotide
| Category | Allele 163 | Allele 166 | Allele 169 | Allele 172 | Allele 175 | Allele 178 | Allele 181 |
|---|---|---|---|---|---|---|---|
| C50 | 0.105 | 0.169 | 0.242 | 0.129 | 0.065 | 0.25 | 0.024 |
| C56 | 0.118 | 0.127 | 0.191 | 0.127 | 0.091 | 0.209 | 0.082 |
#### Chart: Locus 040 Tetranucleotide
| Category | Allele 132 | Allele 136 | Allele 140 | Allele 144 | Allele 148 |
|---|---|---|---|---|---|
| C50 | 0.242 | 0.202 | 0.073 | 0.298 | 0.113 |
| C56 | 0.182 | 0.2 | 0.082 | 0.245 | 0.173 |
#### Chart: Locus 045 Trinucleotide
| Category | Allele 182 | Allele 185 | Allele 188 | Allele 191 | Allele 194 | Allele 197 |
|---|---|---|---|---|---|---|
| C50 | 0.082 | 0.139 | 0.041 | 0.246 | 0.23 | 0.197 |
| C56 | 0.045 | 0.164 | 0.091 | 0.173 | 0.218 | 0.227 |

### Slide 10
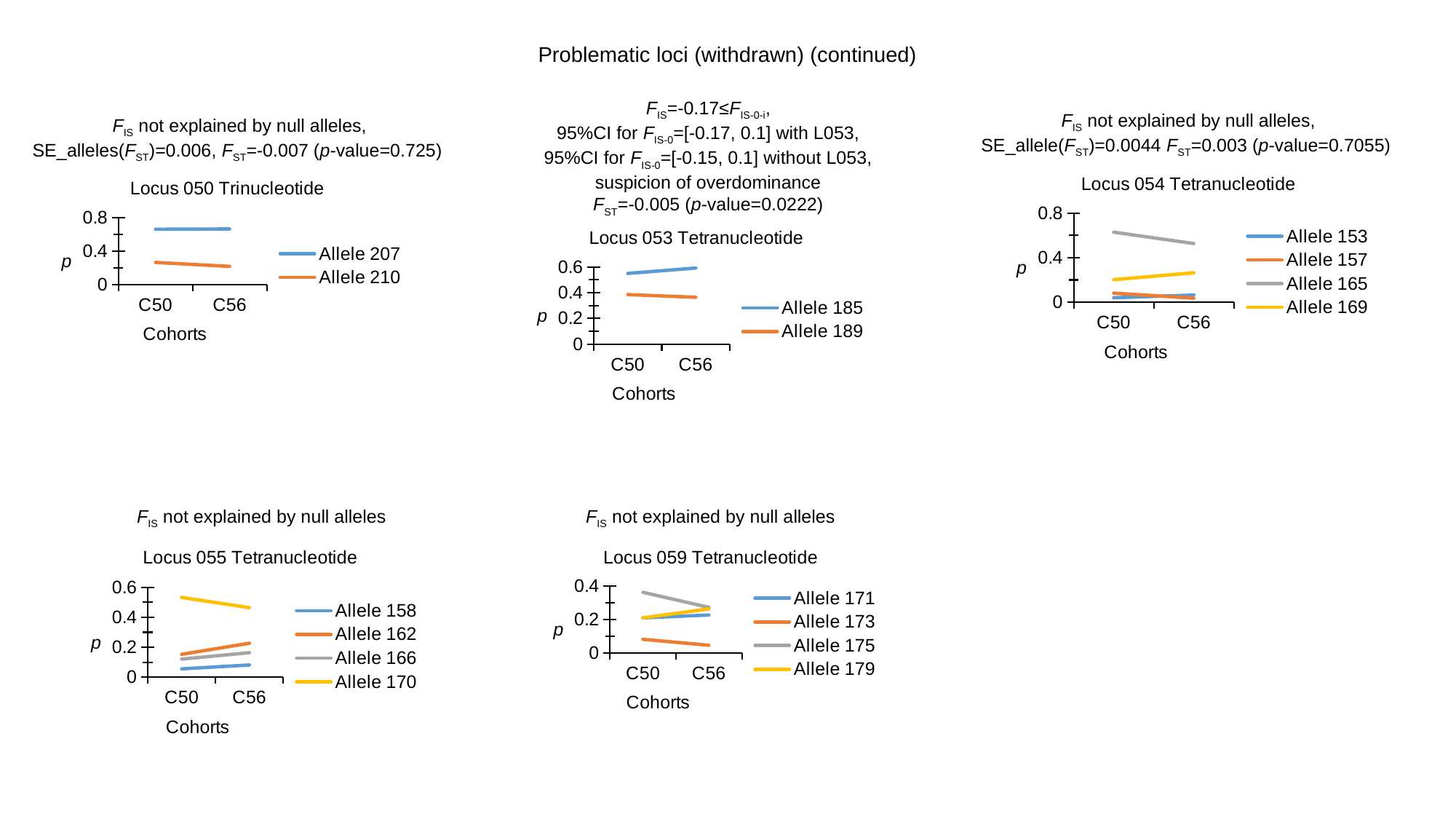

Problematic loci (withdrawn) (continued)
F_IS_=-0.17≤F_IS-0-i_,
95%CI for F_IS-0_=[-0.17, 0.1] with L053,
95%CI for F_IS-0_=[-0.15, 0.1] without L053,
suspicion of overdominance
F_ST_=-0.005 (p-value=0.0222)
F_IS_ not explained by null alleles,
SE_allele(F_ST_)=0.0044 F_ST_=0.003 (p-value=0.7055)
F_IS_ not explained by null alleles,
SE_alleles(F_ST_)=0.006, F_ST_=-0.007 (p-value=0.725)
#### Chart: Locus 054 Tetranucleotide
| Category | Allele 153 | Allele 157 | Allele 165 | Allele 169 |
|---|---|---|---|---|
| C50 | 0.04 | 0.081 | 0.629 | 0.202 |
| C56 | 0.064 | 0.036 | 0.527 | 0.264 |
#### Chart: Locus 050 Trinucleotide
| Category | Allele 207 | Allele 210 |
|---|---|---|
| C50 | 0.661 | 0.266 |
| C56 | 0.664 | 0.218 |
#### Chart: Locus 053 Tetranucleotide
| Category | Allele 185 | Allele 189 |
|---|---|---|
| C50 | 0.549 | 0.385 |
| C56 | 0.591 | 0.364 |F_IS_ not explained by null alleles
F_IS_ not explained by null alleles
#### Chart: Locus 055 Tetranucleotide
| Category | Allele 158 | Allele 162 | Allele 166 | Allele 170 |
|---|---|---|---|---|
| C50 | 0.056 | 0.153 | 0.121 | 0.532 |
| C56 | 0.082 | 0.227 | 0.164 | 0.464 |
#### Chart: Locus 059 Tetranucleotide
| Category | Allele 171 | Allele 173 | Allele 175 | Allele 179 |
|---|---|---|---|---|
| C50 | 0.21 | 0.081 | 0.363 | 0.21 |
| C56 | 0.227 | 0.045 | 0.273 | 0.264 |

### Slide 11
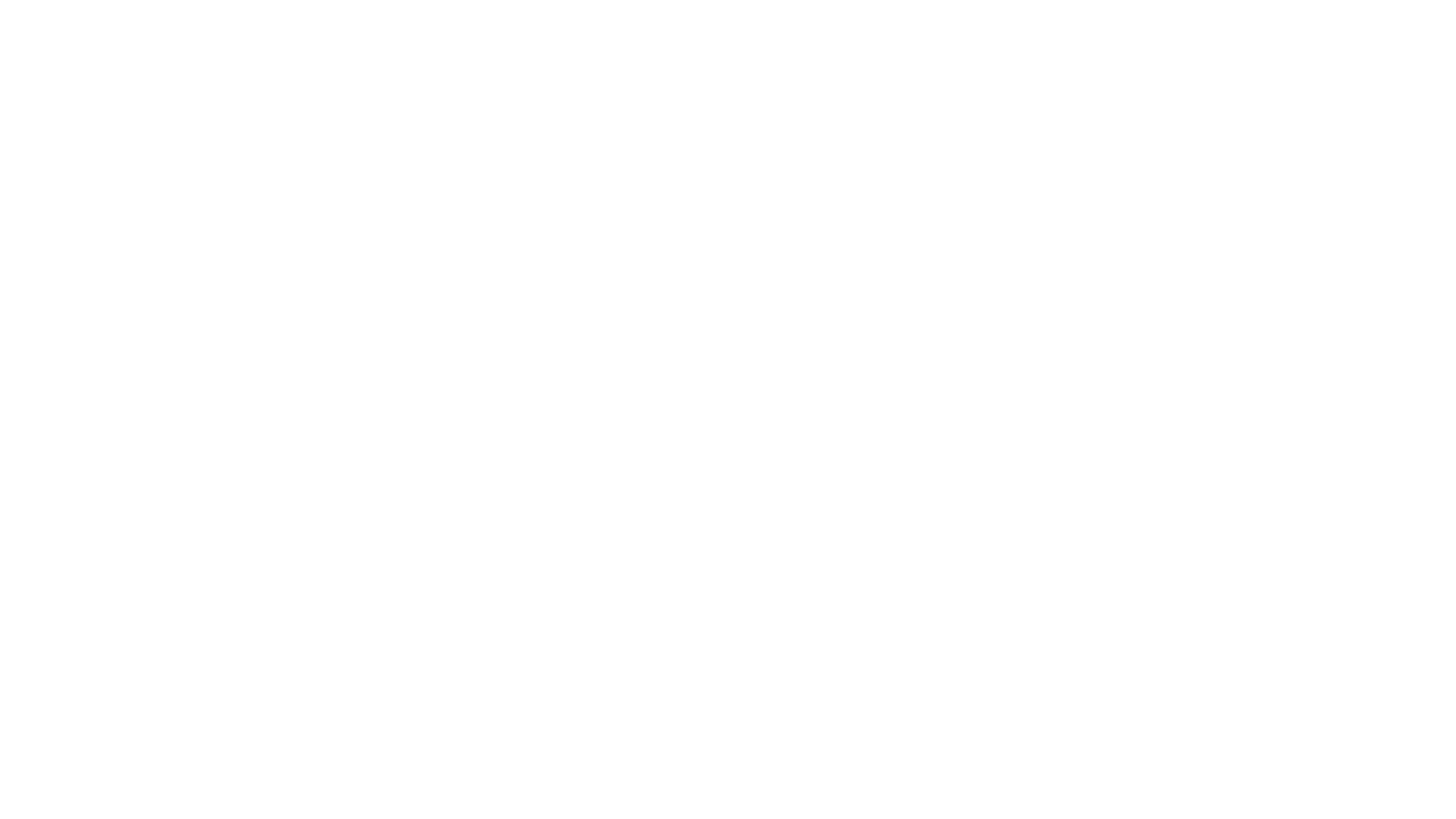

### Slide 12
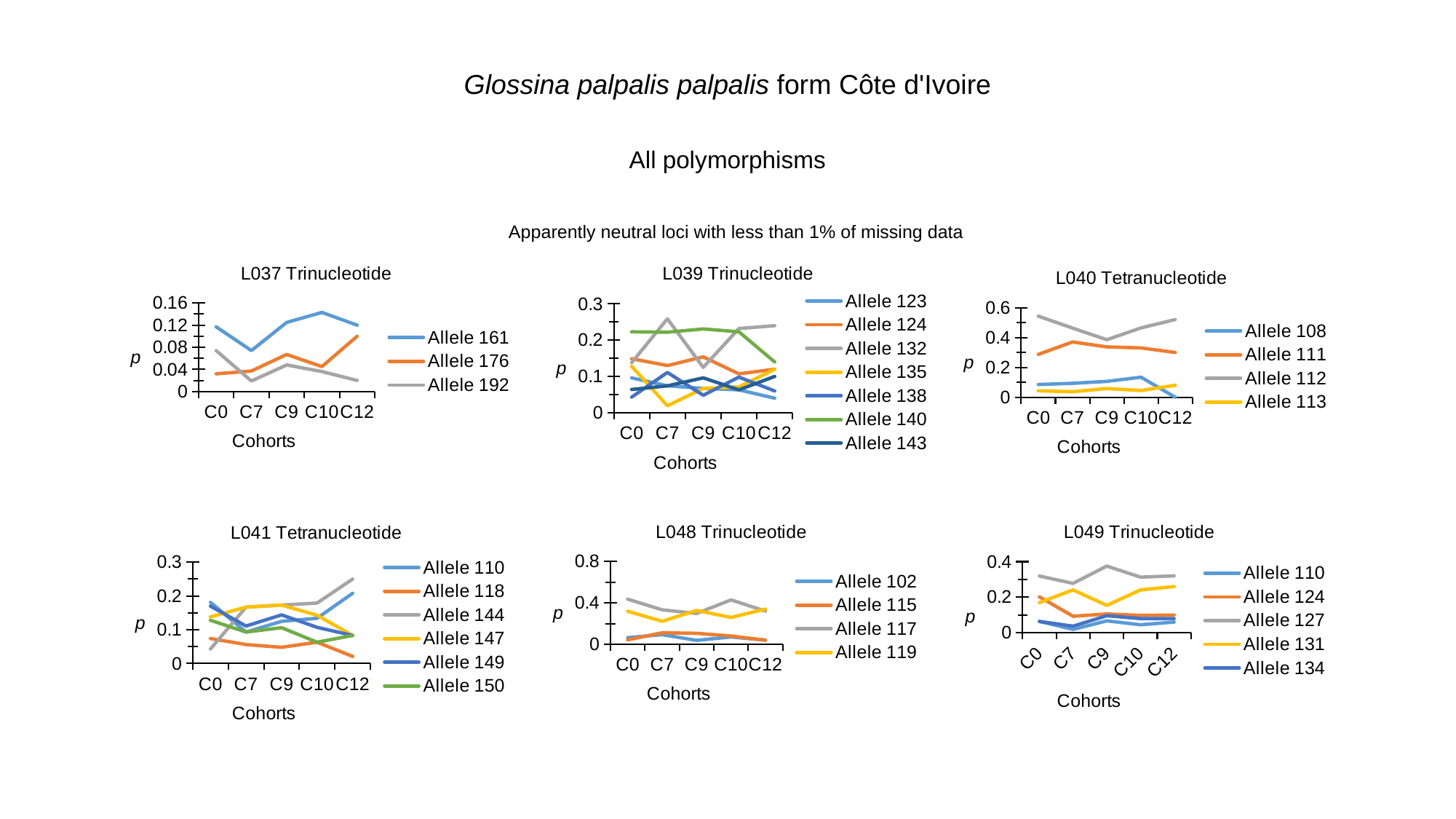

Glossina palpalis palpalis form Côte d'Ivoire
All polymorphisms
Apparently neutral loci with less than 1% of missing data
#### Chart: L037 Trinucleotide
| Category | Allele 161 | Allele 176 | Allele 192 |
|---|---|---|---|
| C0 | 0.117 | 0.032 | 0.074 |
| C7 | 0.074 | 0.037 | 0.019 |
| C9 | 0.125 | 0.067 | 0.048 |
| C10 | 0.143 | 0.045 | 0.036 |
| C12 | 0.12 | 0.1 | 0.02 |
#### Chart: L039 Trinucleotide
| Category | Allele 123 | Allele 124 | Allele 132 | Allele 135 | Allele 138 | Allele 140 | Allele 143 |
|---|---|---|---|---|---|---|---|
| C0 | 0.096 | 0.149 | 0.138 | 0.128 | 0.043 | 0.223 | 0.064 |
| C7 | 0.074 | 0.13 | 0.259 | 0.019 | 0.111 | 0.222 | 0.074 |
| C9 | 0.067 | 0.154 | 0.125 | 0.067 | 0.048 | 0.231 | 0.096 |
| C10 | 0.063 | 0.107 | 0.232 | 0.071 | 0.098 | 0.223 | 0.063 |
| C12 | 0.04 | 0.12 | 0.24 | 0.12 | 0.06 | 0.14 | 0.1 |
#### Chart: L040 Tetranucleotide
| Category | Allele 108 | Allele 111 | Allele 112 | Allele 113 |
|---|---|---|---|---|
| C0 | 0.085 | 0.287 | 0.543 | 0.043 |
| C7 | 0.093 | 0.37 | 0.463 | 0.037 |
| C9 | 0.106 | 0.337 | 0.385 | 0.058 |
| C10 | 0.134 | 0.33 | 0.464 | 0.045 |
| C12 | 0.0 | 0.3 | 0.52 | 0.08 |
#### Chart: L041 Tetranucleotide
| Category | Allele 110 | Allele 118 | Allele 144 | Allele 147 | Allele 149 | Allele 150 |
|---|---|---|---|---|---|---|
| C0 | 0.181 | 0.074 | 0.043 | 0.138 | 0.17 | 0.128 |
| C7 | 0.093 | 0.056 | 0.167 | 0.167 | 0.111 | 0.093 |
| C9 | 0.125 | 0.048 | 0.173 | 0.173 | 0.144 | 0.106 |
| C10 | 0.134 | 0.063 | 0.179 | 0.143 | 0.107 | 0.063 |
| C12 | 0.208 | 0.021 | 0.25 | 0.083 | 0.083 | 0.083 |
#### Chart: L048 Trinucleotide
| Category | Allele 102 | Allele 115 | Allele 117 | Allele 119 |
|---|---|---|---|---|
| C0 | 0.064 | 0.043 | 0.436 | 0.319 |
| C7 | 0.093 | 0.111 | 0.333 | 0.222 |
| C9 | 0.038 | 0.106 | 0.298 | 0.327 |
| C10 | 0.071 | 0.08 | 0.429 | 0.259 |
| C12 | 0.04 | 0.04 | 0.32 | 0.34 |
#### Chart: L049 Trinucleotide
| Category | Allele 110 | Allele 124 | Allele 127 | Allele 131 | Allele 134 |
|---|---|---|---|---|---|
| C0 | 0.064 | 0.202 | 0.319 | 0.17 | 0.064 |
| C7 | 0.019 | 0.093 | 0.278 | 0.241 | 0.037 |
| C9 | 0.067 | 0.106 | 0.375 | 0.154 | 0.096 |
| C10 | 0.045 | 0.098 | 0.313 | 0.241 | 0.08 |
| C12 | 0.06 | 0.1 | 0.32 | 0.26 | 0.08 |

### Slide 13
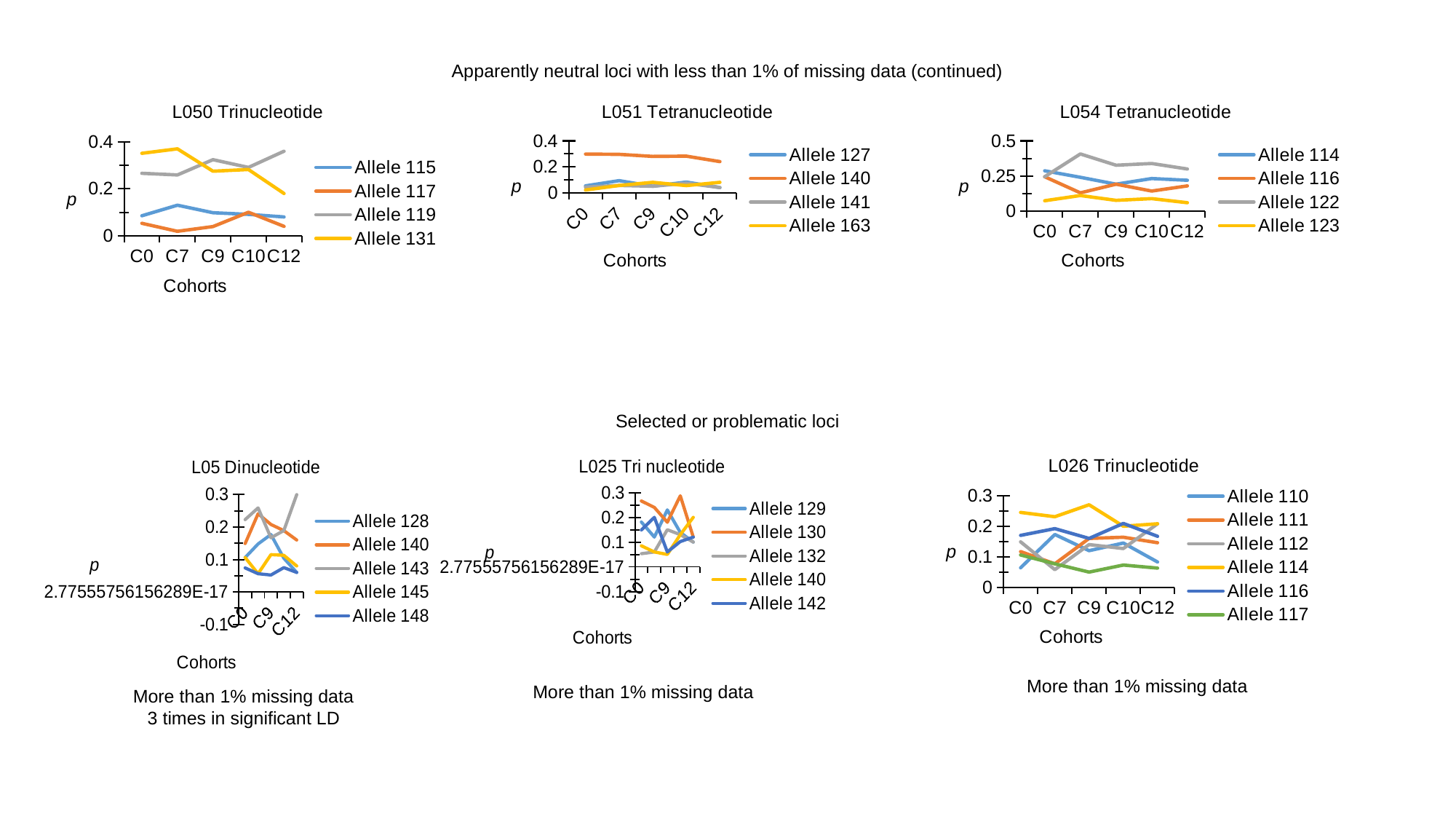

Apparently neutral loci with less than 1% of missing data (continued)
#### Chart: L050 Trinucleotide
| Category | Allele 115 | Allele 117 | Allele 119 | Allele 131 |
|---|---|---|---|---|
| C0 | 0.085 | 0.053 | 0.266 | 0.351 |
| C7 | 0.13 | 0.019 | 0.259 | 0.37 |
| C9 | 0.098 | 0.039 | 0.324 | 0.275 |
| C10 | 0.091 | 0.1 | 0.291 | 0.282 |
| C12 | 0.08 | 0.04 | 0.36 | 0.18 |
#### Chart: L051 Tetranucleotide
| Category | Allele 127 | Allele 140 | Allele 141 | Allele 163 |
|---|---|---|---|---|
| C0 | 0.053 | 0.298 | 0.043 | 0.021 |
| C7 | 0.093 | 0.296 | 0.056 | 0.056 |
| C9 | 0.05 | 0.28 | 0.05 | 0.08 |
| C10 | 0.082 | 0.282 | 0.073 | 0.055 |
| C12 | 0.04 | 0.24 | 0.04 | 0.08 |
#### Chart: L054 Tetranucleotide
| Category | Allele 114 | Allele 116 | Allele 122 | Allele 123 |
|---|---|---|---|---|
| C0 | 0.287 | 0.245 | 0.245 | 0.074 |
| C7 | 0.241 | 0.13 | 0.407 | 0.111 |
| C9 | 0.192 | 0.192 | 0.327 | 0.077 |
| C10 | 0.232 | 0.143 | 0.339 | 0.089 |
| C12 | 0.22 | 0.18 | 0.3 | 0.06 |Selected or problematic loci
#### Chart: L026 Trinucleotide
| Category | Allele 110 | Allele 111 | Allele 112 | Allele 114 | Allele 116 | Allele 117 |
|---|---|---|---|---|---|---|
| C0 | 0.064 | 0.117 | 0.149 | 0.245 | 0.17 | 0.106 |
| C7 | 0.173 | 0.077 | 0.058 | 0.231 | 0.192 | 0.077 |
| C9 | 0.12 | 0.16 | 0.14 | 0.27 | 0.16 | 0.05 |
| C10 | 0.145 | 0.164 | 0.127 | 0.2 | 0.209 | 0.073 |
| C12 | 0.083 | 0.146 | 0.208 | 0.208 | 0.167 | 0.063 |
#### Chart: L05 Dinucleotide
| Category | Allele 128 | Allele 140 | Allele 143 | Allele 145 | Allele 148 |
|---|---|---|---|---|---|
| C0 | 0.106 | 0.149 | 0.223 | 0.106 | 0.074 |
| C7 | 0.148 | 0.241 | 0.259 | 0.056 | 0.056 |
| C9 | 0.177 | 0.208 | 0.167 | 0.115 | 0.052 |
| C10 | 0.104 | 0.189 | 0.189 | 0.113 | 0.075 |
| C12 | 0.06 | 0.16 | 0.3 | 0.08 | 0.06 |
#### Chart: L025 Tri nucleotide
| Category | Allele 129 | Allele 130 | Allele 132 | Allele 140 | Allele 142 |
|---|---|---|---|---|---|
| C0 | 0.181 | 0.266 | 0.053 | 0.085 | 0.149 |
| C7 | 0.12 | 0.24 | 0.06 | 0.06 | 0.2 |
| C9 | 0.23 | 0.18 | 0.15 | 0.05 | 0.06 |
| C10 | 0.139 | 0.287 | 0.13 | 0.13 | 0.102 |
| C12 | 0.1 | 0.12 | 0.1 | 0.2 | 0.12 |More than 1% missing data
More than 1% missing data
More than 1% missing data
3 times in significant LD

### Slide 14
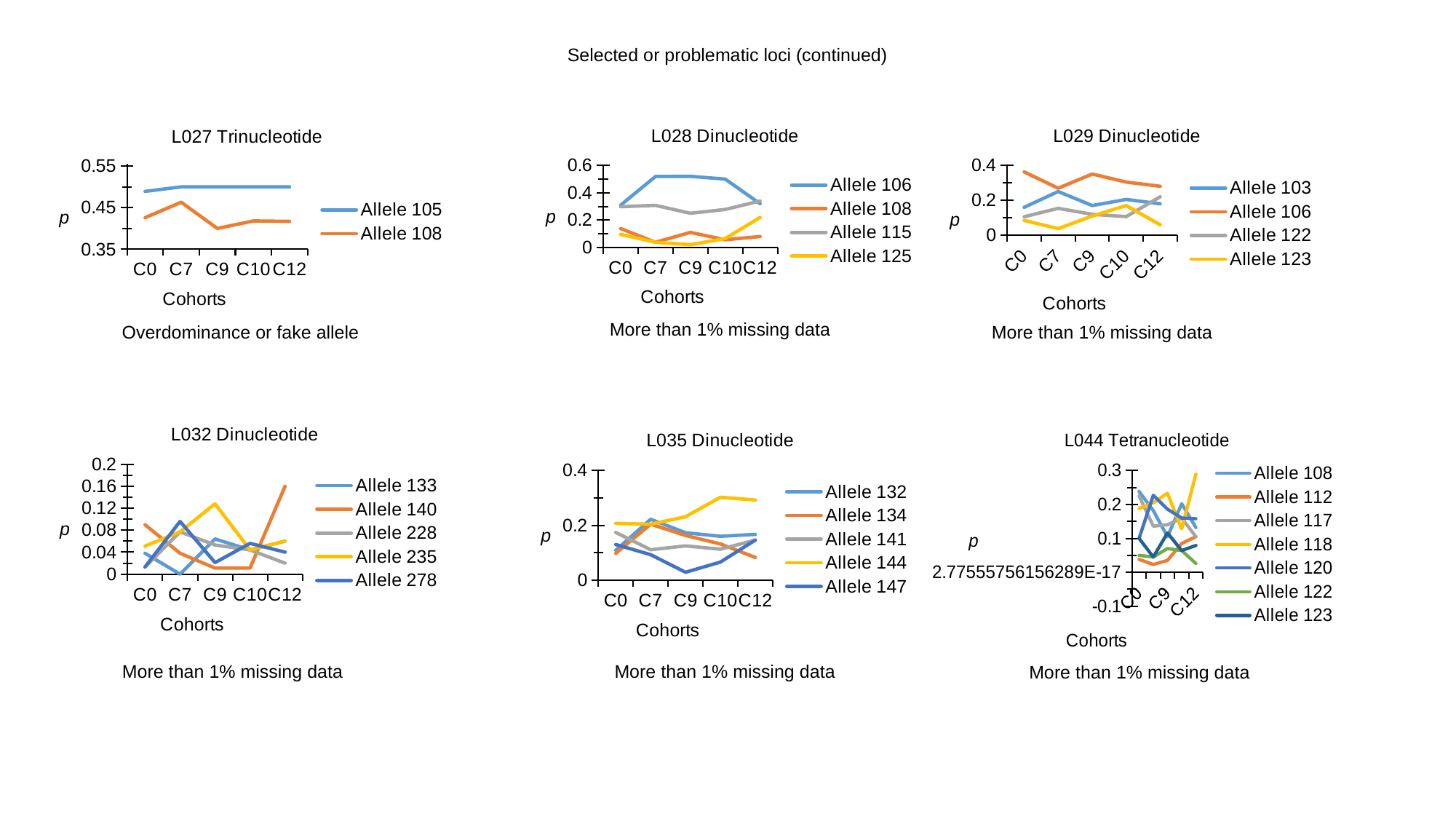

Selected or problematic loci (continued)
#### Chart: L029 Dinucleotide
| Category | Allele 103 | Allele 106 | Allele 122 | Allele 123 |
|---|---|---|---|---|
| C0 | 0.16 | 0.362 | 0.106 | 0.085 |
| C7 | 0.25 | 0.269 | 0.154 | 0.038 |
| C9 | 0.17 | 0.35 | 0.12 | 0.11 |
| C10 | 0.205 | 0.304 | 0.107 | 0.17 |
| C12 | 0.18 | 0.28 | 0.22 | 0.06 |
#### Chart: L028 Dinucleotide
| Category | Allele 106 | Allele 108 | Allele 115 | Allele 125 |
|---|---|---|---|---|
| C0 | 0.309 | 0.138 | 0.298 | 0.096 |
| C7 | 0.519 | 0.038 | 0.308 | 0.038 |
| C9 | 0.52 | 0.11 | 0.25 | 0.02 |
| C10 | 0.5 | 0.056 | 0.278 | 0.065 |
| C12 | 0.32 | 0.08 | 0.34 | 0.22 |
#### Chart: L027 Trinucleotide
| Category | Allele 105 | Allele 108 |
|---|---|---|
| C0 | 0.489 | 0.426 |
| C7 | 0.5 | 0.463 |
| C9 | 0.5 | 0.4 |
| C10 | 0.5 | 0.418 |
| C12 | 0.5 | 0.417 |More than 1% missing data
Overdominance or fake allele
More than 1% missing data
#### Chart: L032 Dinucleotide
| Category | Allele 133 | Allele 140 | Allele 228 | Allele 235 | Allele 278 |
|---|---|---|---|---|---|
| C0 | 0.038 | 0.09 | 0.013 | 0.051 | 0.013 |
| C7 | 0.0 | 0.038 | 0.077 | 0.077 | 0.096 |
| C9 | 0.064 | 0.011 | 0.053 | 0.128 | 0.021 |
| C10 | 0.044 | 0.011 | 0.044 | 0.044 | 0.056 |
| C12 | 0.06 | 0.16 | 0.02 | 0.06 | 0.04 |
#### Chart: L035 Dinucleotide
| Category | Allele 132 | Allele 134 | Allele 141 | Allele 144 | Allele 147 |
|---|---|---|---|---|---|
| C0 | 0.109 | 0.098 | 0.174 | 0.207 | 0.13 |
| C7 | 0.222 | 0.204 | 0.111 | 0.204 | 0.093 |
| C9 | 0.173 | 0.163 | 0.125 | 0.231 | 0.029 |
| C10 | 0.16 | 0.132 | 0.113 | 0.302 | 0.066 |
| C12 | 0.167 | 0.083 | 0.146 | 0.292 | 0.146 |
#### Chart: L044 Tetranucleotide
| Category | Allele 108 | Allele 112 | Allele 117 | Allele 118 | Allele 120 | Allele 122 | Allele 123 |
|---|---|---|---|---|---|---|---|
| C0 | 0.238 | 0.038 | 0.225 | 0.188 | 0.1 | 0.05 | 0.1 |
| C7 | 0.182 | 0.023 | 0.136 | 0.205 | 0.227 | 0.045 | 0.045 |
| C9 | 0.105 | 0.035 | 0.14 | 0.233 | 0.186 | 0.07 | 0.116 |
| C10 | 0.202 | 0.085 | 0.16 | 0.128 | 0.16 | 0.064 | 0.064 |
| C12 | 0.132 | 0.105 | 0.105 | 0.289 | 0.158 | 0.026 | 0.079 |More than 1% missing data
More than 1% missing data
More than 1% missing data

### Slide 15
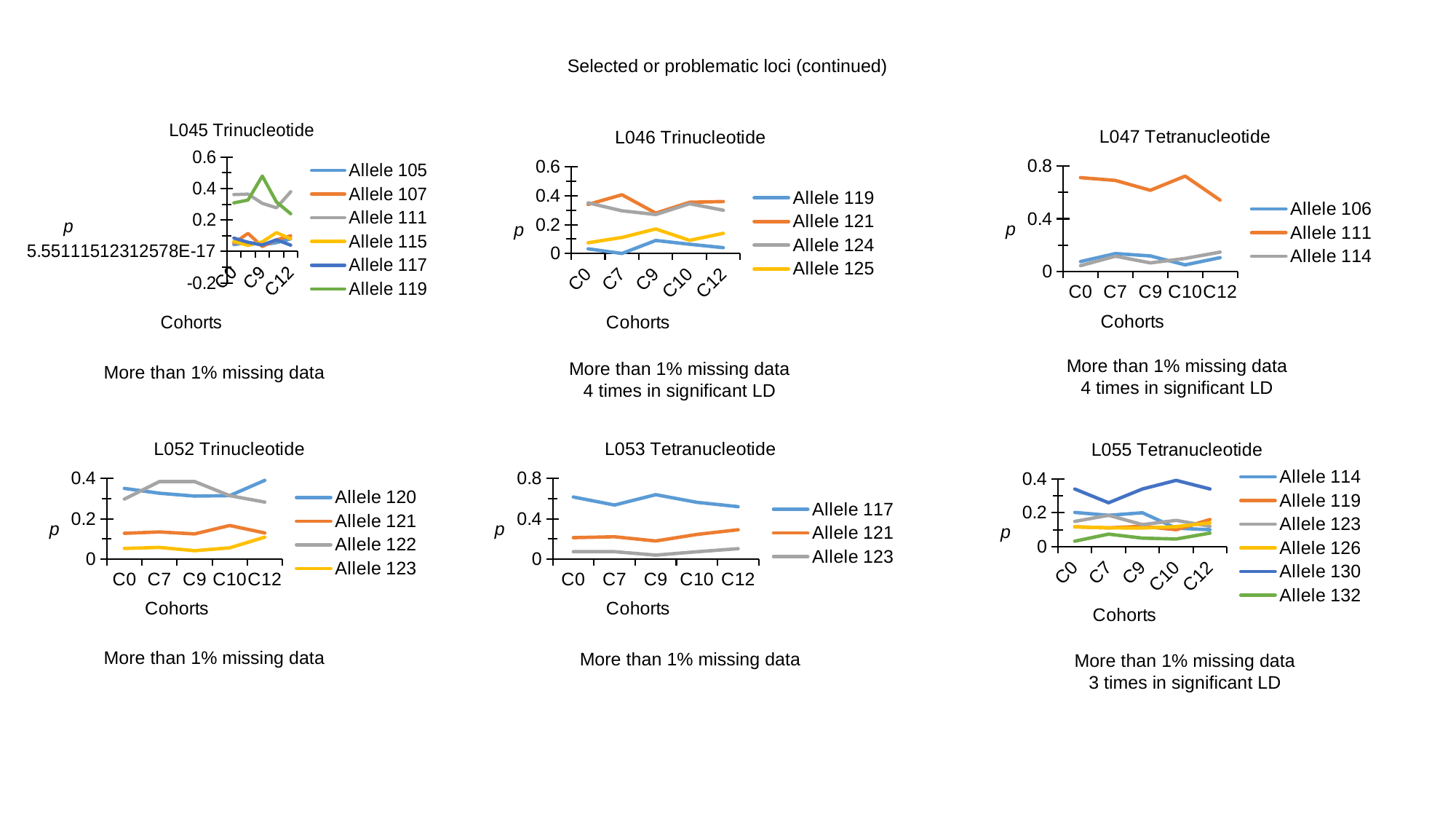

Selected or problematic loci (continued)
#### Chart: L045 Trinucleotide
| Category | Allele 105 | Allele 107 | Allele 111 | Allele 115 | Allele 117 | Allele 119 |
|---|---|---|---|---|---|---|
| C0 | 0.043 | 0.053 | 0.362 | 0.064 | 0.085 | 0.309 |
| C7 | 0.058 | 0.115 | 0.365 | 0.038 | 0.058 | 0.327 |
| C9 | 0.041 | 0.031 | 0.306 | 0.061 | 0.041 | 0.48 |
| C10 | 0.056 | 0.074 | 0.278 | 0.12 | 0.074 | 0.315 |
| C12 | 0.08 | 0.1 | 0.38 | 0.08 | 0.04 | 0.24 |
#### Chart: L047 Tetranucleotide
| Category | Allele 106 | Allele 111 | Allele 114 |
|---|---|---|---|
| C0 | 0.074 | 0.713 | 0.043 |
| C7 | 0.135 | 0.692 | 0.115 |
| C9 | 0.117 | 0.617 | 0.064 |
| C10 | 0.049 | 0.725 | 0.098 |
| C12 | 0.104 | 0.542 | 0.146 |
#### Chart: L046 Trinucleotide
| Category | Allele 119 | Allele 121 | Allele 124 | Allele 125 |
|---|---|---|---|---|
| C0 | 0.032 | 0.34 | 0.351 | 0.074 |
| C7 | 0.0 | 0.407 | 0.296 | 0.111 |
| C9 | 0.09 | 0.28 | 0.27 | 0.17 |
| C10 | 0.064 | 0.355 | 0.345 | 0.091 |
| C12 | 0.04 | 0.36 | 0.3 | 0.14 |More than 1% missing data
4 times in significant LD
More than 1% missing data
4 times in significant LD
More than 1% missing data
#### Chart: L052 Trinucleotide
| Category | Allele 120 | Allele 121 | Allele 122 | Allele 123 |
|---|---|---|---|---|
| C0 | 0.351 | 0.128 | 0.298 | 0.053 |
| C7 | 0.327 | 0.135 | 0.385 | 0.058 |
| C9 | 0.313 | 0.125 | 0.385 | 0.042 |
| C10 | 0.315 | 0.167 | 0.315 | 0.056 |
| C12 | 0.391 | 0.13 | 0.283 | 0.109 |
#### Chart: L053 Tetranucleotide
| Category | Allele 117 | Allele 121 | Allele 123 |
|---|---|---|---|
| C0 | 0.617 | 0.213 | 0.074 |
| C7 | 0.537 | 0.222 | 0.074 |
| C9 | 0.64 | 0.18 | 0.04 |
| C10 | 0.564 | 0.245 | 0.073 |
| C12 | 0.521 | 0.292 | 0.104 |
#### Chart: L055 Tetranucleotide
| Category | Allele 114 | Allele 119 | Allele 123 | Allele 126 | Allele 130 | Allele 132 |
|---|---|---|---|---|---|---|
| C0 | 0.202 | 0.117 | 0.149 | 0.117 | 0.34 | 0.032 |
| C7 | 0.185 | 0.111 | 0.185 | 0.111 | 0.259 | 0.074 |
| C9 | 0.2 | 0.12 | 0.13 | 0.11 | 0.34 | 0.05 |
| C10 | 0.109 | 0.1 | 0.155 | 0.118 | 0.391 | 0.045 |
| C12 | 0.1 | 0.16 | 0.12 | 0.14 | 0.34 | 0.08 |More than 1% missing data
More than 1% missing data
More than 1% missing data
3 times in significant LD

### Slide 16
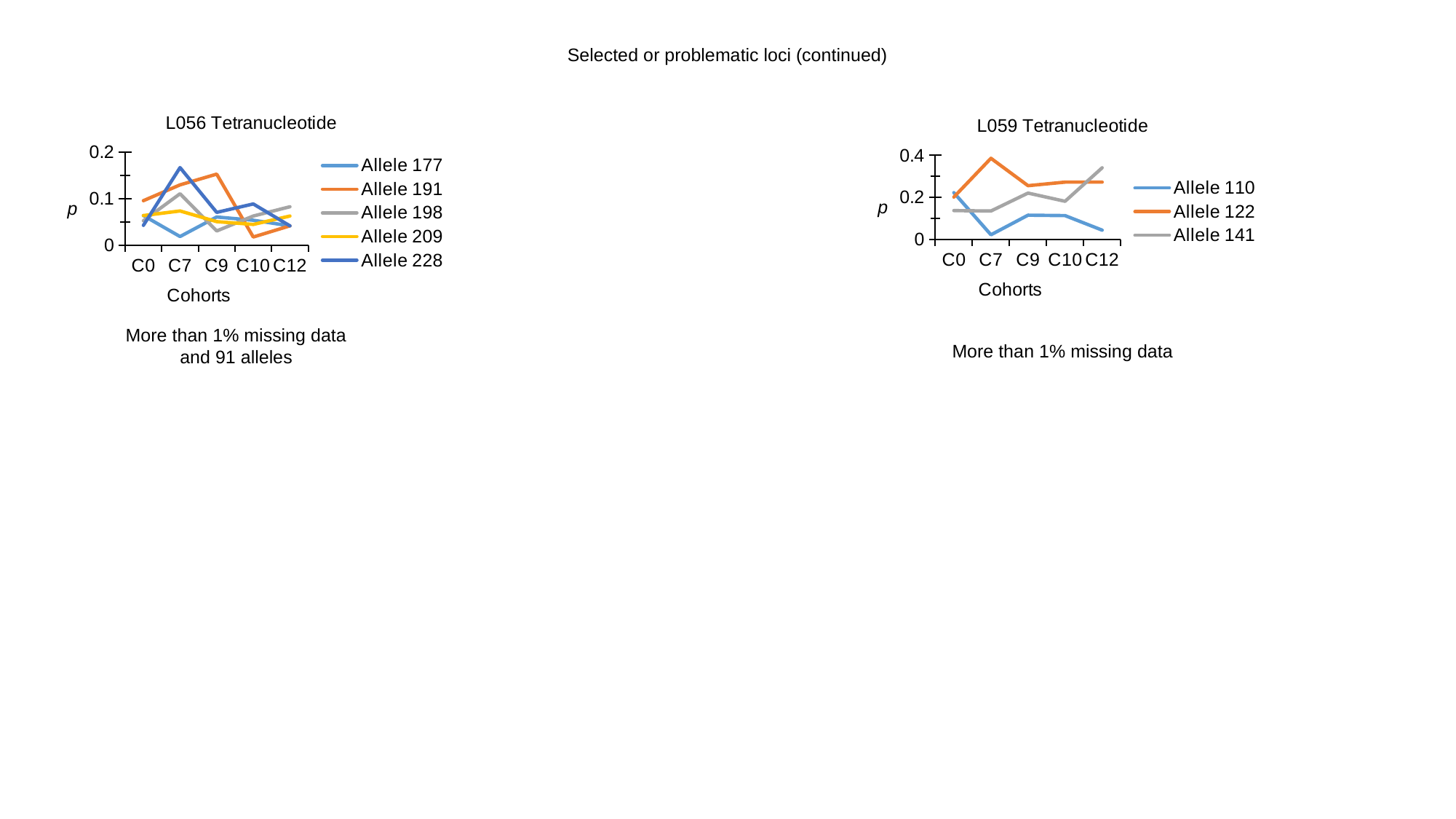

Selected or problematic loci (continued)
#### Chart: L056 Tetranucleotide
| Category | Allele 177 | Allele 191 | Allele 198 | Allele 209 | Allele 228 |
|---|---|---|---|---|---|
| C0 | 0.064 | 0.096 | 0.053 | 0.064 | 0.043 |
| C7 | 0.019 | 0.13 | 0.111 | 0.074 | 0.167 |
| C9 | 0.061 | 0.153 | 0.031 | 0.051 | 0.071 |
| C10 | 0.054 | 0.018 | 0.063 | 0.045 | 0.089 |
| C12 | 0.042 | 0.042 | 0.083 | 0.063 | 0.042 |
#### Chart: L059 Tetranucleotide
| Category | Allele 110 | Allele 122 | Allele 141 |
|---|---|---|---|
| C0 | 0.223 | 0.202 | 0.138 |
| C7 | 0.023 | 0.386 | 0.136 |
| C9 | 0.116 | 0.256 | 0.221 |
| C10 | 0.114 | 0.273 | 0.182 |
| C12 | 0.045 | 0.273 | 0.341 |More than 1% missing data
and 91 alleles
More than 1% missing data

### Slide 17
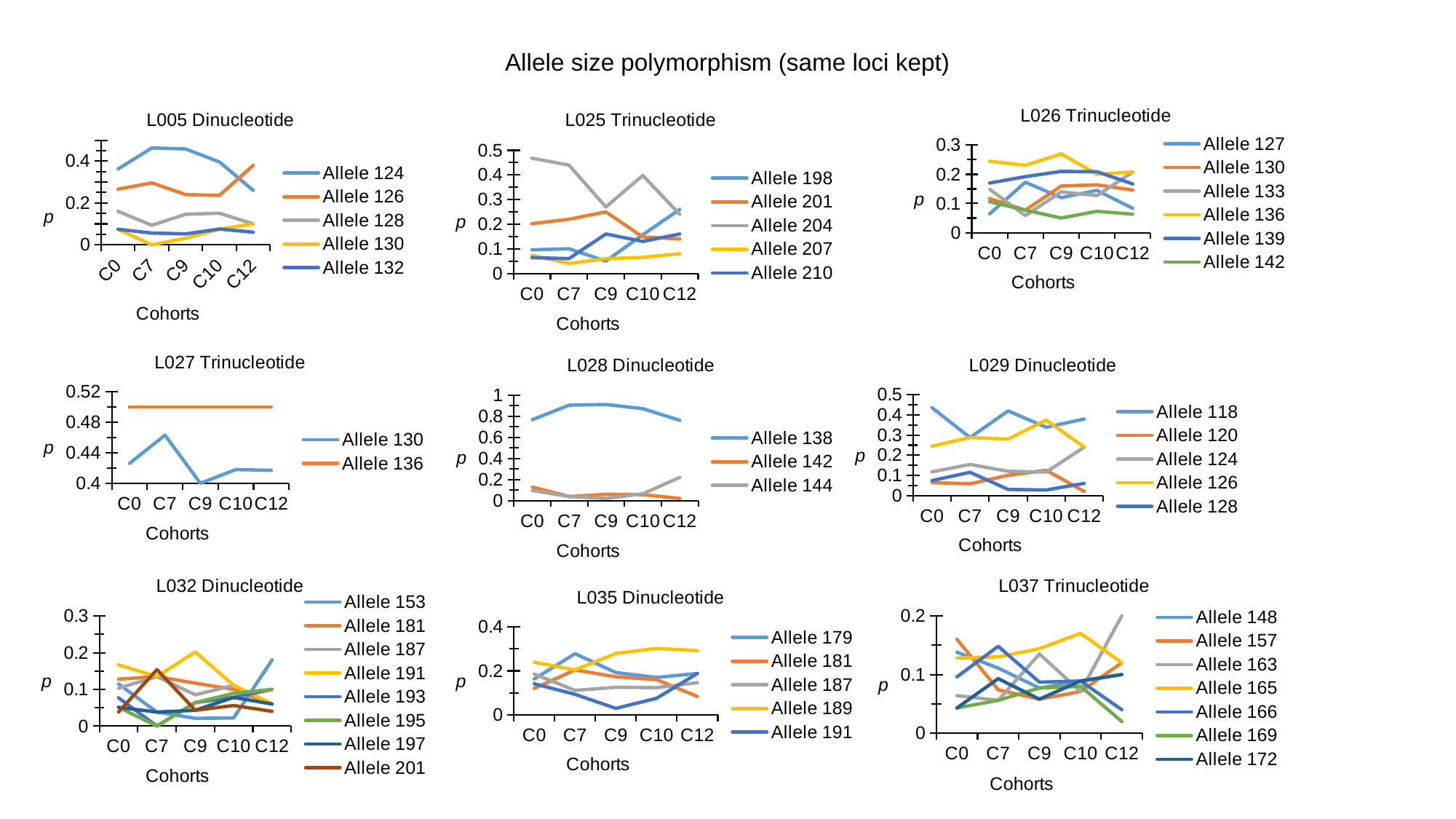

Allele size polymorphism (same loci kept)
#### Chart: L026 Trinucleotide
| Category | Allele 127 | Allele 130 | Allele 133 | Allele 136 | Allele 139 | Allele 142 |
|---|---|---|---|---|---|---|
| C0 | 0.064 | 0.117 | 0.149 | 0.245 | 0.17 | 0.106 |
| C7 | 0.173 | 0.077 | 0.058 | 0.231 | 0.192 | 0.077 |
| C9 | 0.12 | 0.16 | 0.14 | 0.27 | 0.21 | 0.05 |
| C10 | 0.145 | 0.164 | 0.127 | 0.2 | 0.209 | 0.073 |
| C12 | 0.083 | 0.146 | 0.208 | 0.208 | 0.167 | 0.063 |
#### Chart: L005 Dinucleotide
| Category | Allele 124 | Allele 126 | Allele 128 | Allele 130 | Allele 132 |
|---|---|---|---|---|---|
| C0 | 0.362 | 0.266 | 0.16 | 0.074 | 0.074 |
| C7 | 0.463 | 0.296 | 0.093 | 0.0 | 0.056 |
| C9 | 0.458 | 0.24 | 0.146 | 0.031 | 0.052 |
| C10 | 0.396 | 0.236 | 0.151 | 0.075 | 0.075 |
| C12 | 0.26 | 0.38 | 0.1 | 0.1 | 0.06 |
#### Chart: L025 Trinucleotide
| Category | Allele 198 | Allele 201 | Allele 204 | Allele 207 | Allele 210 |
|---|---|---|---|---|---|
| C0 | 0.096 | 0.202 | 0.468 | 0.074 | 0.064 |
| C7 | 0.1 | 0.22 | 0.44 | 0.04 | 0.06 |
| C9 | 0.05 | 0.25 | 0.27 | 0.06 | 0.16 |
| C10 | 0.157 | 0.148 | 0.398 | 0.065 | 0.13 |
| C12 | 0.26 | 0.14 | 0.24 | 0.08 | 0.16 |
#### Chart: L027 Trinucleotide
| Category | Allele 130 | Allele 136 |
|---|---|---|
| C0 | 0.426 | 0.5 |
| C7 | 0.463 | 0.5 |
| C9 | 0.4 | 0.5 |
| C10 | 0.418 | 0.5 |
| C12 | 0.417 | 0.5 |
#### Chart: L028 Dinucleotide
| Category | Allele 138 | Allele 142 | Allele 144 |
|---|---|---|---|
| C0 | 0.766 | 0.128 | 0.096 |
| C7 | 0.904 | 0.038 | 0.038 |
| C9 | 0.91 | 0.06 | 0.02 |
| C10 | 0.87 | 0.056 | 0.065 |
| C12 | 0.76 | 0.02 | 0.22 |
#### Chart: L029 Dinucleotide
| Category | Allele 118 | Allele 120 | Allele 124 | Allele 126 | Allele 128 |
|---|---|---|---|---|---|
| C0 | 0.436 | 0.064 | 0.117 | 0.245 | 0.074 |
| C7 | 0.288 | 0.058 | 0.154 | 0.288 | 0.115 |
| C9 | 0.42 | 0.1 | 0.12 | 0.28 | 0.03 |
| C10 | 0.339 | 0.125 | 0.116 | 0.375 | 0.027 |
| C12 | 0.38 | 0.02 | 0.24 | 0.24 | 0.06 |
#### Chart: L037 Trinucleotide
| Category | Allele 148 | Allele 157 | Allele 163 | Allele 165 | Allele 166 | Allele 169 | Allele 172 |
|---|---|---|---|---|---|---|---|
| C0 | 0.138 | 0.16 | 0.064 | 0.128 | 0.096 | 0.043 | 0.043 |
| C7 | 0.111 | 0.074 | 0.056 | 0.13 | 0.148 | 0.056 | 0.093 |
| C9 | 0.077 | 0.058 | 0.135 | 0.144 | 0.087 | 0.077 | 0.058 |
| C10 | 0.089 | 0.071 | 0.071 | 0.17 | 0.089 | 0.08 | 0.089 |
| C12 | 0.1 | 0.12 | 0.2 | 0.12 | 0.04 | 0.02 | 0.1 |
#### Chart: L032 Dinucleotide
| Category | Allele 153 | Allele 181 | Allele 187 | Allele 191 | Allele 193 | Allele 195 | Allele 197 | Allele 201 |
|---|---|---|---|---|---|---|---|---|
| C0 | 0.115 | 0.128 | 0.103 | 0.167 | 0.077 | 0.051 | 0.051 | 0.038 |
| C7 | 0.038 | 0.135 | 0.135 | 0.135 | 0.0 | 0.0 | 0.038 | 0.154 |
| C9 | 0.021 | 0.117 | 0.085 | 0.202 | 0.064 | 0.064 | 0.043 | 0.043 |
| C10 | 0.022 | 0.1 | 0.111 | 0.111 | 0.078 | 0.089 | 0.078 | 0.056 |
| C12 | 0.18 | 0.06 | 0.06 | 0.06 | 0.1 | 0.1 | 0.06 | 0.04 |
#### Chart: L035 Dinucleotide
| Category | Allele 179 | Allele 181 | Allele 187 | Allele 189 | Allele 191 |
|---|---|---|---|---|---|
| C0 | 0.163 | 0.12 | 0.185 | 0.239 | 0.141 |
| C7 | 0.278 | 0.204 | 0.111 | 0.204 | 0.093 |
| C9 | 0.192 | 0.173 | 0.125 | 0.279 | 0.029 |
| C10 | 0.17 | 0.16 | 0.123 | 0.302 | 0.075 |
| C12 | 0.188 | 0.083 | 0.146 | 0.292 | 0.188 |

### Slide 18
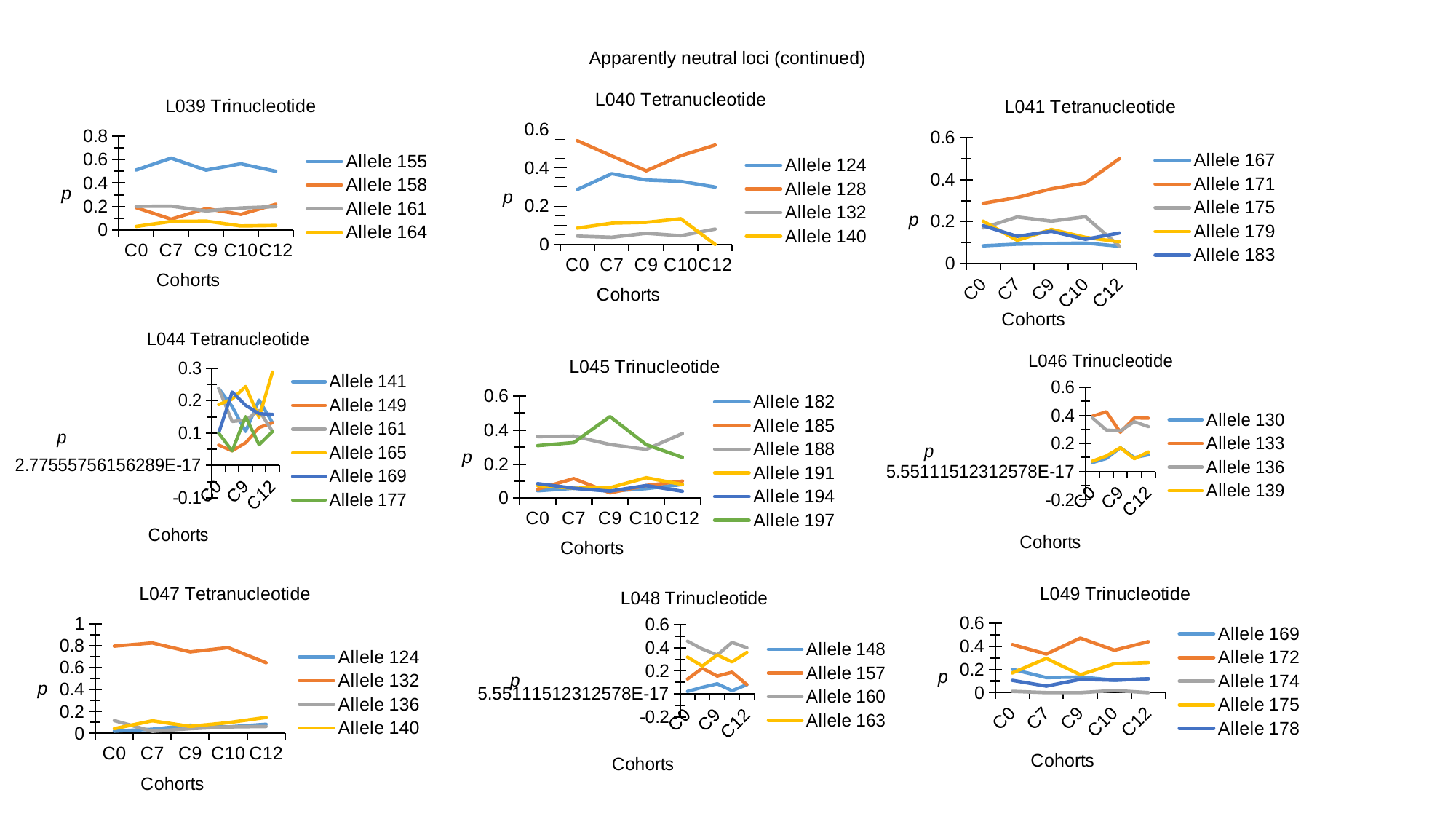

Apparently neutral loci (continued)
#### Chart: L040 Tetranucleotide
| Category | Allele 124 | Allele 128 | Allele 132 | Allele 140 |
|---|---|---|---|---|
| C0 | 0.287 | 0.543 | 0.043 | 0.085 |
| C7 | 0.37 | 0.463 | 0.037 | 0.111 |
| C9 | 0.337 | 0.385 | 0.058 | 0.115 |
| C10 | 0.33 | 0.464 | 0.045 | 0.134 |
| C12 | 0.3 | 0.52 | 0.08 | 0.0 |
#### Chart: L039 Trinucleotide
| Category | Allele 155 | Allele 158 | Allele 161 | Allele 164 |
|---|---|---|---|---|
| C0 | 0.511 | 0.191 | 0.202 | 0.032 |
| C7 | 0.611 | 0.093 | 0.204 | 0.074 |
| C9 | 0.51 | 0.183 | 0.163 | 0.077 |
| C10 | 0.563 | 0.134 | 0.188 | 0.036 |
| C12 | 0.5 | 0.22 | 0.2 | 0.04 |
#### Chart: L041 Tetranucleotide
| Category | Allele 167 | Allele 171 | Allele 175 | Allele 179 | Allele 183 |
|---|---|---|---|---|---|
| C0 | 0.085 | 0.287 | 0.17 | 0.202 | 0.181 |
| C7 | 0.093 | 0.315 | 0.222 | 0.111 | 0.13 |
| C9 | 0.096 | 0.356 | 0.202 | 0.163 | 0.154 |
| C10 | 0.098 | 0.384 | 0.223 | 0.125 | 0.116 |
| C12 | 0.083 | 0.5 | 0.083 | 0.104 | 0.146 |
#### Chart: L044 Tetranucleotide
| Category | Allele 141 | Allele 149 | Allele 161 | Allele 165 | Allele 169 | Allele 177 |
|---|---|---|---|---|---|---|
| C0 | 0.238 | 0.063 | 0.238 | 0.188 | 0.1 | 0.1 |
| C7 | 0.182 | 0.045 | 0.136 | 0.205 | 0.227 | 0.045 |
| C9 | 0.105 | 0.07 | 0.14 | 0.244 | 0.186 | 0.151 |
| C10 | 0.202 | 0.117 | 0.17 | 0.149 | 0.16 | 0.064 |
| C12 | 0.132 | 0.132 | 0.105 | 0.289 | 0.158 | 0.105 |
#### Chart: L046 Trinucleotide
| Category | Allele 130 | Allele 133 | Allele 136 | Allele 139 |
|---|---|---|---|---|
| C0 | 0.064 | 0.394 | 0.383 | 0.074 |
| C7 | 0.093 | 0.426 | 0.296 | 0.111 |
| C9 | 0.17 | 0.28 | 0.29 | 0.17 |
| C10 | 0.1 | 0.382 | 0.355 | 0.091 |
| C12 | 0.12 | 0.38 | 0.32 | 0.14 |
#### Chart: L045 Trinucleotide
| Category | Allele 182 | Allele 185 | Allele 188 | Allele 191 | Allele 194 | Allele 197 |
|---|---|---|---|---|---|---|
| C0 | 0.043 | 0.053 | 0.362 | 0.074 | 0.085 | 0.309 |
| C7 | 0.058 | 0.115 | 0.365 | 0.058 | 0.058 | 0.327 |
| C9 | 0.041 | 0.031 | 0.316 | 0.061 | 0.041 | 0.48 |
| C10 | 0.056 | 0.074 | 0.287 | 0.12 | 0.074 | 0.315 |
| C12 | 0.08 | 0.1 | 0.38 | 0.08 | 0.04 | 0.24 |
#### Chart: L047 Tetranucleotide
| Category | Allele 124 | Allele 132 | Allele 136 | Allele 140 |
|---|---|---|---|---|
| C0 | 0.021 | 0.798 | 0.117 | 0.043 |
| C7 | 0.038 | 0.827 | 0.019 | 0.115 |
| C9 | 0.074 | 0.745 | 0.043 | 0.064 |
| C10 | 0.059 | 0.784 | 0.059 | 0.098 |
| C12 | 0.083 | 0.646 | 0.063 | 0.146 |
#### Chart: L049 Trinucleotide
| Category | Allele 169 | Allele 172 | Allele 174 | Allele 175 | Allele 178 |
|---|---|---|---|---|---|
| C0 | 0.202 | 0.415 | 0.011 | 0.17 | 0.106 |
| C7 | 0.13 | 0.333 | 0.0 | 0.296 | 0.056 |
| C9 | 0.135 | 0.471 | 0.0 | 0.154 | 0.115 |
| C10 | 0.107 | 0.366 | 0.018 | 0.25 | 0.107 |
| C12 | 0.12 | 0.44 | 0.0 | 0.26 | 0.12 |
#### Chart: L048 Trinucleotide
| Category | Allele 148 | Allele 157 | Allele 160 | Allele 163 |
|---|---|---|---|---|
| C0 | 0.021 | 0.128 | 0.457 | 0.319 |
| C7 | 0.056 | 0.222 | 0.389 | 0.241 |
| C9 | 0.087 | 0.154 | 0.337 | 0.337 |
| C10 | 0.027 | 0.188 | 0.446 | 0.277 |
| C12 | 0.08 | 0.08 | 0.4 | 0.36 |

### Slide 19
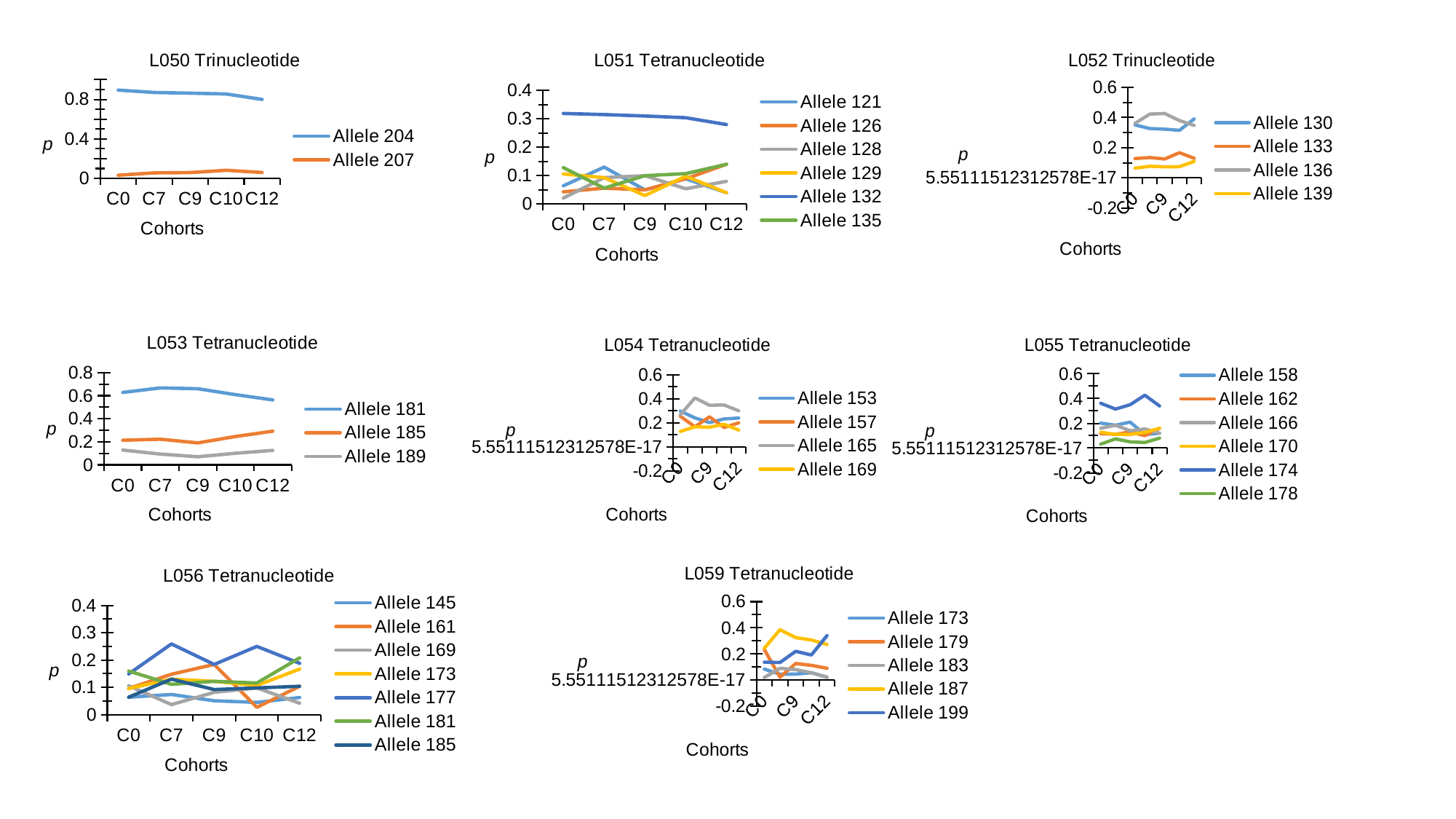

#### Chart: L050 Trinucleotide
| Category | Allele 204 | Allele 207 |
|---|---|---|
| C0 | 0.894 | 0.032 |
| C7 | 0.87 | 0.056 |
| C9 | 0.863 | 0.059 |
| C10 | 0.855 | 0.082 |
| C12 | 0.8 | 0.06 |
#### Chart: L051 Tetranucleotide
| Category | Allele 121 | Allele 126 | Allele 128 | Allele 129 | Allele 132 | Allele 135 |
|---|---|---|---|---|---|---|
| C0 | 0.064 | 0.043 | 0.021 | 0.106 | 0.319 | 0.128 |
| C7 | 0.13 | 0.056 | 0.093 | 0.093 | 0.315 | 0.056 |
| C9 | 0.05 | 0.05 | 0.1 | 0.03 | 0.31 | 0.1 |
| C10 | 0.089 | 0.089 | 0.054 | 0.098 | 0.304 | 0.107 |
| C12 | 0.04 | 0.14 | 0.08 | 0.04 | 0.28 | 0.14 |
#### Chart: L052 Trinucleotide
| Category | Allele 130 | Allele 133 | Allele 136 | Allele 139 |
|---|---|---|---|---|
| C0 | 0.351 | 0.128 | 0.362 | 0.064 |
| C7 | 0.327 | 0.135 | 0.423 | 0.077 |
| C9 | 0.323 | 0.125 | 0.427 | 0.073 |
| C10 | 0.315 | 0.167 | 0.38 | 0.074 |
| C12 | 0.391 | 0.13 | 0.348 | 0.109 |
#### Chart: L053 Tetranucleotide
| Category | Allele 181 | Allele 185 | Allele 189 |
|---|---|---|---|
| C0 | 0.628 | 0.213 | 0.128 |
| C7 | 0.667 | 0.222 | 0.093 |
| C9 | 0.66 | 0.19 | 0.07 |
| C10 | 0.609 | 0.245 | 0.1 |
| C12 | 0.563 | 0.292 | 0.125 |
#### Chart: L055 Tetranucleotide
| Category | Allele 158 | Allele 162 | Allele 166 | Allele 170 | Allele 174 | Allele 178 |
|---|---|---|---|---|---|---|
| C0 | 0.202 | 0.117 | 0.16 | 0.128 | 0.362 | 0.032 |
| C7 | 0.185 | 0.111 | 0.185 | 0.111 | 0.315 | 0.074 |
| C9 | 0.21 | 0.13 | 0.14 | 0.11 | 0.35 | 0.05 |
| C10 | 0.109 | 0.1 | 0.155 | 0.127 | 0.427 | 0.045 |
| C12 | 0.12 | 0.16 | 0.12 | 0.16 | 0.34 | 0.08 |
#### Chart: L054 Tetranucleotide
| Category | Allele 153 | Allele 157 | Allele 165 | Allele 169 |
|---|---|---|---|---|
| C0 | 0.298 | 0.255 | 0.266 | 0.128 |
| C7 | 0.241 | 0.167 | 0.407 | 0.167 |
| C9 | 0.202 | 0.25 | 0.346 | 0.163 |
| C10 | 0.232 | 0.161 | 0.348 | 0.188 |
| C12 | 0.24 | 0.2 | 0.3 | 0.14 |
#### Chart: L059 Tetranucleotide
| Category | Allele 173 | Allele 179 | Allele 183 | Allele 187 | Allele 199 |
|---|---|---|---|---|---|
| C0 | 0.085 | 0.234 | 0.021 | 0.245 | 0.138 |
| C7 | 0.045 | 0.023 | 0.091 | 0.386 | 0.136 |
| C9 | 0.047 | 0.128 | 0.081 | 0.326 | 0.221 |
| C10 | 0.057 | 0.114 | 0.057 | 0.307 | 0.193 |
| C12 | 0.023 | 0.091 | 0.023 | 0.273 | 0.341 |
#### Chart: L056 Tetranucleotide
| Category | Allele 145 | Allele 161 | Allele 169 | Allele 173 | Allele 177 | Allele 181 | Allele 185 |
|---|---|---|---|---|---|---|---|
| C0 | 0.064 | 0.096 | 0.106 | 0.096 | 0.149 | 0.16 | 0.064 |
| C7 | 0.074 | 0.148 | 0.037 | 0.13 | 0.259 | 0.111 | 0.13 |
| C9 | 0.051 | 0.184 | 0.082 | 0.122 | 0.184 | 0.122 | 0.092 |
| C10 | 0.045 | 0.027 | 0.098 | 0.107 | 0.25 | 0.116 | 0.098 |
| C12 | 0.063 | 0.104 | 0.042 | 0.167 | 0.188 | 0.208 | 0.104 |
